# Control of ribosomal protein synthesis by Microprocessor complex

**DOI:** 10.1101/2020.04.24.060236

**Authors:** Xuan Jiang, Amit Prabhakar, Stephanie M. Van der Voorn, Prajakta Ghatpande, Barbara Celona, Srivats Venkataramanan, Lorenzo Calviello, Chuwen Lin, Wanpeng Wang, Brian L. Black, Stephen N. Floor, Giorgio Lagna, Akiko Hata

## Abstract

Ribosome biogenesis in eukaryotes requires stoichiometric production and assembly of 80 ribosomal proteins (RPs) and 4 ribosomal RNAs, and its rate must be coordinated with cellular growth. The indispensable regulator of RP biosynthesis is the 5’-terminal oligopyrimidine (TOP) motif, spanning the transcription start site of all RP genes. Here we show that the Microprocessor complex, previously linked to the first step of processing microRNAs (miRNAs), coregulates RP expression by binding the TOP motif of nascent RP mRNAs and stimulating transcription elongation via resolution of DNA/RNA hybrids. Cell growth arrest triggers nuclear export and degradation of the Microprocessor protein Drosha by the E3 ubiquitin ligase Nedd4, accumulation of DNA/RNA hybrids at RP gene loci, decreased RP synthesis, and ribosome deficiency, hence synchronizing ribosome production with cell growth. Conditional deletion of *Drosha* in erythroid progenitors phenocopies human ribosomopathies, in which ribosomal insufficiency leads to anemia. Outlining a miRNA-independent role of the Microprocessor complex at the interphase between cell growth and ribosome biogenesis offers a new paradigm by which cells alter their protein biosynthetic capacity and cellular metabolism.

## Introduction

Regulation of protein synthesis is essential to cell growth, differentiation, and homeostasis. Its fulcrum is the ribosome, the protein synthesis apparatus composed of 80 RPs and 4 ribosomal RNAs (rRNAs) in eukaryotes. Since ribosomes are abundant, both the stoichiometric synthesis of RPs and rRNAs, and its coordination with cell growth rate, are crucial. Insufficient ribosomes and mutations in RPs underlie human diseases known as ribosomopathies, which include Diamond-Blackfan anemia (DBA), 5q-Myelodysplastic syndrome, and T cell acute lymphoblastic leukemia. In DBA patients, ribosome insufficiency impairs translation of Gata1—a transcription factor essential for erythropoiesis—and causes anemia(Ludwig et al., 2014). In eukaryotes, RPs are regulated at different steps of their synthesis, including RP gene (RPG) transcription, mRNA splicing and translation(Leppek et al., 2018; Simsek and Barna, 2017). In plants, RPGs are encoded by small gene families, each comprising 2-7 RPGs that share 65-100% amino acid sequence identity(Byrne, 2009). It is largely unknown how coordinated expression of RPGs from multigene families is achieved(Byrne, 2009). In animals, a unique regulatory element appears to have evolved to coregulate all 80 RPGs and other house-keeping genes involved in cell growth: the 5’-terminal <u>o</u>ligo<u>p</u>yrimidines tract (TOP), a stretch of 5-25 pyrimidines located at the transcription start site (TSS) and involved in transcriptional and translational regulation (Hamilton et al., 2006; Meyuhas and Kahan, 2015; Perina et al., 2011; Rojas et al., 2018). The TOP motif is absent in fungi (yeast) RPGs, but present at the TSS of all RPGs in animals, from the simplest Placozoa, to Sponges (Porifera), Jellyfish (Cnidaria), *Drosophila* (Arthropoda), and vertebrates, and thus appears to have retained an evolutionarily conserved role in the coregulation of RP biogenesis(Hamilton et al., 2006; Meyuhas and Kahan, 2015; Perina et al., 2011; Rojas et al., 2018). Different molecules—including DNA and RNA binding proteins, as well as miRNAs—have been linked to the TOP-dependent control of RP biosynthesis, but each of these molecules accounts for the control of a few RPs, or is not conserved among all animals. Thus, an elusive molecule that associates with the TOP motif of all RPGs and functions as a master regulator of RP biosynthesis in response to the change in cellular growth environment remains to be identified(Meyuhas and Kahan, 2015; Patursky-Polischuk et al., 2014).

During transcription, a three-stranded nucleic acid structure known as an R-loop spontaneously forms(García-Muse and Aguilera, 2019). It is composed of a DNA/RNA hybrid (a single-stranded template DNA (ssDNA) hybridized with a nascent mRNA) and an associated non-template ssDNA(García-Muse and Aguilera, 2019; Sanz and Chédin, 2019; Sanz et al., 2016). R-loops are critical modulator of gene expression and DNA repair(García-Muse and Aguilera, 2019; Sanz and Chédin, 2019; Sanz et al., 2016), and their extended persistence during transcription is inhibitory to gene expression(Gowrishankar et al., 2013; Nudler, 2012). The abundance of R-loops is determined by the balance between its formation and resolution of DNA/RNA hybrids and various factors, including transcription factors, helicases, ribonucleases, topoisomerases, chromatin remodelers, proteins in DNA repair and RNA surveillance, have been identified to control R-loop homeostasis(García-Muse and Aguilera, 2019). Deregulation of R-loops, which results in aberrant gene expression and chromatin structure, increased DNA breaks, and genome instability, contributes to human disease, including neurological disorders and tumorigenesis(García-Muse and Aguilera, 2019; Groh and Gromak, 2014; Sanz et al., 2016).

The Microprocessor complex comprises two core components, the ribonuclease (RNase) III Drosha and its cofactor Dgcr8. It is essential for the biogenesis of most microRNAs (miRNAs), short RNAs that repress gene expression by binding to messenger RNAs and targeting them for degradation and/or preventing their translation(Han et al., 2004; Siomi and Siomi, 2010). The Microprocessor localizes predominantly in the nucleus, where it recognizes a hairpin structure in the primary-miRNA (pri-miRNA) transcript and cleaves the 5′ and 3′ flanking single-stranded RNA (ssRNA) to generate the stem-loop precursor-miRNA (pre-miRNA). The pre-miRNA is the substrate for Dicer processing in the cytoplasm. The processing of pri-miRNA by the Microprocessor is regulated by multiple accessory proteins that are components of the larger Microprocessor complex, including the DEAD-box RNA helicase 5 (Ddx5) and Ddx17(Hata and Lieberman, 2015). In addition to the genesis of miRNAs, recent studies uncovered various miRNA-independent functions of the Microprocessor, including (i) cleavage of a hairpin structure in selected mRNAs and their destabilization, (ii) processing of ribosomal RNAs, (iii) regulation of RNA polymerase II-mediated transcription, (iv) maintenance of genome integrity by facilitating DNA damage response, and (v) antiviral defense by cleavage of viral RNAs(Lee and Shin, 2018). While the RNase activity of the Microprocessor is well documented, a role for the RNA helicases in the processing of pri-miRNAs or in the miRNA-independent functions remains unknown.

Here we report that the Microprocessor complex associates with the 5’TOP motif of nascent RP transcripts, removes R-loops, and facilitates transcription elongation. Neither the ribonuclease activity of the Microprocessor nor miRNA biogenesis is required for this process, while the helicase activity of Ddx5 is necessary. While depletion of nutrients reduces Drosha protein and inhibits RP synthesis, exogenous expression of Drosha prevents RP synthesis inhibition by nutrients depletion. Evidence of the physiological significance of this new function of the Microprocessor complex was provided by the *Drosha* gene knockout in mouse, which results in impaired erythropoiesis and anemia due to reduced *Gata1* translation by insufficient ribosomes, a mechanism consistent with human ribosomopathies.

## Results

### Endothelial-specific deletion of Drosha in mouse impairs erythropoiesis

Mice with an endothelial-specific deletion of *Drosha* (hereafter referred to as *Drosha*-cKO mice) die around embryonic day (E)14.5-15.5 due to an erythropoiesis defect (Jiang et al., 2017). Although the number of erythroid progenitor cells (EPC) was similar in *Drosha*-cKO and control mice (littermates with at least one wild type *Drosha* allele, hereafter referred to as Ctrl) in the yolk sac (YS) at E9.5 **(Fig. 1a**), their ability to differentiate into mature erythroid cells [(in a methylcellulose colony-forming unit (CFU) assay, (**Fig. 1b**)] was severely affected by deletion of *Drosha*. To measure the effect of *Drosha* on erythroid differentiation in vivo, we separated EPC from the peripheral blood of E10.5 *Drosha*-cKO vs. Ctrl embryos based on their differentiation stage: more mature erythroid precursors (MEP, CD71^high^Ter119^high^) vs. immature erythroid precursors (IEP, CD71^high^Ter119^low^)(Koulnis et al., 2011). The percentage of IEP rose from 1% in Ctrl to 12.5% in cKO (**Fig. 1c, I**), while MEP decreased from 30% to 19% (**Fig. 1c, II**). Further analysis showed that the residual MEP in cKO embryos retained at least one intact *Drosha* allele and expressed *Drosha* mRNA at a level comparable to Ctrl-MEP, confirming that the presence of Drosha is critical for the maturation of EPC. In older embryos (E13.5), IEP remained predominant in *Drosha* cKO (**Fig. 1d, I**: 29.1% cKO vs 7.3% Ctrl) at the expense of MEP (**Fig. 1d, II/III:** 15.4% cKO vs 80.6% Ctrl). When cultured in erythrocyte differentiation media for 3 days, >95% of cKO-IEP remained morphologically immature, while ∼25% of Ctrl-IEP developed a mature erythrocyte morphology (**Supplementary Fig. S1b)**. These results suggest that in the absence of *Drosha*, EPC fail to differentiate into erythrocytes.

**Figure 1:**
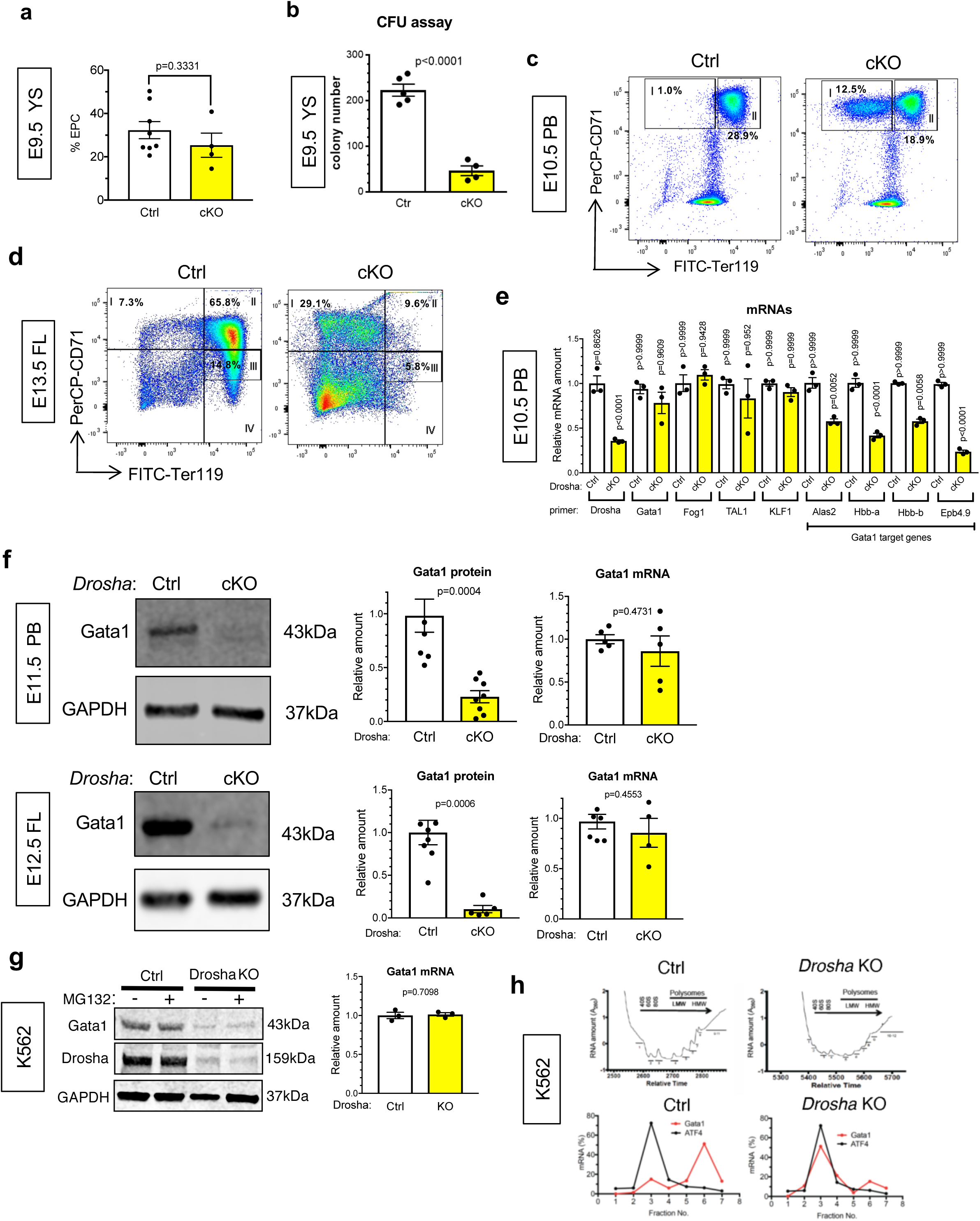
Endothelial-specific deletion of mouse *Drosha* impairs erythropoiesis. **a.** Erythroblasts (CD71^+^) derived from E9.5 yolk sac (YS), quantitated by flow cytometry, are shown as frequency (%) of total live (DAPI^-^) cells (Mean ± SEM). NS, not significant (the statistical significance of all the data presented in this and in the following figures was evaluated by two-tailed unpaired Student’s t test). n=Ctrl: 6 embryos; cKO: 4 embryos. 2 litters. **b.** Total E9.5 yolk sac cells from Ctrl (*Drosha*^fl/+^; *Cdh5-cre*^+^ or *Drosha*^fl/fl^, or Drosha^fl/+^) or cKO (*Drosha*^fl/fl^; *Cdh5-cre*^+^) were subjected to colony formation (CFU) assay in duplicates. Colony counts of the three progenitors (BFU-E, CFU-GM, and CFU-GEMM) were plotted as Mean ± SEM; n=Ctrl: 5 embryos; cKO: 4 embryos. 1 litter. **c.** Representative images of flow cytometry analyses on pro-erythroblasts (I: CD71^high^Ter119^low^) and erythroblasts (II: CD71^high^Ter119^high^) derived from peripheral blood (PB) of E10.5 Ctrl or cKO embryos (upper panel). Mean fraction (%) of each population per total live (DAPI^-^) cells are indicated. n=Ctrl: 14 embryos; cKO: 4 embryos. 2 litters. **d.** Representative images of flow cytometry analysis on erythroblasts at different stages (I: CD71^high^Ter119^low^; II: CD71^high^Ter119^high^; III: CD71^mid^Ter119^high^; IV: CD71^low^Ter119^high^) from E13.5 fetal liver (FL) of Ctrl or cKO embryos (upper panel). Mean fraction (%) of each population per total live (DAPI^-^) cells are indicated. n=Ctrl: 8 embryos; cKO: 4 embryos. 3 litters. **e.** qRT-PCR analysis of different mRNAs (relative to GAPDH) in erythroid population of the PB of E10.5 Ctrl and cKO embryos. (Mean ± SEM). n=Ctrl: 5 embryos; cKO: 5 embryos. 3 litters. **f.** Gata1 protein amount was measured by western blot using CD71^+^Ter119^+^ cell from the PB of E11.5 Ctrl and cKO embryos, or of fetal liver (FL) from E12.5 Ctrl or cKO embryos. Quantitation is plotted on the right. For E11.5 PB, n=Ctrl: 3 embryos; cKO: 4 embryos. 2 litters. For E12.5 FL, n=Ctrl: 5 embryos; cKO: 3 embryos. 2 litters. Gata1 mRNA was evaluated by qRT-PCR in total RNA prepared from the PB of E11.5 Ctrl or cKO embryos, or of FL from E12.5 Ctrl or cKO embryos. For E11.5 PB, n=Ctrl: 5 embryos; cKO: 4 embryos. 2 litters. For E12.5 FL, n=Ctrl: 6 embryos; cKO: 3 embryos. 2 litters. **g.** K562 cells expressing CRISPR/Cas9 targeting *Drosha* (*Drosha* KO) or non-specific control (Ctrl) were treated with or without proteasome inhibitor MG132 (5 nM) for 6 hrs prior to the preparation of total cell lysates, followed by immunoblot with anti-Gata1 and anti-GAPDH (loading control) antibody. **h.** Polysome fraction of Ctrl or *Drosha* KO K562 cells. Upper panel: polysome profile of Ctrl or *Drosha* KO K562 cells. Lower panel: qRT-PCR analysis of Gata1 and ATF4 mRNAs from each fraction normalized by 18S rRNA.

To elucidate the molecular mechanism underlying the maturation defects of EPC in *Drosha*-cKO embryos (cKO-EPC), we investigated the expression in IEP (CD71^high^Ter119^low^) of transcription factors involved in erythroid differentiation, such as Gata binding protein 1 (Gata1) (Pevny et al., 1995; Pevny et al., 1991; Suzuki et al., 2011; Zon et al., 1991), Friend of Gata1 (Fog1, also known as ZFPM1)(Welch et al., 2004), T cell acute lymphocytic leukemia 1 (TAL1, also known as SCL)(Aplan et al., 1992; Green et al., 1991), and Krüppel-like factor 1 (KLF1, also known as EKLF)(Coghill et al., 2001). All these transcription factors mRNAs were comparably expressed in cKO and Ctrl, although Drosha mRNA in cKO was 70% lower than Ctrl, as expected (**Fig. 1e**.) However, Gata1 protein—abundant in Ctrl—was undetectable in cKO (**Fig. 1f).** Furthermore, transcripts of Gata1 target genes—such as *Alas2*, *Hbb-a*, *Hbb-b*, and *Epb4.9* (Campbell et al., 2013)—were reduced in cKO compared to Ctrl (**Fig. 1e, Gata1 targets**), which is consistent with the loss of Gata1 protein. We observed a similar reduction of Gata1 protein (**Fig. 1f, left**), but not Gata 1 mRNA (**Fig. 1f, right**), in human erythroleukemia K562 cells in which the *Drosha* gene was mutated by CRISPR/Cas9 (*Drosha* KO cells). Loss of Gata1 protein in *Drosha* KO cells was not reversed by the proteasome inhibitor MG132 (**Fig. 1g, left**), suggesting that a mechanism other than protein degradation might be responsible for Gata1 depletion. Cell growth analysis showed that *Drosha* KO cells grew at a rate ∼50% lower than Ctrl K562 cells, but appeared otherwise normal (**Supplementary Fig. S2A**). MiR-451, which is critical for erythroid differentiation(Patrick et al., 2010; Rasmussen et al., 2010), was abundantly expressed in *Drosha* KO cells (**Supplementary Fig. S2b).** K562 cells can undergo partial differentiation into benzidine-positive (blue) mature erythrocytes when treated with hemin, a ferric (Fe^3+^) form of heme(Tomoda et al., 1991),(Hafner et al., 1995). While ∼60% Ctrl K562 cells turned benzidine-positive, *Drosha* KO treated with hemin remained benzidine-negative, a sign of undifferentiated state (**Supplementary Fig. S2c**), confirming a requirement of Drosha for erythrocyte differentiation, as seen in cKO mice (**Fig. 1c and 1d**). Furthermore, a polysome fractionation analysis showed *Gata1* mRNA was enriched in the high molecular weight polysome fractions in Ctrl K562 cells, indicating active translation **(Fig. 1h, bottom left, red line)**. In *Drosha* KO cell, however, *Gata1* mRNA was augmented in the monosome fractions overlapping *ATF4* mRNA, which are translationally inhibited in cells under regular growth condition(Pakos-Zebrucka et al., 2016) (**Fig. 1h, bottom right, black line**)—a sign of translational inactivation **(Fig. 1h, bottom right, red line)**. Thus, we conclude that depletion of Drosha in erythroid progenitors results in decreased Gata1 translation, which impairs differentiation.

### The Microprocessor complex is required for the expression of ribosomal proteins

To investigate how the expression of Gata1 is attenuated upon depletion of Drosha, we performed a transcriptome analysis. RNA-seq analysis was performed on EPC from E10.5 Ctrl and cKO embryos. As expected, the amount of Drosha mRNA was reduced in cKO-EPC compared to Ctrl-EPC (**Fig. 2a and Supplementary Fig. S3a)**. Depletion of Drosha in cKO-EPC was also validated by an increase of transcripts processed and degraded by Drosha—such as *Dgcr8*, *Anks6*, and *Stat6*(Kim et al., 2017) (**Supplementary Fig. S3b**). ∼60% of transcripts were increased in cKO-EPC compared to Ctrl-EPC, likely reflecting a global reduction of miRNAs and de-repression of their mRNA targets (**Supplementary Fig. S3a**). DAVID pathway analysis of transcripts revealed that mRNAs tagged as “ribosome” were the most significantly reduced, with 68 out of 71 small (40S) and large (60S) ribosomal protein (Rps and Rpl) transcripts down on average by 25% in cKO-EPC compared to Ctrl-EPC (**Fig. 2a**), a result confirmed by qRT-PCR analysis (**Fig. 2b**).

**Figure 2.**
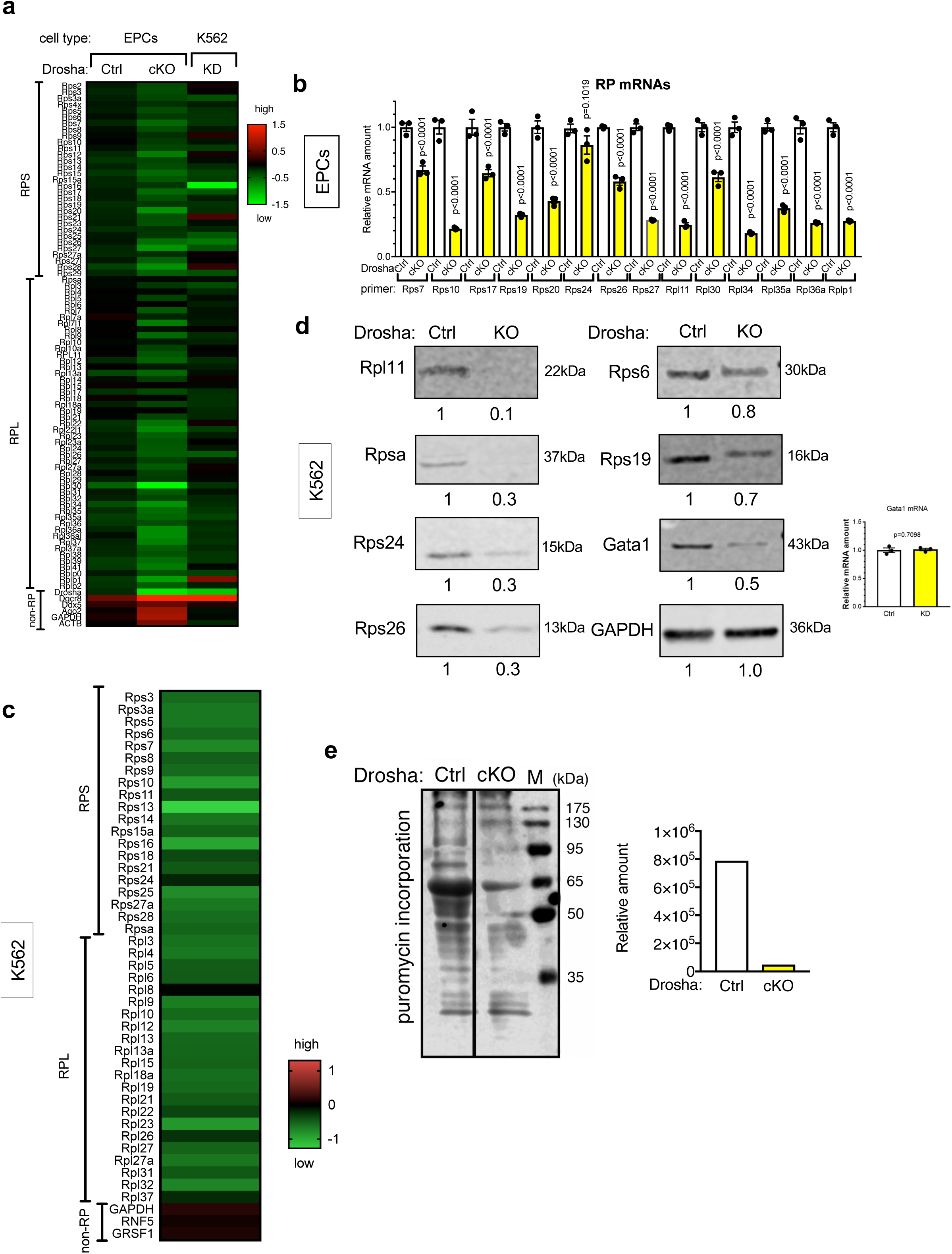
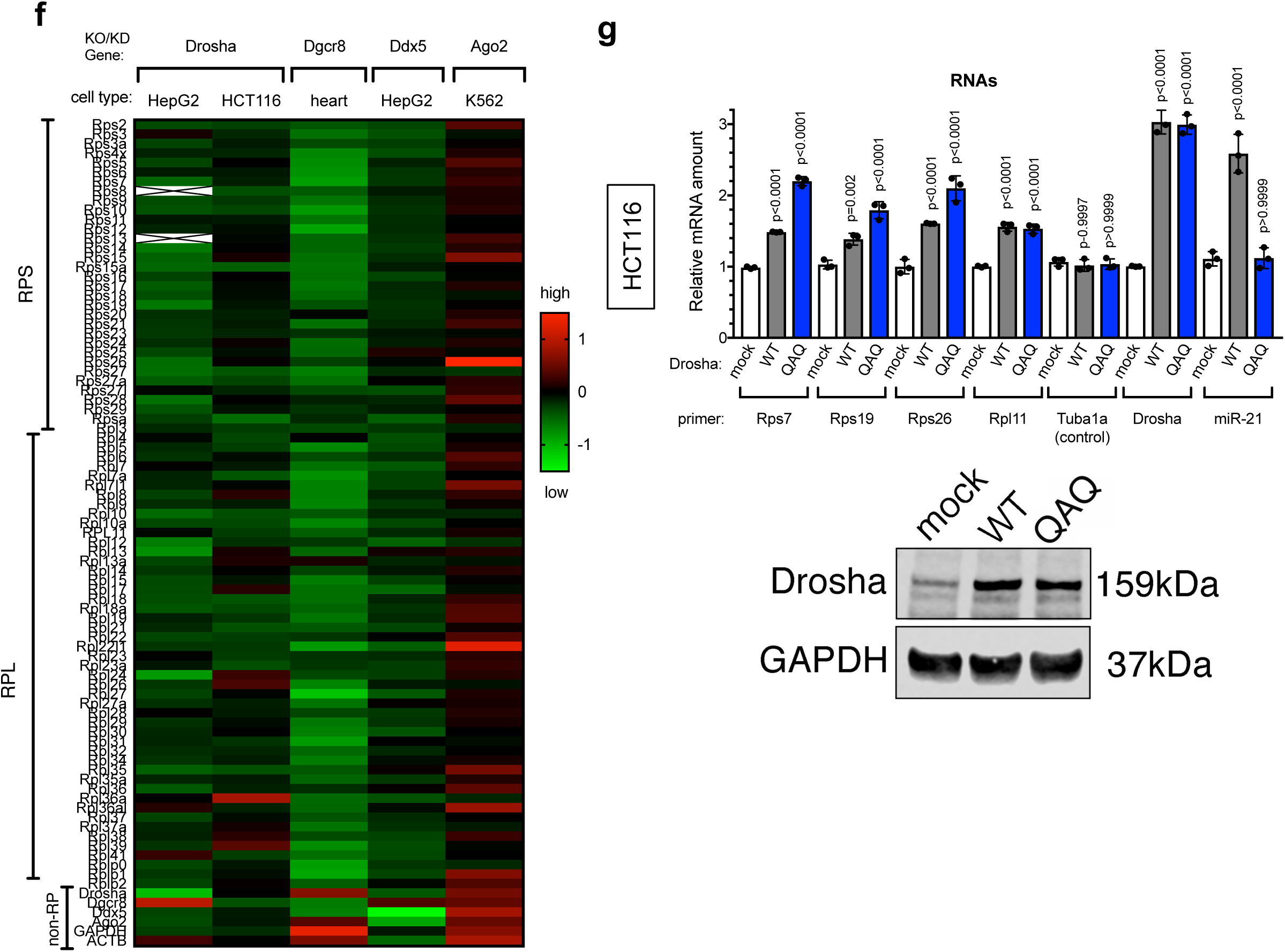
Reduced translation of Gata1 in *Drosha*-null erythroid progenitors. **a.** Heatmap illustrates changes in mRNAs encoding ribosomal proteins in EPC (CD71^high^Ter119^+^) sorted from E10.5 peripheral blood of Ctrl and cKO mice. Heatmap illustrating changes in mRNAs encoding ribosomal proteins in *Drosha* KD K562 cells versus Ctrl K562 cells is plotted in the right lane. Log _2_(fold changes relative to ctrl) is plotted. **b.** qRT-PCR analysis of mRNAs (relative to GAPDH) in EPC sorted from the peripheral blood of Ctrl and cKO mice (E10.5). (Mean ± SEM). n=Ctrl: 5 embryos; cKO: 5 embryos. 3 litters. **c.** Equal number (3×10^5^ cells) of Ctrl and Drosha KO K562 cells were subjected to TMT-based mass-spectrometric analysis analysis. n=5. Log _2_(fold changes relative to ctrl) is plotted. of RP proteins or negative control protein amount in KO vs Ctrl cells are plotted as a heatmap. **d.** Ribosomal proteins were examined by western blot of total cell lysates from Ctrl or *Drosha* KO K562 cells. The relative quantitation is in the label beneath each lane. Gata1 mRNA was analyzed by qRT-PCR in Ctrl or Drosha KO K562 cells, and plotted on the right as Mean ± SEM. NS, not significant; n=3. **e.** Total cell lysates were generated from an equal number (3X10^5^ cells) of EPCs from the PB of E11.5 Ctrl and cKO embryos were subjected to immunoblot analysis with anti-puromycin antibody (left). Intensity of protein bands were quantitated by ImageJ (right). n=Ctrl: 5 embryos; cKO: 5 embryos. 3 litters. M stands molecular weight marker. **f.** Heatmap of changes in the amount of ribosomal proteins transcripts. The name of the gene that is knockdown is labelled on top. The cell type is labelled at bottom. Log 2(fold changes relative to ctrl) is plotted. Crossed (X) cells: lack of data. **g.** Human HCT116 cells were transfected with a vector (mock), a wild type Drosha (WT) or the RNase-defective mutant (QAQ) construct, followed by qRT-PCR analysis of Rps/l mRNAs (relative to b-actin), Tuba1a (control), Drosha and miR-21. Mean ± SEM. n=3.

Like in primary mouse EPC, Rps and Rpl mRNAs also declined in K562 cells with silenced *Drosha* (**Fig. 2a, KD**). Tandem mass-tag (TMT)-based quantitative proteomic analysis (**Fig. 2c and Supplementary Fig. S4**) and immunoblots (**Fig. 2d**) confirmed that the majority of Rps and Rpl proteins decreased in Drosha-depleted K562 cells. To examine how the rate of global protein synthesis is affected as a result of reduced RP expression in *Drosha* null cells, we performed an in vivo puromycin incorporation assay (also known as puromycin-associated nascent chain proteomics; PUNCH-P)(Aviner et al., 2013). Puromycin is an analog of tyrosyl-tRNA that is incorporated into nascent polypeptide chains, allowing the measurement of global protein synthesis by detection of puromycin-labeled proteins with an anti-puromycin antibody (Aviner et al., 2013; Park et al., 2016). An equal number of EPC was sorted from the PB of E11.5 Ctrl or cKO embryos from puromycin-injected pregnant female mice. The amount of puromycin-labeled protein was 90% lower in cKO-EPC compared to Ctrl-EPC (**Fig. 2e, lane 1 vs 2**), indicating a reduction of the rate of protein synthesis in cKO-EPC. Thus, we conclude that Drosha depletion causes a decrease of RPs and general reduction of protein synthesis. The degree of the effect of ribosome insufficiency on translation is mRNA-dependent(Ludwig et al., 2014), and *Gata1* translation appears to be more severely attenuated than that of other housekeeping gene transcripts in EPC and K562 cells. A search of published RNA-seq data revealed that Rps and Rpl transcripts are also decreased in non-erythroid cells—such as human hepatocarcinoma HepG2 and human colon carcinoma HCT116 cells (**Fig. 2f**)—upon silencing of *Drosha* by RNAi (KD)(Kim et al., 2016), indicating that the Drosha-RP regulatory axis is not confined to erythroid lineages.

We next tested the involvement of other subunits of the Microprocessor complex in the regulation of RP expression. We observed that both DGCR8—a key partner of Drosha—and the DEAD-box RNA helicase Ddx5—an auxiliary subunit of the Microprocessor complex—are required for RP gene regulation, since nearly all Rps and Rpl mRNAs were reduced in the embryonic heart of E9.5 homozygous *Dgcr8* cKO mice (Dgcr8 cKO; *Dgcr8*^loxP/loxP^: *Mesp1*^Cre/+^)(Chen et al., 2019) and in Ddx5-depleted HepG2 cells (**Fig. 2f, Dgcr8 and Ddx5**). Conversely, and unexpectedly, depletion of Argonaute2 (Ago2), a component of the RNA silencing complex (RISC) that uses miRNAs to inhibit mRNA expression, resulted in a small increase, rather than a decrease, of Rps and Rpl transcripts in K562 cells (**Fig. 2f, Ago2**). If impairment of miRNA synthesis and function were responsible for the RP synthesis block, we would have expected a similar result from depletion of Microprocessor and RISC components. However, this result suggests that miRNA biogenesis may not be the main mechanism driving the regulation of RP genes by the Drosha. To investigate further the possibility of miRNA-independent control of RPGs by the Microprocessor complex, we expressed in HCT116 cells a ribonuclease-defective Drosha mutant (QAQ; R^938^K^939^K^940^ to QAQ), which is unable to process pri-miRNAs(Kwon et al., 2016), and measured the amount of RP transcripts. The Drosha (QAQ) mutant, expressing levels of mRNA (**Fig. 2g, top)** and protein (**Fig. 2g, bottom)** similar to wild type Drosha (WT), increased Rps and Rpl mRNAs as effectively as Drosha(WT) (**Fig. 2g, top**) while the level of miR-21 was increased by Drosha(WT) but not by Drosha(QAQ) mutant (**Fig. 2g, top)**, confirming that miRNA processing by the Microprocessor complex is dispensable for RP gene regulation. Therefore, we explored alternative mechanisms by which the Microprocessor complex might regulate RPGs.

### The Microprocessor complex binds to the transcription start site of RP gene loci

Chromatin immunoprecipitation-DNA sequencing (ChIP-seq) analysis indicated that both Drosha and Dgcr8 interact with the genomic loci proximal to the transcription start site (TSS) of all 80 RP genes (**Supplementary Fig. S5 and Table 1**) (Suzuki et al., 2017). The Drosha and Dgcr8 association sites within the *Rps15a*, *Rps24*, *Rpl4*, and *Rpl28* loci overlap with the RNA polymerase II (RNAPII) binding sites and is marked by histone H3 Lysine (K) 4 trimethylation (H3K4me3), which is indicative of transcriptionally active chromatin (**Fig. 3a**)(Gromak et al., 2014; Suzuki et al., 2017). We validated the association of Drosha with RPG loci by ChIP-qPCR assay in MEFs (**Fig. 3b**). Transcription inhibition by Actinomycin D (ActD) abolished the association of Drosha with RPG loci (**Fig. 3c**), as did RNA digestion with ribonuclease A (RNase A) (**Fig. 3d**), indicating the involvement of RNA in the interaction between Drosha and RPGs. Furthermore, when ChIP samples were pretreated with ribonuclease H (RNase H), which specifically degrades the RNA in a DNA/RNA hybrid, Drosha association with Rps/Rpl loci was abolished, suggesting that Drosha interacts with the nascent RP transcript on or near R-loops, which are composed of a DNA/RNA hybrid of template DNA and nascent mRNA and a single-stranded non-template DNA (**Fig. 3e**). Dgcr8 is not essential for Drosha association to RP gene loci because Drosha enrichment was unaffected in *Dgcr8* homozygous-null MEFs (DGCR8 KO) compared to wild type control MEFs (Ctrl) (**Fig. 3f**). However, the amounts of Rps/Rpl mRNAs were reduced in Dgcr8 KO cells (**Fig. 3g**), suggesting that Dgcr8 is required for transcriptional regulation of RP genes at a stage that follows the binding of Drosha to RPG loci. These results are consistent with a model in which the Microprocessor associates with the newly synthesized Rps/Rpl mRNAs via Drosha.

**Figure 3.**
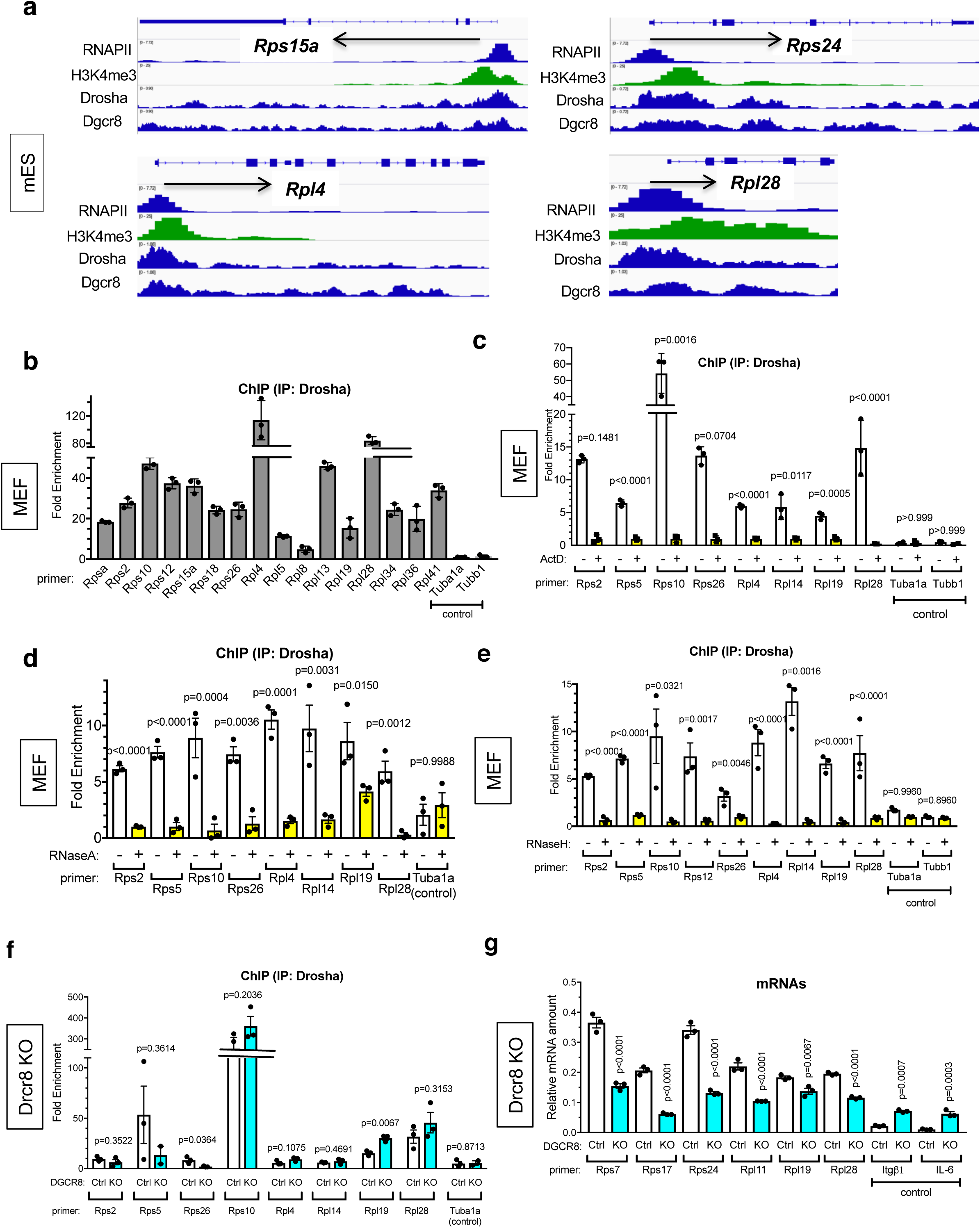
The microprocessor binds to the transcription start sites at RP gene loci. **a.** ChIP-seq profiles of RNAPII, H3K4me3, Drosha and Dgcr8 at the Rps15a, Rps24, Rpl4, and Rpl28 loci in mouse embryonic stem cells (mES). **b.** ChIP (IP: Drosha)-qPCR analysis of different RP genes as indicated with anti-Flag (M2) antibody or nonspecific IgG (control) in Flag tagged Drosha-expressing MEF (F-Drosha) or control (pBABE-MEF). Tuba1a and Tubb1 (negative control). Fold enrichment of anti-Flag IP over IgG IP is plotted as Mean ± SEM. n=3.**c.** ChIP (IP: Drosha)-qPCR analysis of different RP genes as indicated with anti-Flag (M2) antibody or nonspecific IgG (negative control) in Flag-Drosha-expressing MEF or control-MEF. Tuba1a and Tubb1 (negative control). Cells were treated with 1μg/ml actinomycin D (ActD) or vehicle (DMSO), followed by ChIP-qPCR analysis. Fold enrichment of anti-Flag IP over IgG IP is plotted as Mean ± SEM. n=3.**d.** ChIP (IP: Drosha)-qPCR analysis of different RP genes as indicated in MEF treated with 1μg/μl RNase A or vehicle (water) with anti-Drosha antibody or nonspecific IgG (control). Tuba1a and Tubb1 (negative control). Fold enrichment of Drosha IP over IgG IP is plotted as Mean ± SEM.n=3. **e.** ChIP (anti-Drosha antibody)-qPCR analysis of different RP genes in MEFs treated with 100U/ml RNase H or vehicle (water) with anti-Drosha antibody. Tuba1a and Tubb1 (negative control). Fold enrichment of Drosha IP over IgG IP is plotted as Mean ± SEM.n=3. **f.** ChIP (IP: Drosha)-qPCR analysis of different RP genes as indicated with anti-Drosha antibody or nonspecific IgG (control) in control MEFs (Ctrl) or MEFs deleted in Dgcr8 gene (Dgcr8-KO). Tuba1a (negative control). Fold enrichment of Drosha IP over IgG IP is plotted as Mean ± SEM.n=3. **g.** qRT-PCR analysis of different RP mRNAs and control mRNAs (Itgb1 and IL-6) (relative to GAPDH) as indicated in Ctrl or Dgcr8-KO MEFs. Result is plotted as Mean ± SEM. n=3.

**Table 1:**
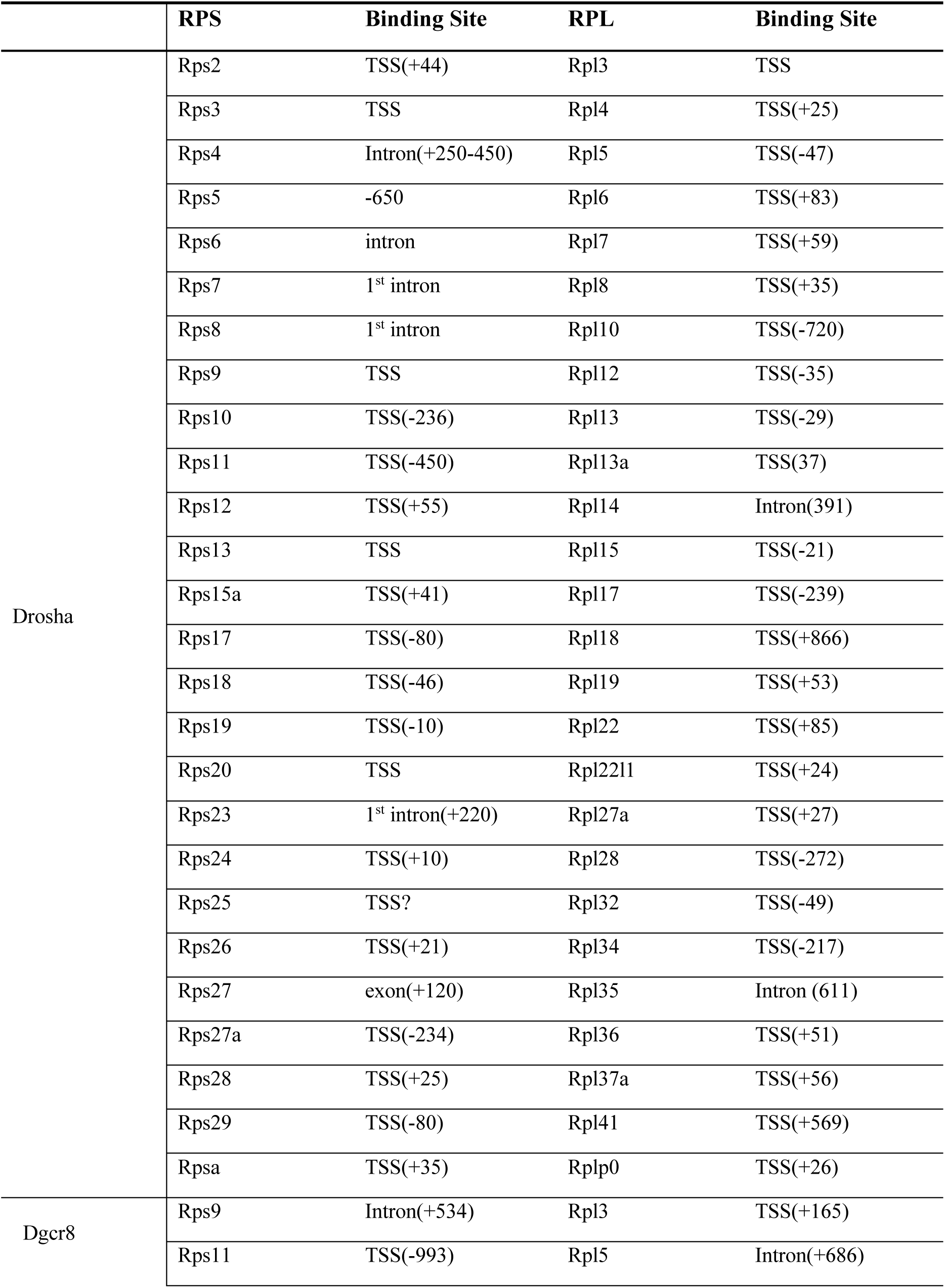
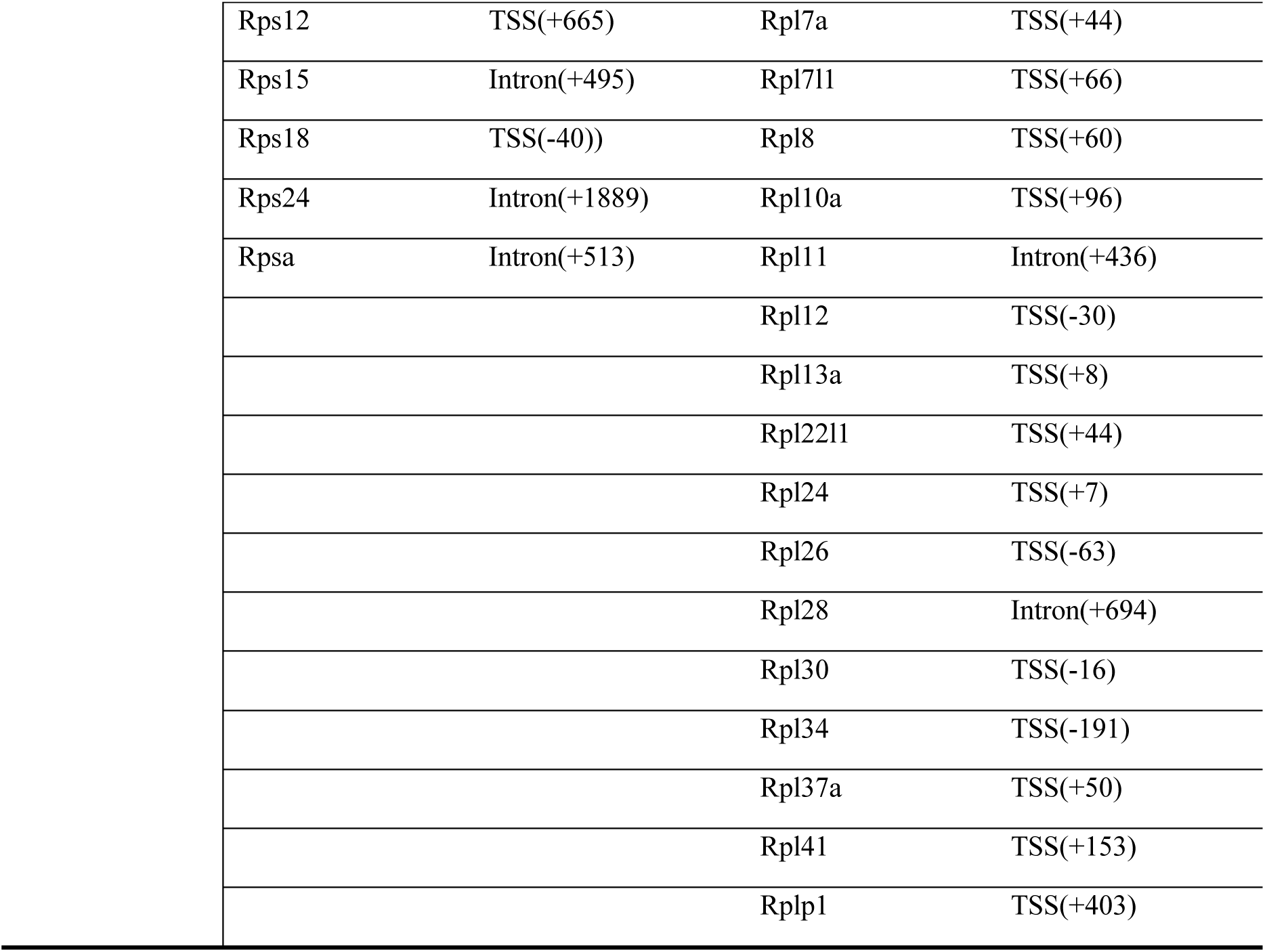
Association of Drosha/Dgcr8 with Rps/Rpl Transcription Start Site (TSS) by ChIP-seq.

|  | **RPS** | **Binding Site** | **RPL** | **Binding Site** |
| --- | --- | --- | --- | --- |
| Drosha | Rps2 | TSS(+44) | Rpl3 | TSS |
|  | Rps3 | TSS | Rpl4 | TSS(+25) |
|  | Rps4 | Intron(+250-450) | Rpl5 | TSS(-47) |
|  | Rps5 | -650 | Rpl6 | TSS(+83) |
|  | Rps6 | intron | Rpl7 | TSS(+59) |
|  | Rps7 | 1<sup>st</sup> intron | Rpl8 | TSS(+35) |
|  | Rps8 | 1<sup>st</sup> intron | Rpl10 | TSS(-720) |
|  | Rps9 | TSS | Rpl12 | TSS(-35) |
|  | Rps10 | TSS(-236) | Rpl13 | TSS(-29) |
|  | Rps11 | TSS(-450) | Rpl13a | TSS(37) |
|  | Rps12 | TSS(+55) | Rpl14 | Intron(391) |
|  | Rps13 | TSS | Rpl15 | TSS(-21) |
|  | Rps15a | TSS(+41) | Rpl17 | TSS(-239) |
|  | Rps17 | TSS(-80) | Rpl18 | TSS(+866) |
|  | Rps18 | TSS(-46) | Rpl19 | TSS(+53) |
|  | Rps19 | TSS(-10) | Rpl22 | TSS(+85) |
|  | Rps20 | TSS | Rpl22l1 | TSS(+24) |
|  | Rps23 | 1<sup>st</sup> intron(+220) | Rpl27a | TSS(+27) |
|  | Rps24 | TSS(+10) | Rpl28 | TSS(-272) |
|  | Rps25 | TSS? | Rpl32 | TSS(-49) |
|  | Rps26 | TSS(+21) | Rpl34 | TSS(-217) |
|  | Rps27 | exon(+120) | Rpl35 | Intron (611) |
|  | Rps27a | TSS(-234) | Rpl36 | TSS(+51) |
|  | Rps28 | TSS(+25) | Rpl37a | TSS(+56) |
|  | Rps29 | TSS(-80) | Rpl41 | TSS(+569) |
|  | Rpsa | TSS(+35) | Rplp0 | TSS(+26) |
| Dgcr8 | Rps9 | Intron(+534) | Rpl3 | TSS(+165) |
|  | Rps11 | TSS(-993) | Rpl5 | Intron(+686) |

|  |  |  |  |
| --- | --- | --- | --- |
| Rps12 | TSS(+665) | Rpl7a | TSS(+44) |
| Rps15 | Intron(+495) | Rpl7l1 | TSS(+66) |
| Rps18 | TSS(-40)) | Rpl8 | TSS(+60) |
| Rps24 | Intron(+1889) | Rpl10a | TSS(+96) |
| Rpsa | Intron(+513) | Rpl11 | Intron(+436) |
|  |  | Rpl12 | TSS(-30) |
|  |  | Rpl13a | TSS(+8) |
|  |  | Rpl22l1 | TSS(+44) |
|  |  | Rpl24 | TSS(+7) |
|  |  | Rpl26 | TSS(-63) |
|  |  | Rpl28 | Intron(+694) |
|  |  | Rpl30 | TSS(-16) |
|  |  | Rpl34 | TSS(-191) |
|  |  | Rpl37a | TSS(+50) |
|  |  | Rpl41 | TSS(+153) |
|  |  | Rplp1 | TSS(+403) |

### Recruitment of the Microprocessor to the 5’-TOP motif of RP transcripts promotes transcription elongation

Association of Drosha and Dgcr8, but not Polypyrimidine binding protein 1 (Ptbp1, negative control), with the 5’-untranslated region (UTR) of RP mRNAs was detected by analysis of ENCODE datasets (Boucas, 2018; Consortium, 2012; Davis et al., 2018) of enhanced UV crosslinking followed by immunoprecipitation (eCLIP) in K562 cells (**Fig. 4a and Supplementary Fig. S6).** Neither Drosha nor Dgcr8 were found to interact with control mRNAs, such as tubulins (**Supplementary Fig. S6)**. All Rps/Rpl genes in metazoa contain the 5’-TOP motif, a pyrimidine-rich stretch that plays a critical co-regulatory role in the expression of RP genes through various mechanisms (Hamilton et al., 2006; Meyuhas and Kahan, 2015; Perina et al., 2011; Rojas et al., 2018). A functionally equivalent TOP motif has been found in a limited set of non-RP genes (non-RP TOP genes)(Meyuhas and Kahan, 2015). eCLIP data showed that Drosha and Dgcr8 also associate with the mRNA of non-RP TOP genes, including *Nucelophosmin1* (*NPM1*), *eukaryotic initiation factor 3F* (*eIF3F*), *eIF4B*, *heterogeneous nuclear ribonucleoprotein A1* (*hnRNPA1*), *nucleosome assembly protein1 like1* (*NAP1L1*), and *Vimentin* (*VIM*) (**Supplementary Fig. S6)**. Therefore, we speculated that the TOP motif might play a critical role in the association of the Microprocessor with a subset of mRNAs, including RPG mRNAs. A RNA secondary structure prediction algorithm (Vienna RNAfold) detects a stable stem-loop structure in the first 37-nt sequence of the *Rpl28* mRNA (**Fig. 4b)**. Electrophoretic mobility shift assay (EMSA) confirmed that the double-strand RNA binding domain (RBD; amino acid 1259-1337) of Drosha is sufficient to bind the 37-40-nt sequence (WT) of the *Rpl28* (**Fig. 4b, lanes 4-6**), Rps13, and Rpl4 mRNA (**Supplementary Fig. S11**), but fails to bind the TOP1 mutant probe, in which three nucleotides within the TOP motif have been mutated from UUU to AAA to disrupt both the TOP motif and the stem structure (**Fig. 4b, lanes1-3).** An excess of unlabeled TOP1 RNA probe did not prevent Drosha binding to the WT probe (**Fig. 4b, lanes 12 and 13**), unlike the WT competitor (**Fig. 4b, lanes 10 and 11**), confirming that Drosha does not bind the TOP1 RNA mutant. When we introduced additional mutations in TOP1 to create TOP1stem, in which the TOP sequence is still mutated, but the complementary strand of the stem structure has been restored, the TOP1stem RNA once again efficiently competed for Drosha binding to the WT probe (**Fig. 4b, lanes 14 and 15**). On the other hand, the Down mutant, in which the stem structure downstream of the TOP motif was disrupted without mutating the TOP motif, was unable to compete for Drosha binding (**Fig. 4b, lanes 16 and 17**). Thus, the stem structure at the 5’ terminus, but not the TOP sequence proper, is required for Drosha association with the *Rpl28* mRNA.

**Figure 4.**
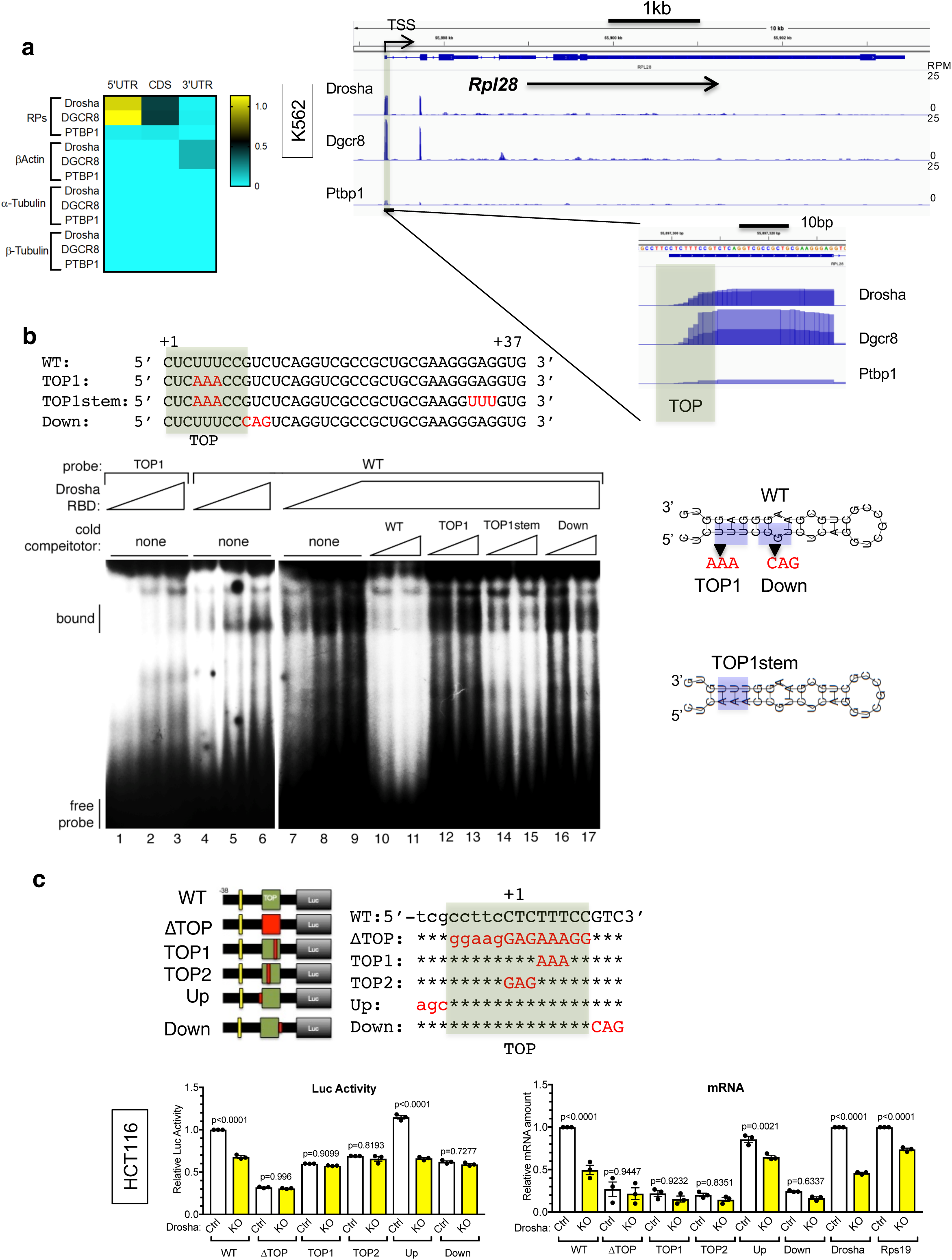
Recruitment of the Microprocessor to the 5’ TOP motif of RP transcripts facilitates RP expression. **a.** Heatmap illustrating the number of peaks per 100-nt in e-CLIP profile of all the RPs, β-Actin, α-Tubulin or β-Tubulin (left). e-CLIP profiles of Drosha, Dgcr8 and PTBP1 at the Rpl28 locus in K562 cells (right). The underlined profile is magnified in the right bottom. **b.** Representative image of RNA EMSA with Drosha RBD domain, radiolabeled WT or TOP1 probes, and unlabeled (cold) competitors. Probe and competitor sequences are shown above the RNA EMSA, the predicted secondary structures of the probes are shown on the right. **c.** Luciferase assay was performed in HCT116 cells expressing CRISPR/Cas9 targeting *Drosha* (*Drosha* KO) or non-specific control (Ctrl). The Rpl28 promoter was inserted upstream of the luciferase gene. The numbers are relative to the transcription starting site (+1). In the construction structures, green boxes indicate WT sequences, red boxes indicate mutant sequences. Luciferase activity is plotted as Mean ± SEM on the bottom left panel. n=3. qRT-PCR analysis of Drosha and Rps19 mRNAs (relative to GAPDH) is plotted as Mean ± SEM on the bottom right panel. n=3.

To examine the functional significance of the association of Drosha with the 5’-end of RP mRNAs, a luciferase (luc) reporter construct containing the TATA box and the TOP motif of the *Rpl28* gene was transfected into control HCT116 cells (Ctrl) or cells in which the *Drosha* gene had been deleted by CRISPR/Cas9 (*Drosha* KO). Both the reporter activity and the amount of luciferase mRNA were lower in *Drosha* KO cells compared to Ctrl cells (**Fig. 4c, WT-luc**), indicating that the WT-luc reporter recapitulates the Drosha-dependent transcriptional regulation of *Rpl28* (**Fig. 4c, WT-luc**). The Up mutant (in which the nucleotides upstream of the TSS were mutated) had no effect and displayed a reporter activity and mRNA level nearly identical to WT (**Fig. 4c**).

Inversely, all three TOP motif mutations tested exhibited reduced luc activity and mRNA level, but also failed to respond to Drosha depletion: ΔTOP (in which the entire TOP sequence was mutated), TOP1, which failed to bind Drosha-RBD by EMSA (**Fig. 4b**), and TOP2 (with mutations in the first three nucleotides of the transcript within the TOP motif) (**Fig. 4c,** ΔTOP, TOP1, and TOP2). The Down mutant, which failed to bind Drosha by EMSA (**Fig. 4b**), also lost Drosha-dependency (**Fig. 4c, Down-luc**). Thus, both the presence of Drosha and its ability to bind the mRNA were required for maximal transcription and expression of the luciferase reporter. These results indicate that Drosha controls the amount of RP mRNAs via the TOP motif, which is shared among all RP mRNAs.

### The Ddx5 helicase reduces R-loops and facilitates transcription elongation

Stable formation during transcription of an R-loop, which is composed of the template DNA/nascent mRNA hybrid and the displaced non-template DNA strand, is inhibitory to elongation by RNAPII(García-Muse and Aguilera, 2019; Gowrishankar et al., 2013; Nudler, 2012). R-loops formed in a GC-rich region are stable and can impede transcription(Ginno et al., 2013; Huertas and Aguilera, 2003). Since a GC-rich sequence of 5-20 bp is often present immediately downstream of the TOP motif in RP genes(Meyuhas and Kahan, 2015), we hypothesized that there is an abundance of stable R-loops at the RP loci, which might be inhibitory to RPG transcription, especially in actively proliferating cells, and may require a mechanism to resolve R-loops, possibly in a Drosha-dependent manner.

DRIP-seq data validated the presence of R-loops at the RPG loci (**Supplementary Fig.S7**). Analysis of genome-wide mapping of R-loops (DRIP-seq) in human U-2 OS cells using the S9.6 antibody, which specifically recognizes DNA/RNA hybrids, indicated that upon depletion of *Drosha* R-loops increased at RPG loci, but unchanged at control loci (GAPDH and Tuba1a) (**Fig. 5a)**(Lopez-Carballo et al., 2002). Similar results were obtained by DRIP assay in K562 cells (**Fig. 5b, Ctrl vs KO**). The DRIP signal was abolished when DNA/RNA hybrids were degraded by RNase H, confirming a specific recognition of DNA/RNA hybrids by the S9.6 antibody (**Fig. 5b**). These results support an essential role of Drosha in the resolution of R-loops specifically at RPG loci. A ChIP assay showed the amount of RNAPII associated with the 3’-end of RPGs was ∼40% lower in Drosha KO cells compared in Ctrl cells (**Fig. 5c**). But there was no change in RNPAII association at control loci (**Fig. 5c**), indicating that RNAPII elongation is interfered by the R-loops accumulated at RPG loci, leading to a reduced rate of transcription of RPGs.

**Figure 5.**
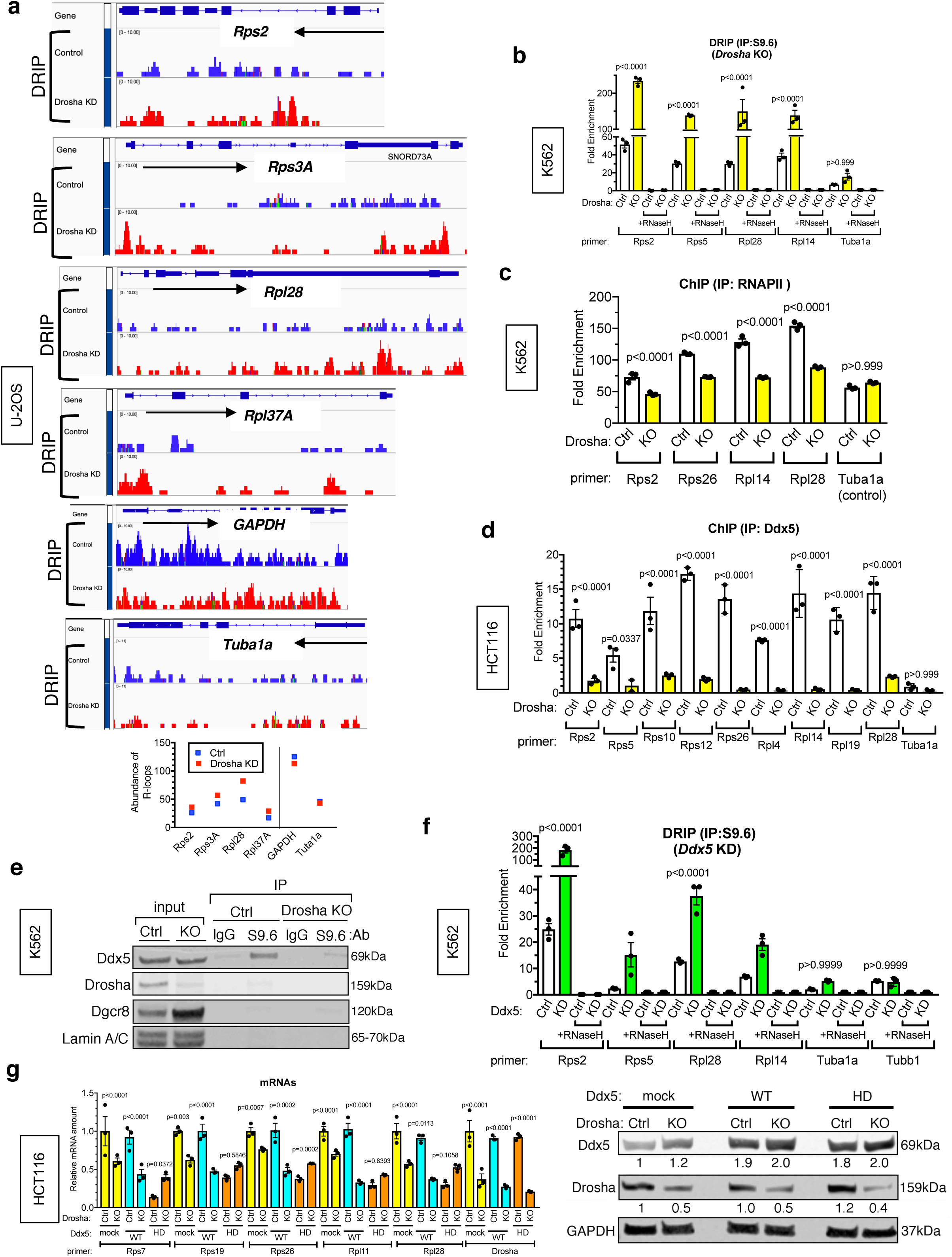
The Ddx5 helicase reduces DNA/RNA hybrid and facilitates transcription elongation. **A.** DNA/RNA hybrid IP-sequencing (DRIP-seq) data indicate increased R-loops at the RPG loci (Rps2, Rps3A, Rpl28, and Rpl37A) and control loci (GAPDH and Tuba1a) in Drosha KD cells (red) compared to control U-2 OS cells (blue) (top). Quantitation of the DRIP-seq data is shown (bottom). **b.** DRIP analysis of RPG loci (Rps2, Rps5, Rpl14) and control locus (Tuba1a) locus in the presence or absence of RNase H in HCT116 cells expressing CRISPR/Cas9 against *Drosha* (KO) or non-specific control (Ctrl). Four sets of primers used for DRIP analysis are indicated as black boxes in the top panel (bottom). Signal relative to input is plotted as Mean ± SEM. n=3 **c.** ChIP-qPCR analysis using anti-RNAPII antibody was performed in K562 cells cells expressing CRISPR/Cas9 against *Drosha* (KO) or non-specific control (Ctrl). Primers for RPG loci (Rps2, Rps26, Rpl14, Rpl28) and control locus (Tuba1a) are shown. Mean ± SEM. n=3. **d.** ChIP-qPCR analysis using anti-Ddx5 antibody of RPG loci (Rps2, Rps5, Rps10, Rps12, Rps26, Rpl4, Rpl14, Rpl19, Rpl28) and control locus(Tuba1a) in control (Ctrl) or *Drosha* KO HCT116 cells. Fold enrichment of Ddx5 antibody pull-down against IgG pull-down is plotted as Mean ± SEM. n=3. **e.** Immunoprecipitation of DNA/RNA hybrids with the S9.6 antibody, followed by immunoblot analysis of Ddx5, Drosha, Dgcr8 and Lamin A/C (negative control) in Ctrl or Drosha KO K562 cells. **f.** DRIP analysis of RPG loci (Rps2, Rps5, Rpl4, Rpl28) and control loci (Tuba1a and Tubb1) locus in the presence or absence of RNase H in K562 cells targeting *Ddx5* gene (*Ddx5* KO) by RNAi or non-specific control (Ctrl). Result is plotted as Mean ± SEM. n=3 **g.** qRT-PCR analysis of various RP mRNAs and Drosha mRNAs (relative to GAPDH) in Ctrl or *Drosha* KO HCT116 cells transfected with empty plasmid (mock), Ddx5 wild type (WT) or the RNA helicase dead (HD) mutant expression plasmid (left). Result is plotted as Mean ± SEM. n=3 Drosha, Ddx5 and GAPDH proteins were examined by western blot in total cell lysates from HCT116 cells (right). Relative protein amount normalized to GAPDH is shown below each blot.

Because (i) silencing of Ddx5 causes RP mRNA reduction (**Fig. 2f, 4^th^ column**) and (ii) an ATP-dependent RNA helicases, Ddx5 is implicated in the resolution of R-loops(Cristini et al., 2018; Mersaoui et al., 2019), we hypothesize that Ddx5, in association with Drosha, might be required to resolve R-loops and promote RP gene expression. A ChIP assay confirmed the association of Ddx5 with RPG loci, which is Drosha-dependent as the association was reduced in Drosha KO cells (**Fig. 5d**). Immunoprecipitation with the S9.6 antibody followed by immunoblot showed that a fraction of Ddx5—but not Drosha or Dgcr8—is associated with R-loops (**Fig. 5e, Ctrl**). Consistently with the ChIP assay (**Fig. 5d**), the association of Ddx5 and R-loops was reduced in *Drosha* KO cells (**Fig. 5e, *Drosha* KO**), indicating that Ddx5 requires Drosha for R-loop interaction. A DRIP assay in K562 cells was performed to further exmine the change of abundance of R-loops with depletion of Ddx5. When Ddx5 was depleted, the abundance of R-loops at the RPG loci became higher than Ctrl cells, while it was unchanged at the control loci (*Tuba1a* and *Tubb1*) (**Fig. 5f, Ctrl vs *Ddx5* KD**), suggesting that Ddx5 is specifically required for the resolution of R-loops at RPG loci. When a RNA helicase-dead mutant of *Ddx5* [Ddx5(HD)] (Lys^144^ to Asn)(Huang et al., 2015) was expressed at a level similar to endogenous Ddx5 in control HCT116 cells (Ctrl) expressing Drosha (**Fig. 5g, right**), Ddx5(HD) acted as a dominant negative and reduced the Rps and Rpl mRNAs compared to Ctrl cells expressing Ddx5(WT) (**Fig. 5g, left**), indicating that the RNA helicase activity of Ddx5 is required to facilitate RP gene transcription. When Ddx5(HD) was expressed in cells in which *Drosha* was deleted by CRISPR/Cas9 (KO), the amount of Rps and Rpl mRNAs remained similar in cells expressing Ddx5(HD) and Ddx5(WT) (**Fig. 5g, left**), suggesting that Drosha and Ddx5 cooperatively regulate RPG expression. Thus, the evidence suggests that the Microprocessor complex interacts with RPG mRNAs via TOP motif and facilitates R-loops resolution and RNAPII elongation.

### Drosha controls RP biosynthesis upon changes in growth condition

Ribosome production is essential for fueling cell growth and proliferation, but its considerable energy costs require it to be tightly controlled and attuned to cellular growth conditions. The synchronized stoichiometric production of RPs permits the energy-efficient assembly of ribosomes(Hamilton et al., 2006), and the TOP motif is vital for the control of RP biosynthesis in line with cell proliferation (Hamilton et al., 2006; Meyuhas and Kahan, 2015; Patursky-Polischuk et al., 2014). To test whether the Microprocessor complex plays a role in the change in RPG expression in response to the cellular growth environment, we cultured K562 cells under low (1%) or normal (10%) serum condition for 6 h. RPs and Gata1 were reduced under low serum, as expected (**Fig. 6a, left**), as well as RP mRNAs (**Fig. 6a. right**). Consistently, the puromycin incorporation assay indicated that global protein synthesis was reduced by 50% in low serum (**Supplementary Fig. S8**). We observed that the amount of Drosha—but not Dgcr8 or Ddx5— was decreased by 40% in low serum (**Fig. 6a, left**). When Flag-tagged Drosha (F-Drosha) was exogenously expressed, RPs remained at the same levels under nutrients-deprivation as in normal growth media, while in control cells expressing only the endogenous Drosha, RPs were markedly reduced upon nutrients-deprivation (**Fig. 6b**). These results show that nutrients-starvation reduces RP mRNA biosynthesis, at least in part, by reducing the amount of Drosha protein, and indirectly the activity of the Microprocessor complex.

**Figure 6.**
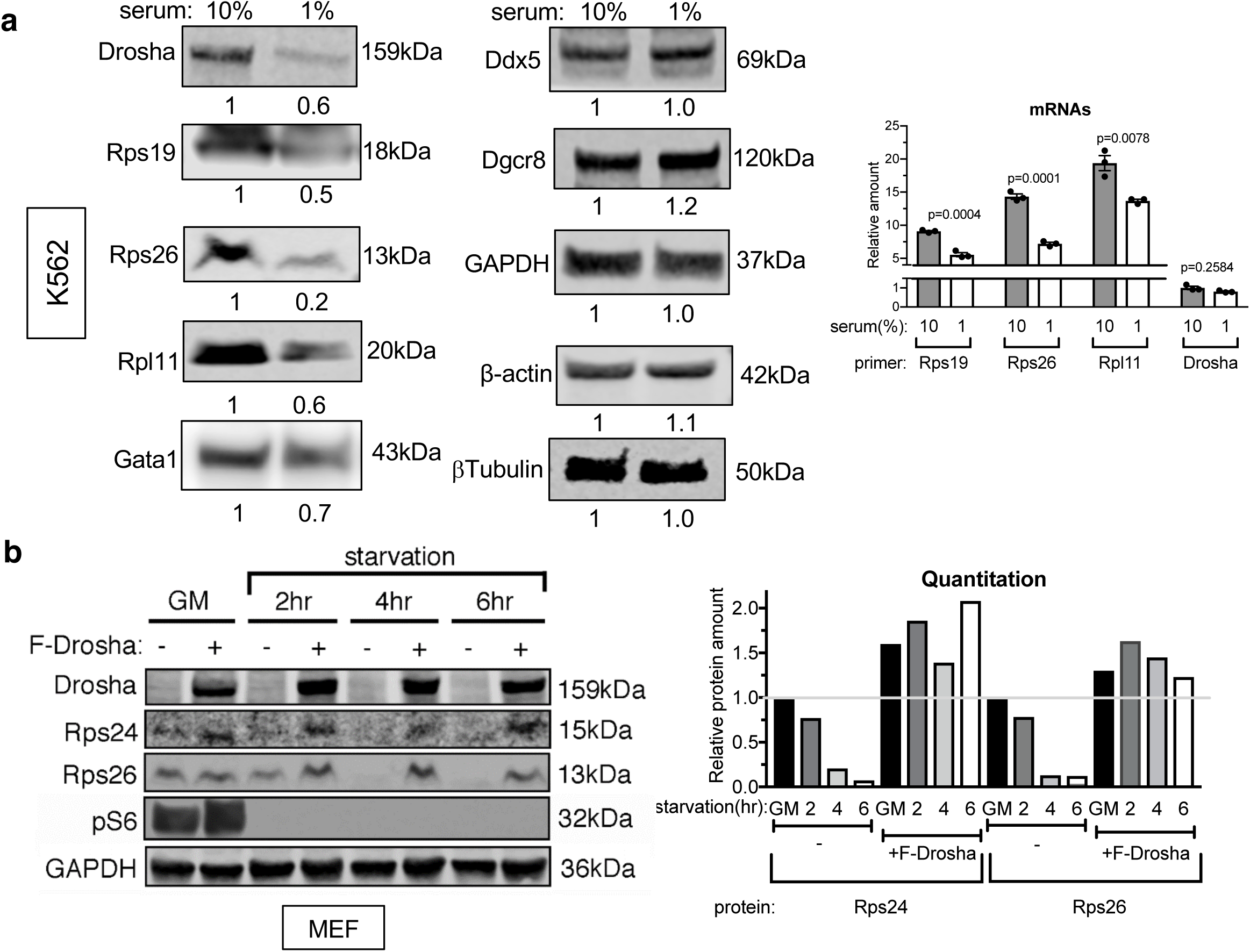
Nutrients deprivation decreases Drosha. **a.** Drosha, Rps19, Rps26, Rpl11, Dgcr8, Ddx5, Gata1, GAPDH (control), β-actin (control) β-Tubulin (control) proteins were examined by western blot in total cell lysates from K562 cells cultured in serum-starved (1% serum) or normal (10% serum) media for 6 hrs (left). Relative protein amount normalized to GAPDH is shown below each blot. qRT-PCR analysis of Rps19, Rps26, Rpl11, Dgcr8, Drosha mRNAs normalized to GAPDH (right). Mean ± SEM. **b.** The amount of Drosha, Rps26, Rps24, phospho-Rps6 (pS6), and GAPDH protein was examined by western blot in total cell lysates from Drosha-expressing or Ctrl MEFs cultured in growth media (GM, 10% serum) or starvation media (no serum) for a period of time as indicated (left). Relative amount of the protein normalized to GAPDH is plotted (right).

We then addressed the question of the effector of Drosha’s decrease upon starvation. After testing a small panel of candidate E3 ubiquitin ligases, we found that Nedd4 (Neural precursor cell expressed, developmentally downregulated 4, also known as Nedd4-1) is responsible for serum starvation-mediated degradation of Drosha. When Nedd4 was silenced by siRNA (siNedd4), the amount of Drosha as well as RPs in 1% serum remained as high as in 10% serum (**Fig. 7a, siNedd4**), demonstrating that Nedd4 is responsible for the degradation of Drosha. On the contrary, silencing of Nedd4L (also known as Nedd4-2)—a closely related member of the Nedd family of E3 ubiquitin ligases—did not rescue degradation of Drosha upon serum starvation (**Supplementary Fig. S9)**. We detected the interaction between Drosha and Nedd4 by immunoprecipitation of Flag-tagged Drosha (F-Drosha) by anti-Flag antibody, followed by immunoblot by anti-Nedd4 antibody (**Fig. 7b**). When Myc-tagged Ubiqutin (Ub) and Flag-Drosha were expressed in HEK293T cells, a small amount of poly-ubiquitinated Drosha (Ub-Drosha) was detected, which was greatly increased by overexpression of Nedd4 (**Fig. 7c**). Nedd4 contains WW domains that recognize PPxY (PY) motif on its substratate(Huang et al., 2019; Kanelis et al., 2006; Staub et al., 1996). We noted that human Drosha contains evolutionarily conserved PPGY sequence at amino acid 169-172, which is identical to the PY motif on one of Nedd4 substrates: Connexin 43(Leykauf et al., 2006; Spagnol et al., 2016). When PPGY sequence was mutated to AAGY in Drosha (AY mut), the level of Ub-Drosha was reduced, indicating that the PY motif is required for the ubiquitination by Nedd4 (**Fig. 7d**). When K562 cells were serum starved (in 1% FCS) for 6 h and 16 h, Nedd4 protein (**Fig. 7e**)— but not Nedd4 mRNA (**Supplementary Fig. S10**)—increased 1.6-fold and 2-fold, respectively, indicating a post-transcriptional induction of Nedd4 upon serum starvation. While Nedd4 increased, Drosha, Rps26, and Gata1 proteins were all decreased by serum starvation (**Fig. 7e**), as expected. Reduction of phosphorylated Rps6 (pS6) was detected after 16 h of serum starvation, suggesting an inhibition of the mTOR-p70 S6 kinase pathway (**Fig. 7e**). To examine the subcellular localization of Drosha and Nedd4 upon serum starvation, we performed a nuclear/cytoplasmic fractionation of K562 cells. pS6, Lamin A/C and β-Tubulin indicated a successful separation of the nuclear and cytoplasmic fractions (**Fig. 7f**). As expected, Drosha was predominantly localized in the nucleus under normal serum condition (**Fig. 7f, 0 hr**). Although the total amount of Drosha gradually declined upon serum starvation, the cytoplasmic fraction of Drosha was increased from 0.6% (0 h) to 33% (6 h) and 53% (16 h after serum starvation) (**Fig. 7f, bottom**). Unlike Drosha, Nedd4 was predominantly present in the cytoplasm, regardless of serum concentration (**Fig. 7f, bottom**). These results suggest that upon serum starvation, Drosha is exported from the nucleus and degraded by Nedd4 in the cytoplasm. Phosphorylation by p38 mitogen activated protein kinase (p38 MAPK) has been implicated in nuclear export and subsequent degradation of Drosha upon oxidative stress and heat(Yang et al., 2015). Inhibition of p38 MAPK by the specific inhibitor SB203580(Cuenda et al., 1995) partially inhibited Drosha nuclear export after serum starvation (**Fig. 7g**), suggesting that Drosha nuclear export is driven, in part, by p38 MAPK-dependent phosphorylation (**Fig. 7h**). Thus, we delineated the framework of a regulatory pathway connecting the abundance of serum in the growth medium to the export of Drosha from the nucleus (in part driven by p38 MAPK phosphorylation) into the cytoplasm, where Drosha is degraded through the action of Nedd4, which in turn is moderately increased by starvation. This mechanism involving the subcellular localization and protein stability of Drosha appears to play a key role in the control of ribosome abundance and global protein synthesis in response to a change in the growth stimulus.

**Figure 7.**
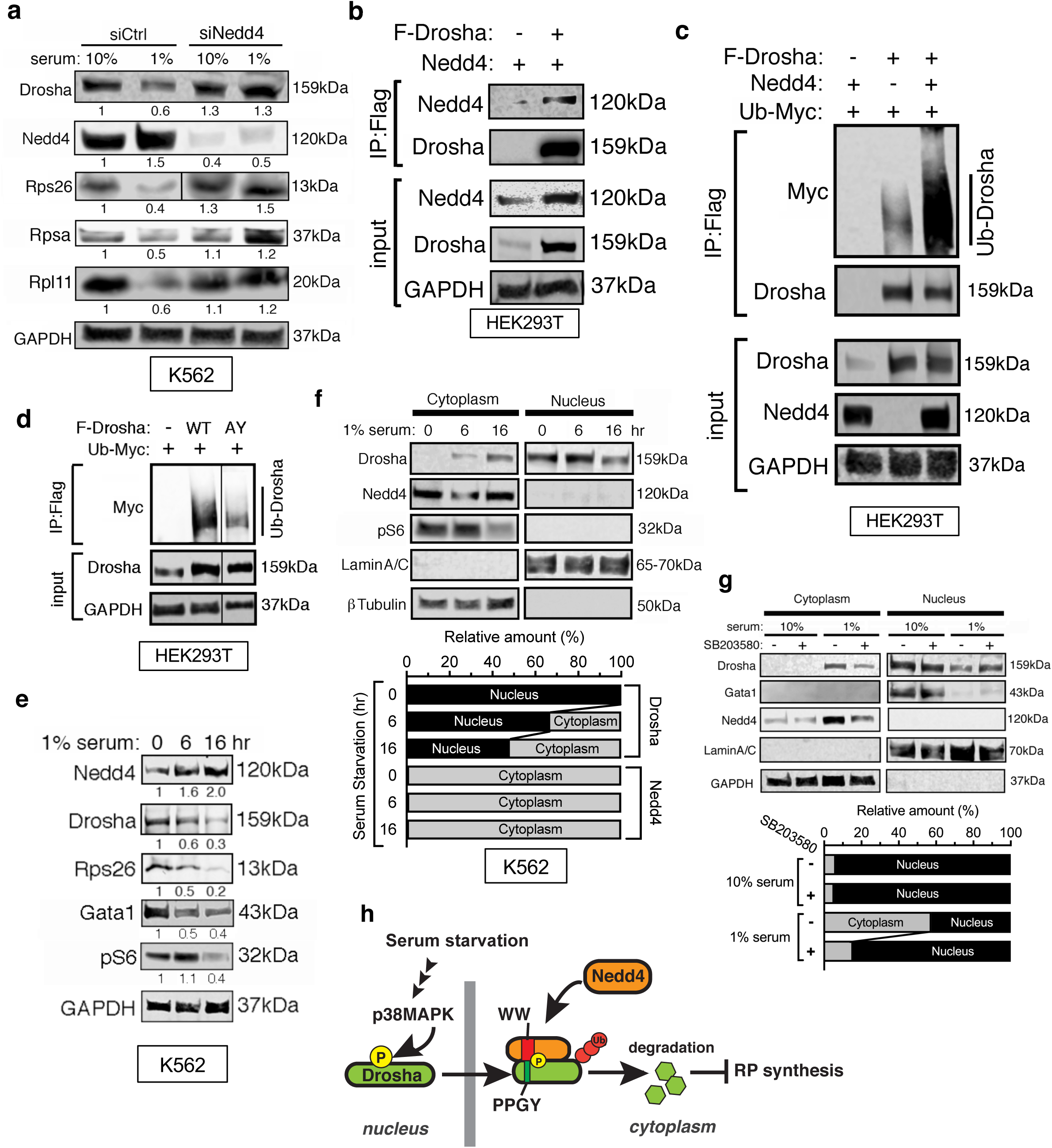
Serum starvation leads to the cytoplasmic shuttling and degradation of Drosha by Nedd4. **a.** Drosha, Nedd4, Rps26, Rpsa, Rpl11, and GAPDH protein was examined by western blot in total cell lysates from K562 cells transfected with non-specific control siRNA (siCtrl) or siRNA against Nedd4 (siNedd4) cultured in growth media (10% serum) or serum starvation media (1% serum) for 6 hrs. Right: Relative amount of the protein normalized to GAPDH is plotted. **b.** Association of Drosha with Nedd4 was examined by exogenously expressing Flag tagged Drosha (F-Drosha) and Nedd4 in HEK293T cells. F-Drosha was immunoprecipitated by anti-Flag (M2) antibody, followed by immunoblot by anti-Nedd4 and anti-Flag antibody. Total cell lysates (input) were immunoblotted with antibodies by Nedd4, Drosha, and GAPDH (loading control). **c**. Empty vector (pcDNA; mock) or Flag-tagged Drosha were expressed with Myc-tagged ubiquitin (Ub-Myc) in the presence or absence of Nedd4 in HEK293T cells. Total cell lysates were subjected to immunoprecipitation with anti-Flag antibody, followed by immunoblot with anti-Myc antibody (for Ub-Drosha) and anti-Drosha antibody. Input samples were subjected to immunoblot with anti-Drosha and anti-GAPDH (control) antibody. **d.** Flag-tagged Drosha (WT or AY mutant) and Ub-Myc were exogenously expressed in HEK293T cells. Total cell lysates were subjected to immunoprecipitation with anti-Flag antibody, followed by immunoblot with anti-Myc antibody for Ub-Drosha (top). Total cell lysates (input) were subjected to immunoblot with anti-Drosha and anti-GAPDH antibody (control). **e.** The amount of Nedd4, Drosha, Rps26, Gata1, phospho-Rps6 (pS6), and GAPDH (control) was examined by immunoblot in total cell lysates from K562 cells under serum starvation (1% serum) for 0, 6 or 16 hrs. Relative protein amount normalized to GAPDH is shown below below each blot. **f.** The cytoplasmic and the nuclear fraction were prepared from K562 cells treated with serum starvation (1% serum) for 0, 6, or 16 hrs, and subjected to immunoblot analysis of Drosha, Nedd4, phospho-Rps6 (pS6, control for cytoplasmic fraction), Lamin A/C (control for nuclear fraction) and bTubulin (control for cytoplasmic fraction) (top). Distribution of Drosha and Nedd4 in the cytoplasm vs nucleus (%) is quantitated (bottom). **g.** The cytoplasmic and the nuclear fraction were prepared from K562 cells were cultured in 10% or 1% serum containing media with vehicle (DMSO) or SB203580 (10 mM) for 16 hrs, harvested, and subjected to immunoblot analysis of Drosha, Gata1, Nedd4, Lamin A/C (control for nuclear fractions) and GAPDH (control for cytoplasmic fractions) (top). Relative amount of Drosha in the cytoplasm vs nucleus (%) is shown (bottom). **h.** Schematic mechanism of Nedd4-dependent regulation of Drosha upon serum starvation. Serum starvation promotes phosphorylation of Drosha by p38 MAPK in the nucleus, followed by translocation to the cytoplasm where the WW domain of Nedd4 ubiquitin ligase associates with the PPGY motif of Drosha, ubiquitinates and degrades Drosha. As a result, RP biosynthesis is inhibited.

## Discussion

Ribosome biogenesis is an energy-intensive process, hence it is crucial to coordinate the synthesis of RPs and rRNAs with the demand for protein synthesis determined by the cellular environment (Genuth and Barna, 2018). In this study, we found that the Microprocessor complex mediates the coordinated synthesis of RP mRNAs in response to changes in the cellular environment. Drosha was originally characterized as an enzyme involved in rRNA maturation, based on the observed accumulation of 45S and 32S pre-rRNAs upon Drosha knockdown (Oskowitz et al., 2011; Wu et al., 2000). We also found increased 45S pre-rRNA in Drosha-depleted erythrocyte progenitors compared to control cells (data not shown). Ddx5 has been involved in the processing of the 32S pre-rRNA (Jalal et al., 2007; Saporita et al., 2011). Thus, both Drosha and Ddx5 may control ribosome synthesis by coordinating RPs and rRNAs synthesis.

In *E. coli*, 54 RP genes are encoded in 20 operons and regulated by a translational feedback mechanism in which specific RPs directly bind to their own mRNAs and inhibit translation (Nomura et al., 1984; Zengel and Lindahl, 1994). In yeast, the RP genes, which lack the TOP motif, are regulated in large part by transcription factors, such as Ifh1, Fhl1, and Rap1 (Lieb et al., 2001; Martin et al., 2004; Rudra et al., 2005; Schawalder et al., 2004; Wade et al., 2004; Warner, 1999). A small number of yeast RPs, such as Rpl33p, Rps14p, and Rpl22p, bind their own precursor-mRNAs (pre-mRNA) and inhibit splicing to provide negative feedback regulation (Gabunilas and Chanfreau, 2016; Warner and McIntosh, 2009). In multicellular organisms, since it is the only known sequence element conserved among all RP genes, the 5’TOP motif is considered a key regulatory element for the coordinated control of RP synthesis. Previous studies suggest that the TOP motif provides control of both the transcription of TOP-genes(Parry et al., 2010; Shibui-Nihei et al., 2003) and the translation of TOP-mRNAs (Fonseca et al., 2018; Gentilella et al., 2017; Hamilton et al., 2006; Lahr et al., 2017; Meyuhas and Kahan, 2015; Patursky-Polischuk et al., 2014). A linear correlation of the activity of the luciferase reporters containing the *Rpl28* promoter and the abundance of their transcripts (**Fig. 4c)** suggests that the Microprocessor mainly contributes to the transcriptional control of RPs through the TOP motif, but we cannot exclude a contribution of the Microprocessor to the translational control of RP synthesis.

A TOP motif that is functionally equivalent to that in RPGs can also be found in non-RP genes(Meyuhas and Kahan, 2015). The eCLIP data show that Drosha and DGCR8 associate with non-ribosomal TOP-genes (**Supplementary Fig. S6**). The Quantitative mass-spectrometry analyses confirm that the expression of four non-ribosomal TOP-genes—*EIF4B*, *PABPC1*, and *VIM*—requires Drosha, similarly to RPs (**Supplementary Fig. S4**). Thus, the Microprocessor controls the synthesis of various components of the protein synthesis apparatus in addition to RPs(Meyuhas and Kahan, 2015).

Various molecules have been identified in association with the TOP sequence both on the DNA and the RNA, including DNA-binding [zinc finger protein 9 (ZFP9)](Avni et al., 1994; Huichalaf et al., 2009; Pellizzoni et al., 1997) and RNA-binding proteins [T cell intracellular antigen-1 (TIA-1) and TIA-1-related (TIAR)](Damgaard and Lykke-Andersen, 2011; Miloslavski et al., 2014), and microRNA-10a (Ørom et al., 2008). However, Drosha is the first protein to be described in association with the TOP motif of all RP transcripts, and as a factor able to synchronize RP synthesis to cell growth. The ssRNA binding protein La-related protein 1 (LARP1) was found to associate with several TOP-containing mRNAs, including a few RPG transcripts, and to repress translation in an mTOR-dependent manner(Fonseca et al., 2015; Hong et al., 2017; Lahr et al., 2017; Mura et al., 2015; Philippe et al., 2018; Tcherkezian et al., 2014). Although it may play an important role as a translational repressor, LARP1 binds ∼3,000 mRNAs, most of which do not contain a TOP motif(Mura et al., 2015), and is also implicated in the regulation of ATP production in mitochondria(To et al., 2019). Thus, it appears that LARP1 may exhibit a range of biological activities beyond the specific regulation of RP biosynthesis.

We have observed that nutrients deprivation promotes nuclear-to-cytoplasmic translocation of Drosha, followed by degradation by the cytoplasmic E3 ubiquitin ligase Nedd4. Nuclear-to-cytoplasmic shuttling of the Microprocessor complex and cleavage of viral RNAs upon viral infection has been reported as an antiviral mechanism, although the mechanism of regulation of the cytoplasmic shuttling of the Microprocessor upon viral infection is unknown(Shapiro et al., 2012; Shapiro et al., 2014). Our results suggest that p38 MAPK-dependent phosphorylation of Drosha at Ser^355^ contributes to the nuclear export of Drosha, as described in cells under oxidative or heat stress(Yang et al., 2015). Under these conditions of stress, cells globally reduce new protein synthesis, except for a small subset of proteins that are essential for the stress response(Duncan and Hershey, 1989; Reichmann et al., 2018). We propose that the p38 MAPK-Drosha-Nedd4 axis might be the mediator of several cellular stress stimuli, including starvation, oxidation and heat, which result in reduced protein synthesis. Unlike Ser^355^ phosphorylation by p38 MAPK, phosphorylation at Ser^300^ and Ser^302^ by Glycogen synthase kinase 3β (GSK3β) is required for the nuclear retention of Drosha(Tang et al., 2011). Because GSK3β activity is regulated by various extracellular signals(Beurel et al., 2015), it is possible that serum starvation might cause an inhibition of GSK3β and mediate dephosphorylation of Ser^300/302^ of Drosha, resulting in the accumulation in the cytoplasm. An alternatively spliced form of Drosha skipping exon 6, which encodes the putative nuclear localization signal, is reported to localize in both the cytoplasm and the nucleus(Dai et al., 2016; Link et al., 2016). However, the molecular size of cytoplasmic Drosha after serum starvation is equivalent to full-length Drosha (159 kDa) and indistinguishable from nuclear Drosha, thus it is unlikely that alternative splicing is involved in the change of nuclear/cytoplasmic ratio of Drosha upon serum starvation.

The transcriptional regulation of RP genes by Ddx5 through resolution of an R-loop is a new paradigm of regulation of ribosome biogenesis. We speculate that the presence of a GC-rich sequence downstream of the TOP motif in RP genes(Meyuhas and Kahan, 2015) may explain in part why R-loops formed at RPG loci are stable and capable of reducing the rate of transcription, unless they are actively destabilized by the Ddx5 helicase. It has been reported that the Microprocessor complex is recruited to DNA double strand break (DSB) sites, where it facilitates R-loop formation and promotes DSB repair (Bonath et al., 2018; Crossley et al., 2019; Lu et al., 2018). Thus, the Microprocessor may participate in both the formation and the resolution of R-loops, depending on the cellular context. Besides Ddx5, many RNA helicases—such as Dhx9, Ddx1, Ddx3x, Ddx15, Ddx17, Ddx18, Ddx21, Ddx27, Ddx39B, and Ddx54—are found in association with DNA/RNA hybrids (Cristini et al., 2018). For example, the Dhx9 helicase can either form or remove R-loops depending on the gene locus (Chakraborty and Grosse, 2011; Chakraborty et al., 2018; Cristini et al., 2018). Our study indicates that the TOP-Drosha axis allows the Ddx5 helicase to resolve R-loops specifically at RPG loci to promote RP transcription. Because there are 64 RNA helicases in human and 11 are found in association with the R-loops(Cristini et al., 2018), we speculate that other RNA helicases might be recruited to different gene loci, where they promote or resolve R-loops depending on context, and thus participate in gene regulation similarly to Ddx5 and Dhx9.

In yeast, only 280 genes contain introns, and nearly half of them are RP genes (Hooks et al., 2014). Yeast strains in which introns are deleted are viable in nutrient-rich media, but are unable to adapt to nutrient deprivation and die (Morgan et al., 2019; Parenteau et al., 2008; Parenteau et al., 2019). A subset of stable intronic RNAs accumulate during starvation and serve as mediators of the cellular response by either (1) limiting the efficiency or availability of the spliceosome or of other RNA-binding proteins (RBPs) and thus inhibiting RP mRNA splicing, and/or (2) by inhibiting target of rapamycin 1 (TORC1) activity (Morgan et al., 2019; Parenteau et al., 2008; Parenteau et al., 2019). We found no change in the unspliced form of RP mRNAs upon depletion of Drosha in mouse or human cells, indicating that Drosha does not control splicing of RP mRNAs (data not shown). Furthermore, nearly all genes in multicellular eukaryotes contain introns, and thus an intronic RNA-dependent regulatory mechanism would be neither specific nor sufficient to provide a robust control over RP biogenesis. We propose that the TOP-Microprocessor-dependent transcriptional regulation of RP genes evolved in multicellular organisms as an alternative to intronic RNAs-mediated regulation of RP mRNAs in yeast. Interestingly, both mechanisms involve a presumably ancient interaction between RNA and RBPs, whereas the abundance of RNA or RBPs is controlled by growth conditions.

Loss-of-function mutations in RPGs or in *Gata1* cause Diamond-Blackfan anemia (DBA; OMIM 105650)(Gripp et al., 2014; Ludwig et al., 2014; Ulirsch et al., 2018). In addition to anemia, DBA patients develop symptoms including short stature, arrhythmia, craniofacial defects, and thumb abnormalities—with various severity and penetrance(Ulirsch et al., 2018). The tissue-specific manifestation of physical abnormalities in DBA suggests a differential susceptibility to ribosome abnormalities or insufficiency of each cell type, presumably due to specific demands on protein synthesis. For example, we found that erythropoiesis is more severely impaired than vascular development in Drosha cKO mice (Jiang et al., 2017; Jiang et al., 2018). We speculate that erythrocyte progenitors, rapidly proliferating with high demand for Gata1 and globin protein synthesis, are more susceptible to ribosome shortages than endothelial cells. Currently, only ∼80% of DBA cases are accounted for by known gene mutations (Ulirsch et al., 2018). Our study opens the possibility that the remaining DBA patients with no known gene mutations might carry hypomorphic alleles of the components of the Microprocessor. Finally, altered ribosome biogenesis and translation is a hallmark of cancer cells. The signals that activate oncogenic signaling pathways, such as the Myc, Ras, and Akt pathways, and the loss of tumor suppressors, such as p53 and retinoblastoma (RB), lead to increased ribosome biogenesis(Sulima et al., 2017; Truitt and Ruggero, 2016). Frequent amplification of the *Drosha* locus correlates with a decreased survival rate in non-small cell lung carcinoma(Czubak et al., 2015). The *Ddx5* locus is also frequently amplified in breast carcinoma cells, which are “addicted” to Ddx5 for cell growth(Mazurek et al., 2012; Sulima et al., 2017). We speculate that elevated levels of Drosha and Ddx5 drive rapid cell proliferation and growth through overproduction of RPs and increased ribosome content in various tumors. Inhibiting the Ddx5 helicase could be a novel therapy to reduce tumor growth.

## Methods

### Animal care and use

All animal experiments were conducted in accordance with the guidelines of the Institutional Animal Care and Use Committee (IACUC) at University of California, San Francisco. *Cdh5-Cre* line (Chen et al., 2009) and *Drosha*^tm1Litt^ floxed line(Chong et al., 2008) have been previously described(Jiang et al., 2017). Embryos were dated by the presence of vaginal plug in the female mouse as embryonic day 0.5. The protocol number for the relevant animals and procedures approved by IACUC is AN170920-03: Title: Role of Growth Factor Signaling in Vascular Physiology” (Approval Date: August 20, 2019).

### Genotyping of mouse

Genomic DNAs were isolated from tail tips or conceptus yolk sacs of postnatal day 12 pups and genotyped with regular PCR. Primers for genotyping and RT-PCR are listed in Supplmentary Table S2.

### Flow Cytometry and cell sorting

Flow cytometry and cell sorting were performed as described previously^(Jiang et al., 2017)^. Briefly, fetal liver, yolk sac or AGM was dissected from embryos and mechanically dissociated by pipetting into single cell suspension in Hank’s Balanced Salt Solution containing 2% fetal bovine serum (FBS), 1% penicillin/streptomycin and buffered with 10 mM HEPES, pH7.2 (FACS buffer). E10.5 embryos were collected in 200 ul FACS buffer, followed by a collection of peripheral blood. Cells were stained with fluorochrome conjugated antibody at 4°C for 1 hr, washed with DAPI (0.5 µg ml−1) containing FACS buffer and analyzed by FACS Verse (BD

Biosciences) or sorted on a FACS Aria III (BD Biosciences) located at the UCSF FACS core. Data were analyzed with FlowJo v10.0.7. Single color stained samples were run with each flow cytometry analysis for compensation when analyzed with FlowJo. DAPI positive cells were gated out for analysis, then positive gating was applied when IgG staining sample was smaller than 0.1%.

### Antibodies

For Immunoblot, the following antibodies are used: Anti-Drosha antibody (1:500, Bethyl, A301-866A), anti-Dgcr8 (1:500, Proteintech,10996-1-AP), anti-Ddx5 (1:200, Abcam,ab21696), anti-GAPDH (1:5000, Millipore, MAB374), anti-Lamin A/C (1:2500, Cell signaling Technology, 2032), anti-Gata1 (1:200, R&D,MAB17791-SP), anti-γTubulin (1:5000, Santa Cruz Biotechnology, sc-7396), anti-puromycin (1:2000, Kerafast, 3RH11), anti-Rpl11 (1:300, Proteintech,16277-1-AP) anti-Rpsa (1:300, Abcam, ab137388), anti-Rps24 (1:300, Abcam, ab102986), anti-Rps26 (1:300, Abcam, ab104050), anti-Rps6 (1:500, Cell signaling Technology, 2317), anti-Rps19 (1:300, Santa Cruz Biotechnology, sc-100836), anti-β-actin (1:5000, Sigma-Aldrich, A5441), anti-NEDD4(1:2000, Cell signaling Technology, 2740), anti-Flag M2(1:2000, Sigma, F3165), anti-Myc (1:2500, Cell signaling Technology, 2278), anti-Myc-Tag (Cell signaling Technology, 2278), anti-RNA polymerase II (Millipore, 05-623), IRDye® 800CW Donkey anti-Rabbit IgG (H + L) (LI-COR, 926-32213), IRDye® 680RD Donkey anti-Mouse IgG (H + L) (LI-COR, 926-68072). For DRIP analysis, 10μg S9.6 (Millipore, MABE1095) antibody or normal mouse IgG (Santa Cruz, sc-2025) is used. For flow cytometry analysis or cell sorting, the following antibodies are used: PerCP anti-CD71 (1:200, BioLegend, 113815), FITC anti-Ter119 (1:200 BioLegend, 116205), PE anti-Itga4 (1:200, BioLegend,103607), Percp-IgG2bk (1:200, Biolegend 400336), FITC-IgG2ak (BD 553929), APC-IgG2bk (BD 556924), PE-IgG (Biolegend 405307), DAPI (ThermoFisher D1306).

### SDS-PAGE and immunoblot

SDS-PAGE and immunoblot were performed as described previously(Jiang et al., 2018).

### Cell culture and luciferase assay

K562 cells were cultured in 10%FBS in RPMI1640 media (Corning 10-040-CV) supplemented with 200μM L-glutamine, 100 μM sodium pyruvate and 1% penicillin/streptomycin at 37°C, 5% CO_2_. HCT116 cells, HEK293 or MEF cells were cultured in 10% FBS (Hyclone, SH3007103) in high glucose DMEM (Gibco 11965118) with 1% penicillin/streptomycin at 37°C, 5% CO_2_. The promoter area of human *Rpl28* gene was cloned at HindIII/NcoI site of the pGL3-basic (Promega, E1751) to make pGL3-reporters. pGL3-reporter, pGL3-basic (400 ng each), and renilla luciferase plasmid (1 ng) were transfected into HEK293 cells or HCT116 cells by lipofectamine2000 (Invitrogen, 11668030) following manufacturer’s manual. 48 h after transfection, total cell lysate was prepared and subjected to the luciferase assay as described(Chang et al., 2013). The firefly luciferase activity was normalized by the renilla luciferase activity to normalize transfection efficiency. For hemin treatment, K562 cells were treated with 25 μM hemin (Sigma-Aldrich, T8768) for 24 h, followed by benzidine ([1,1’-Biphenyl]-4,4’-diamine) staining as previously described(Rowley et al., 1985; Tomoda et al., 1991).

For serum starvation of K562 cells, 1% FBS was used to replace 10% FBS in regular culture medium. For serum starvation of MEF cells, HBSS (Thermo Fisher, 14025092) was used to replace the culture medium.

### Colony formation unit (CFU) assay

CFU assay was performed as described previously(Jiang et al., 2017). Briefly, E9.5 yolk sac was pipetted into single cell suspension which was seeded into methocult medium (Stem Cell Technologies M3434) and cultured for 7-10 days prior to counting the number of CFU-GM, BFU-E and CFU-GEMM colonies.

### Quantitative reverse transcriptase-polymerase chain reaction (RT-PCR) analysis

Five ng RNA from CD71^+^Itgα4^+^ (cKO) or CD71^+^Itgα4^-^ (Ctrl) cells sorted from peripheral blood of E10.5 embryos are amplified and reverse transcribed with a Nugen ovation picoSL WTA system V2 (Nugen 3312-24). Amplified cDNAs were 100-times diluted. RT-PCR reactions were then performed in triplicates using iQ SYBR Green supermix (Bio-RAD 1708882). Primer sequences are listed in Supplementary Table S2.

### Retrovirus, lentivirus production and infection

Lenti-crispr Drosha V2 (gRNA for Drosha) and lenti-crispr NS V2(none-specific gRNA as control) were obtained from Dr. Graveley (University of Conneticut). pLKO.1-shDDX5 (Broadinstitute, TRCN000000113) and pLKO.1-scramble were also obtained from Dr. Graveley and used for generating lentivirus shDDX5 or scramble sh RNA. 15 μg Lenti-crispr Drosha V2 or lenti-crispr NS V2, 7.5μg PMD2.G (Addgene plasmid #12259) and 7.5 μg psPAX2 (Addgene plasmid #12260) were transfected into HEK293T cells seeded in a 15cm dish at 70% confluence by lipofectamine2000 (Invitrogen, 11668030) following manufacturer’s manual. The media were replaced with DMED containing 10%FBS and high glucose 6 h after transfection. Lenti-virus supernatant was collected after 48 h and filtered with 0.45μm filter. Lenti-virus supernatant was aliquoted and stored at −80°C. Lenti-virus was added to HCT116 or HEK293T cells at 30% confluency. Lenti-virus containing media and culture media were mixed at 1:1 ratio. Polybrene was added to a final concentration of 8 μg/ml. Cell culture media were replaced with the lenti-virus and polybrene containing medium, grow in 5%CO_2_ at 37°C overnight. The media were replaced with regular culture media. 48 h after infection, puromycin was added to the media at a final concentration of 5 ng/μl to select the lenti-virus infected cells. For retrovirus production, twenty μg pBABE-Drosha or pBABE, 10μg pVSVG (Addgene plasmid #8454) and 10 μg psPAX2 (Addgene plasmid #12260) were transfected to HEK293T cells seeded in a 15cm dish at 70% confluency. After changing the media, supernatant was collected, filtered with 0.45μm filter, aliquoted and stored at −80°C. MEFs were infected with viral supernatant in 2 μg/ml polybrene, and select with 5ng/μl puromycin.

### Plasmid construction

Human Drosha cDNA with a Flag-tag at the amino-terminus was cloned into pBABE-puro vector (Addgene plasmid#1764) for producing retrovirus of human Drosha wild type (WT). Mouse Ddx5 WT(Addgene plasmid #88869) and Lys144 to Asn mutant(Addgene plasmid #88870) were obtained from Addgene(Huang et al., 2015). Ubiquitin_Myc_His plasmid is a gift from Dr. Jeff Wrana (<u>Lunenfeld-Tanenbaum Research Institute</u>, Mount Sinai Hospital, Toronto). WT or mutant promoter of human Rpl28 gene was cloned into pGL-3-basic (Promega, E1751). The number is relative to transcriptional start site, which is referred as +1. Drosha RNA binding domain (RBD, 1259aa-1337aa) was cloned into pCITE-2a (Novagen TB050) for in vitro transcription translation assay. pCI HA NEDD4(Addgene plasmid #27002) were obtained from Addgene. Human Drosha cDNA with a Flag-tag at the amino-terminus and a 6 x his tag at the carboxyl-terminus was cloned into pcDNA4/TO (Invitrogen) to construct the inducible WT Drosha expressing plasmid for immunoprecipitation assay and ubiquitination assay. The human Drosha Arg938 Lys939 Lys940 was mutated into Gln938 Ala939 Gln940 to construct a ribonuclease-defective Drosha expressing plasmid(Kwon et al., 2016).

### Electrophoretic mobility shift assay (EMSA)

100 pmol of WT or mutant RNA oligonucleotides (synthesized by Integrated DNA Technologies, Inc.) were 5’- end labeled with [γ^32^P] ATP (Perkin Elmer, NEG035C001MC) and T4 polynucleotide kinase (New England Biolabs) as previously described (Celona et al., 2017). Unincorporated ATP was removed by Illustra MicroSpin G-25 Columns (GE Healthcare Life Sciences, UK). The radiolabeled RNA probe was denatured in buffer (50 mM Tris-Cl ; 100 mM KCl; 2.5 mM MgCl2; 100 mM NaCl) at 72°C, and then renatured gradually at a rate of 1℃/min. EMSA was performed by incubating the radiolabeled probe (100,000 cpm) with Drosha RBD protein, which was synthesized in vitro with reticulocyte lysate systems (Promega, L5020), for 2 h at room temperature in binding buffer (50 mM Tris-Cl, pH 7.5; 100 mM KCl; 2.5 mM MgCl2; 100 mM NaCl; 0.01% NP-40; 1 mM DTT; 5% glycerol; 10 μg/ml bovine serum albumin; 0.1mg/ml sperm DNA) (Celona et al., 2017). The RNA-protein mixtures were then electrophoresed in 8% acrylamide-TBE gels. Gels were dried and exposed to X-ray film for analysis. For the competition experiments, the proteins were incubated with labeled WT RNA and 50-fold molar excess of the unlabeled WT or mutated single-stranded RNA at room temperature for 2 hours before electrophoresis.

### Chromatin immunoprecipitation (ChIP)

Flag-Drosha overexpressing MEFs and control MEFs (pBABE) were crosslinked treated with 2 mM disuccinimidyl glutarate (DSG; Thermo Fisher, 20593) at room temperature for 40 min. Remove the DSG, and wash with PBS. After MEFs were crosslinked with 1% Formaldehyde for 15 min at room temperature, followed by quenching with 1M Glycine, cells were washed with PBS and lysed with lysis buffer (50 mM Tris-Cl pH 8.1, 10 mM EDTA, 1% SDS and protease inhibitor). Genomic DNAs were sheared to average length of 200-500bp by sonication, followed by clearing lysates by centrifugation at 12,000g for 10 min at 4°C. Incubate the supernatant with protein A/G dynabeads (invitrogen 10002D) for 1 h at 4°C, dilute the pre-cleared sample to 1:10 ration with dilution buffer (20 mM Tris-Cl pH 8.1, 150 mM NaCl, 2 mM EDTA, 1% Triton X-100 and protease inhibitor) and 1/10 volume was kept as input before incubation with M2 dynabeads (Sigma-Aldrich, M8823) for 40 h at 4°C. After dynabeads were washed with a buffer I (20 mM Tris-Cl pH 8.1, 150 mM NaCl, 2 mM EDTA, 1% Triton X-100, 0.1% SDS), buffer II (20 mM Tris-Cl pH 8.1, 500 mM NaCl, 2 mM EDTA, 1% Triton X-100, 0.1% SDS), and buffer III (10mM Tris-Cl pH8.1, 250mM LiCl, 1mM EDTA, 1%NP-40, 1% Deoxycholate) at 4°C, the dynabeads were further washed twice with cold TE (10 mM Tris-Cl pH 8.1, 1 mM EDTA). The dynabeads were incubated in 250 μl elution buffer (200 mM NaHCO_3_, 1% SDS) at room temperature for 15 min twice. The eluates were mixed with 1/25 volume 5M NaCl and incubated at 65°C for 4 h. 1/50 volume of 0.5 M EDTA, 1/25 volume of Tris-Cl pH 6.5, 1/100 volume of Proteinase K (10 mg/ml) were added and incubated at 45°C for 1 hr. Precipitated genome fragments were purified with QIAquick PCR Purification Kit, followed by PCR analysis with primers in Supplementary Table S2. For Actinomycin D treatment, 1μg/ml Actinomycin D (Sigma-Aldrich, A9415) was applied in the culture medium for 30 min before ChIP assay. For RNase A treatment, RNaseA (Thermo Fisher, 12091021) was added into the lysed ChIP sample at a final concentration of 1μg/μl. For RNase H treatment, RNase H(New England Biolab M0523S) was added into the lysed ChIP sample at a final concentration of 100U/ml. Primer sequences are listed in Supplementary Table S2. For ChIP with anti-RNA polymerase II in K562 cells, crosslinking with DSG was omitted. Cells were crosslinked with 1% formaldehyde and all the other experimental procedures are the same as in MEFs.

### Immunoprecipitation assay

293T cells were transfected with pcDNA4-hDrosha and pCI HA NEDD4 or pCI empty vector. 2μg/ml doxycycline was added into medium to induce Drosha overexpression. 48 hours after transfection, cells were lysed in SBB buffer (1% Triton X-100, 150mM NaCl, 50mM Tris-Cl at pH 7.5, 1mM EDTA) supplemented with protease inhibitors (cOmplete, Roche-11836170001) and PhosStop (Roche-04906845001). Lysates were incubated at 4°C for 30 min, followed by centrifugation at 12,000 g for 10 min at 4°C. 1/10 of the lysate was saved as input sample for immunoblot. The lysate was incubated with anti-Flag M2 Magnetic Beads (sigma, M8823) and rocked overnight at 4°C. M2 beads were then washed in SBB buffer for 5 min at 4°C for three times, and boiled in sample buffer (Invitrogen, NP0007) with reducing agent (Invitrogen, NP0009). The elute and input are subjected to immunoblot analysis.

### DNA/RNA hybrid Immunoprecipitation (DRIP)

K562 cells were lysed in digestion buffer (100 mM NaCl, 10mM Tris-Cl at pH 8, 25mM EDTA at pH 8, 0.5% SDS, 0.65 mg/ml protease K) at 55°C overnight. 1 volume of phenol:chloroform:isoamyl alcohol (25:24:1, invitrogen, AM9722) was added into the lysate, mixed well, followed by centrifuge at 12000 x g for 10 minutes at 4°C. Supernatant was transferred to a new tube. 1 volume of phenol:chloroform:isoamyl alcohol was added again, followed by mixing and centrifuge. Transfer the supernatant and into new tube and precipitate with isopropanol. Take half of the genomic DNA (gDNA) for RNase H (New England Biolabs, M0297) treatment at 5 μl per 30 μg gDNA in 200 μl final volume for 24 h. After Rnase H treatment, the gDNA were subjected to restriction enzyme digestion: MseI, DdeI, AluI and MboI at 5U/50μl for each enzyme, shaking at 37°C for 24 h or longer until the gDNA are all shredded into fragments with length of 100-500 bp. The digested gDNA were then incubated with 10 μg S9.6 antibody (Millipore MABE1095) or mouse nonspecific IgG (Santa Cruz Biotechnology, sc-2025) in DRIP binding buffer (10mM NaPO4, PH7.0, 140mM NaCl, 0.05% Triton-X-100) at 4°C for overnight. Protein A was added to the sample and rotated at 4°C for 3 h followed by washing with 1X DRIP binding buffer for four times at room temperature. 0.5μg/μl Proteinase K was added to beads/antibody complexes and incubated for 40 min in a eppendorf ThermoMixer at 55°C, 1000 rpm. DNA was then extracted by cleaning up elution with same-volume phenol:chloroform:isoamyl alcohol, supernatant was moved to a new tube and precipitated with 1/10 volume 3M Sodium Acetate, 1μl glycogen (Invitrogen 10814010) and 2.5 volume ethanol. The DNA was then subjected to qPCR analysis, with primers listed in Supplementary Table S2.

### In vivo Ubiquitin Assay

pcDNA4-hDrosha, pCI HA NEDD4 or Ubiquitin_Myc_His plasmids were transfected into 293T cells at 30% confluence with lipofectamine 2000 (Invitrogen, 11668030) following the manufacture’s instructions. 2μg/ml doxycycline was added into medium to induce Drosha overexpression. 48 hours after transfection, cells of one 10 cm dish were lysed in 100 μl SBB buffer +1% SDS (1% Triton X-100, 150mM NaCl, 50mM Tris-Cl at pH 7.5, 1mM EDTA, 1% SDS) supplemented with protease inhibitors (cOmplete, Roche-11836170001) and PhosStop (Roche-04906845001), rocking at 4°C for half an hour. Each sample was then diluted with 900 μl SBB buffer and sonicated until cells were totally lysed. After sonication, cells were rocked at 4°C for 30-60 min and centrifuged at 15000 g for 20 min. Supernatant were transferred into a new tube, 100 μl was aliquoted as input, the other 900 μl were incubated with anti-Flag M2 Magnetic Beads (sigma, M8823) and rocked overnight at 4°C. M2 beads were then washed in SBB buffer for 5 min at 4°C for three times, and boiled in sample buffer (Invitrogen, NP0007) with reducing agent (Invitrogen, NP0009). The elute and input are subjected to westernblot analysis.

### Sucrose gradient fractionation of polysomes

Sucrose gradient fractionation of polysomes of K562 cells were performed as previously described(Floor and Doudna, 2016). Briefly, 2 T75 flasks of K562 cells (Ctrl and KO) were grown to a 2∼3X10^5^ cells/ml confluency in the growth media. 100μg/ml cycloheximide was added into the growth medium and incubated at 37°C for 5 min. The cells were then collected by centrifugation, and re-suspended PBS+100 in μg/ml cycloheximide. Cells were then pelleted again by centrifugation and lysed with three pellet-volumes ice cold hypotonic lysis buffer (10 mM HEPES pH 7.9, 1.5 mM MgCl_2_, 10 mM KCl, 0.5 mM DTT, 1% Triton X-100 and 100 μg/ml cycloheximide). After 10 min, cells were lysed on ice by ten strokes through a 26-gauge needle and nuclei were pelleted at 1,500× g for 5 min. Lysate from ∼15 million cells was layered on top of triplicate 10–50% (w/v) sucrose gradients (20 mM HEPES:KOH pH 7.6, 100 mM KCl, 5 mM MgCl_2_, 1 mM DTT and 100 μg/ml cycloheximide) made using a Biocomp Instruments (Canada) gradient master. Gradients were centrifuged for 2 h at 36,000 RPM in a SW-41 rotor, punctured, and manually peak fractionated using real-time A_260_ monitoring with a Brandel (Gaithersburg, MD) gradient fractionator and ISCO (Lincoln, NE) UA-6 detector. Fractions were subjected to RNA prep with RNeasy Plus micro kit (Qiagen 74034) and a RT-PCR analysis with SYBR green (Biorad, 1725120).

### Quantitative mass spectrometry

K562 cells were infected with none specific or Drosha CRISPR lenti-virus as described above with polybrene at a final concentration of 8μg/ml. 48 hours after infection, K562 cells were selected with 2.5 ng/μl puromycin. Five replicates of each genotype K562 cells were collected, washed with PBS, flash-frozen with liquid nitrogen. iTRAQ-TMT mass spectrometry was performed using Q Exactive mass spectrometer (Thermo Fisher Scientific, USA) by Creative Proteomics, Inc. Ten million cells were collected for each replicate. 5 replicates per sample. The 6 raw MS files were analyzed and searched against human protein database based on the species of the samples using Maxquant (1.6.2.6). The parameters were set as follows: the protein modifications were carbamidomethylation (C) (fixed), oxidation (M) (variable); the enzyme specificity was set to trypsin; the maximum missed cleavages were set to 2; the precursor ion mass tolerance was set to 10 ppm, and MS/MS tolerance was 0.6 Da. Only high confident identified peptides were chosen for downstream protein identification analysis.

### Next Generation RNA-sequencing and high-throughput data analysis

Erythroblast cells (CD71^high^Ter119^+^) from E10.5 PB of five Ctrl embryos with genotype of *Drosha*^fl/+^; *Cdh5- cre*^+^ or *Drosha*^fl/fl^, or Drosha^fl/+^ and five cKO embryos with genotype of *Drosha*^fl/fl^; *Cdh5-cre*^+^ were sorted on a FACS Aria III (BD Biosciences) located at the UCSF FACS core and subjected to RNA preperation with RNeasy Plus micro kit (Qiagen 74034). The quality of RNAs was evaluated with 2100 Bioanalyzer Instrument (Agilent Technologies). RNA samples with RIN>8.0 were shipped to Beijin Genome Institute for a library preparation and sequencing (Illumina HiSeq 2500). Integrative Genomics Viewer (2.4.14 Broad Institute) was used for data analysis.

Fastq files for DRIP-seq were downloaded from GEO datasets (GSE97648) and aligned to the human genome(GRCh38) using bowtie2-2.3.4.1.

### Puromycin incorporation assay

For in vivo puromycin incorporation, puromycin (0.04 μmol/g body weight) was injected into E11.5 pregnant mice intraperitoneally 35 min before embryos were harvested. Erythroid progenitors (Ter119^+/^CD71^+^) were isolated by flow cytometry from the peripheral blood of embryos, homogenized, and prepared for immunoblotting with anti-puromycin antibody. For in vitro puromycin incorporation in K562 cells, after serum starvation for 16 h, 1 μmol/l puromycin was added to the culture media for 10 min. Cells were then harvested, total cell lysates were generated from 5X10^6^ cells from both 10% or 1% serum treated K562 cells, and subjected to immunoblotting with anti-puromycin antibody (Kerafast, #EQ0001).

### Statistical analysis

Graphs were generated with GraphPad PRISM software. Statistical significance was calculated in R version 3.2.3 by Student’s t test. The null hypothesis of the medians/means being equal was rejected at α = 0.05 and p values were generated by unpaired Student’s t test and presented in figures. The sample size was estimated by power analysis and is presented in the figure legend. The investigators were blinded during experiments because genotyping was performed after experiments. No animals were excluded, and animals were allocated based on genotype. Cells for experiments were randomized. For animal analysis, at least three animals were used in each experiment and all experiments were completed in gender- and genotype-blinded manner. All the other experiments were performed at least three times with biological triplicates each time. For PCR analysis, each biological sample was analyzed with a specific primer set in triplicates each time.

### Data availability

RNA sequencing data were submitted to SRA-NCBI database and currently waiting for an accession number. In a mean time, we will make data available to the reviewers upon request.

## Supporting information

Supplemental Figurse and Tables

## Acknowledgements

We thank Kyle Soo, Bryn Sullivan, Armita Norouzi for contributing to the research. We thank Dr. Brenton Graveley (Univ. of Connecticut) for Lenti-crispr-Drosha and -NS construct, and Dr. Narry, V. Kim (Seoul National Univ.) for Drosha RKK/QAQ mutant construct. This study was funded by NIH HL132058 to A.H, DK119621 to B.L.B., the UCSF Program for Breakthrough Biomedical Research, funded in part by the Sandler Foundation to S.N.F, the California Tobacco-Related Disease Research Grants Program 27KT-0003 to S.N.F, and the National Institutes of Health DP2GM132932 to S.N.F. X.J. was supported by NIH T32 training grant to B.L.B. The authors have no conflicts of interest.

## Author Contributions

J.X., A.P., S. VDV, P.G., L.C., B.C., C.L., W.W. and S.V. designed the experiments and conducted the experiments. B.L.B., S.N.F., G.L, and A.H. designed the experiments and wrote the paper. All authors reviewed and approved the manuscript.

## References

Aplan, P.D., Nakahara, K., Orkin, S.H., and Kirsch, I.R. (1992). The SCL gene product: a positive regulator of erythroid differentiation. EMBO J 11, 4073–4081.

Aviner, R., Geiger, T., and Elroy-Stein, O. (2013). Novel proteomic approach (PUNCH-P) reveals cell cycle-specific fluctuations in mRNA translation. Genes Dev 27, 1834–1844.

Avni, D., Shama, S., Loreni, F., and Meyuhas, O. (1994). Vertebrate mRNAs with a 5’-terminal pyrimidine tract are candidates for translational repression in quiescent cells: characterization of the translational cis-regulatory element. Mol Cell Biol 14, 3822–3833.

Beurel, E., Grieco, S.F., and Jope, R.S. (2015). Glycogen synthase kinase-3 (GSK3): regulation, actions, and diseases. Pharmacol Ther 148, 114–131.

Bonath, F., Domingo-Prim, J., Tarbier, M., Friedländer, M.R., and Visa, N. (2018). Next-generation sequencing reveals two populations of damage-induced small RNAs at endogenous DNA double-strand breaks. Nucleic Acids Res 46, 11869–11882.

Boucas, J. (2018). Integration of ENCODE RNAseq and eCLIP Data Sets. Methods in molecular biology (Clifton, NJ 1720, 111–129.

Byrne, M.E. (2009). A role for the ribosome in development. Trends Plant Sci 14, 512–519.

Campbell, A.E., Wilkinson-White, L., Mackay, J.P., Matthews, J.M., and Blobel, G.A. (2013). Analysis of disease-causing GATA1 mutations in murine gene complementation systems. Blood 121, 5218–5227.

Celona, B., Dollen, J.V., Vatsavayai, S.C., Kashima, R., Johnson, J.R., Tang, A.A., Hata, A., Miller, B.L., Huang, E.J., Krogan, N.J., et al. (2017). Suppression of C9orf72 RNA repeat-induced neurotoxicity by the ALS-associated RNA-binding protein Zfp106. Elife 6.

Chakraborty, P., and Grosse, F. (2011). Human DHX9 helicase preferentially unwinds RNA-containing displacement loops (R-loops) and G-quadruplexes. DNA Repair (Amst) 10, 654–665.

Chakraborty, P., Huang, J.T.J., and Hiom, K. (2018). DHX9 helicase promotes R-loop formation in cells with impaired RNA splicing. Nat Commun 9, 4346.

Chang, J., Davis-Dusenbery, B.N., Kashima, R., Jiang, X., Marathe, N., Sessa, R., Louie, J., Gu, W., Lagna, G., and Hata, A. (2013). Acetylation of p53 stimulates miRNA processing and determines cell survival following genotoxic stress. EMBO J 32, 3192–3205.

Chen, M.J., Yokomizo, T., Zeigler, B.M., Dzierzak, E., and Speck, N.A. (2009). Runx1 is required for the endothelial to haematopoietic cell transition but not thereafter. Nature 457, 887–891.

Chen, X., Wang, L., Huang, R., Qiu, H., Wang, P., Wu, D., Zhu, Y., Ming, J., Wang, Y., Wang, J., et al. (2019). Dgcr8 deletion in the primitive heart uncovered novel microRNA regulating the balance of cardiac-vascular gene program. Protein Cell 10, 327–346.

Chong, M.M., Rasmussen, J.P., Rudensky, A.Y., and Littman, D.R. (2008). The RNAseIII enzyme Drosha is critical in T cells for preventing lethal inflammatory disease. J Exp Med 205, 2005–2017.

Coghill, E., Eccleston, S., Fox, V., Cerruti, L., Brown, C., Cunningham, J., Jane, S., and Perkins, A. (2001). Erythroid Kruppel-like factor (EKLF) coordinates erythroid cell proliferation and hemoglobinization in cell lines derived from EKLF null mice. Blood 97, 1861–1868.

Consortium, E.P. (2012). An integrated encyclopedia of DNA elements in the human genome. Nature 489, 57–74.

Cristini, A., Groh, M., Kristiansen, M.S., and Gromak, N. (2018). RNA/DNA Hybrid Interactome Identifies DXH9 as a Molecular Player in Transcriptional Termination and R-Loop-Associated DNA Damage. Cell Rep 23, 1891–1905.

Crossley, M.P., Bocek, M., and Cimprich, K.A. (2019). R-Loops as Cellular Regulators and Genomic Threats. Mol Cell 73, 398–411.

Cuenda, A., Rouse, J., Doza, Y.N., Meier, R., Cohen, P., Gallagher, T.F., Young, P.R., and Lee, J.C. (1995). SB 203580 is a specific inhibitor of a MAP kinase homologue which is stimulated by cellular stresses and interleukin-1. FEBS Lett 364, 229–233.

Czubak, K., Lewandowska, M.A., Klonowska, K., Roszkowski, K., Kowalewski, J., Figlerowicz, M., and Kozlowski, P. (2015). High copy number variation of cancer-related microRNA genes and frequent amplification of DICER1 and DROSHA in lung cancer. Oncotarget 6, 23399–23416.

Dai, L., Chen, K., Youngren, B., Kulina, J., Yang, A., Guo, Z., Li, J., Yu, P., and Gu, S. (2016). Cytoplasmic Drosha activity generated by alternative splicing. Nucleic Acids Res 44, 10454–10466.

Damgaard, C.K., and Lykke-Andersen, J. (2011). Translational coregulation of 5’TOP mRNAs by TIA-1 and TIAR. Genes Dev 25, 2057–2068.

Davis, C.A., Hitz, B.C., Sloan, C.A., Chan, E.T., Davidson, J.M., Gabdank, I., Hilton, J.A., Jain, K., Baymuradov, U.K., Narayanan, A.K., et al. (2018). The Encyclopedia of DNA elements (ENCODE): data portal update. Nucleic Acids Res 46, D794–D801.

Duncan, R.F., and Hershey, J.W. (1989). Protein synthesis and protein phosphorylation during heat stress, recovery, and adaptation. J Cell Biol 109, 1467–1481.

Floor, S.N., and Doudna, J.A. (2016). Tunable protein synthesis by transcript isoforms in human cells. Elife 5.

Fonseca, B.D., Lahr, R.M., Damgaard, C.K., Alain, T., and Berman, A.J. (2018). LARP1 on TOP of ribosome production. Wiley interdisciplinary reviews RNA, e1480.

Fonseca, B.D., Zakaria, C., Jia, J.J., Graber, T.E., Svitkin, Y., Tahmasebi, S., Healy, D., Hoang, H.D., Jensen, J.M., Diao, I.T., et al. (2015). La-related Protein 1 (LARP1) Represses Terminal Oligopyrimidine (TOP) mRNA Translation Downstream of mTOR Complex 1 (mTORC1). J Biol Chem 290, 15996–16020.

Gabunilas, J., and Chanfreau, G. (2016). Splicing-Mediated Autoregulation Modulates Rpl22p Expression in Saccharomyces cerevisiae. PLoS Genet 12, e1005999.

García-Muse, T., and Aguilera, A. (2019). R Loops: From Physiological to Pathological Roles. Cell 179, 604–618.

Gentilella, A., Morón-Duran, F.D., Fuentes, P., Zweig-Rocha, G., Riaño-Canalias, F., Pelletier, J., Ruiz, M., Turón, G., Castaño, J., Tauler, A., et al. (2017). Autogenous Control of 5′TOP mRNA Stability by 40S Ribosomes. Mol Cell 67, 55–70.e54.

Genuth, N.R., and Barna, M. (2018). Heterogeneity and specialized functions of translation machinery: from genes to organisms. Nat Rev Genet 19, 431–452.

Ginno, P.A., Lim, Y.W., Lott, P.L., Korf, I., and Chédin, F. (2013). GC skew at the 5’ and 3’ ends of human genes links R-loop formation to epigenetic regulation and transcription termination. Genome Res 23, 1590–1600.

Gowrishankar, J., Leela, J.K., and Anupama, K. (2013). R-loops in bacterial transcription: their causes and consequences. Transcription 4, 153–157.

Green, A.R., DeLuca, E., and Begley, C.G. (1991). Antisense SCL suppresses self-renewal and enhances spontaneous erythroid differentiation of the human leukaemic cell line K562. EMBO J 10, 4153–4158.

Gripp, K.W., Curry, C., Olney, A.H., Sandoval, C., Fisher, J., Chong, J.X., Pilchman, L., Sahraoui, R., Stabley, D.L., Sol-Church, K., et al. (2014). Diamond-Blackfan anemia with mandibulofacial dystostosis is heterogeneous, including the novel DBA genes TSR2 and RPS28. Am J Med Genet A 164A, 2240–2249.

Groh, M., and Gromak, N. (2014). Out of balance: R-loops in human disease. PLoS Genet 10, e1004630.

Gromak, N., Dienstbier, M., Macias, S., Plass, M., Eyras, E., Cáceres, J.F., and Proudfoot, N.J. (2014). Drosha Regulates Gene Expression Independently of RNA Cleavage Function. Cell Rep 7, 1753–1754.

Hafner, J., Haenseler, E., Ossent, P., Burg, G., and Panizzon, R.G. (1995). Benzidine stain for the histochemical detection of hemoglobin in splinter hemorrhage (subungual hematoma) and black heel. Am J Dermatopathol 17, 362–367.

Hamilton, T.L., Stoneley, M., Spriggs, K.A., and Bushell, M. (2006). TOPs and their regulation. Biochem Soc Trans 34, 12–16.

Han, J., Lee, Y., Yeom, K.H., Kim, Y.K., Jin, H., and Kim, V.N. (2004). The Drosha-DGCR8 complex in primary microRNA processing. Genes Dev 18, 3016–3027.

Hata, A., and Lieberman, J. (2015). Dysregulation of microRNA biogenesis and gene silencing in cancer. Sci Signal 8, re3.

Hong, S., Freeberg, M.A., Han, T., Kamath, A., Yao, Y., Fukuda, T., Suzuki, T., Kim, J.K., and Inoki, K. (2017). LARP1 functions as a molecular switch for mTORC1-mediated translation of an essential class of mRNAs. Elife 6.

Hooks, K.B., Delneri, D., and Griffiths-Jones, S. (2014). Intron evolution in Saccharomycetaceae. Genome Biol Evol 6, 2543–2556.

Huang, W., Thomas, B., Flynn, R.A., Gavzy, S.J., Wu, L., Kim, S.V., Hall, J.A., Miraldi, E.R., Ng, C.P., Rigo, F., et al. (2015). DDX5 and its associated lncRNA Rmrp modulate TH17 cell effector functions. Nature 528, 517–522.

Huang, X., Chen, J., Cao, W., Yang, L., Chen, Q., He, J., Yi, Q., Huang, H., Zhang, E., and Cai, Z. (2019). The many substrates and functions of NEDD4-1. Cell Death Dis 10, 904.

Huertas, P., and Aguilera, A. (2003). Cotranscriptionally formed DNA:RNA hybrids mediate transcription elongation impairment and transcription-associated recombination. Mol Cell 12, 711–721.

Huichalaf, C., Schoser, B., Schneider-Gold, C., Jin, B., Sarkar, P., and Timchenko, L. (2009). Reduction of the rate of protein translation in patients with myotonic dystrophy 2. J Neurosci 29, 9042–9049.

Jalal, C., Uhlmann-Schiffler, H., and Stahl, H. (2007). Redundant role of DEAD box proteins p68 (Ddx5) and p72/p82 (Ddx17) in ribosome biogenesis and cell proliferation. Nucleic Acids Res 35, 3590–3601.

Jiang, X., Hawkins, J.S., Lee, J., Lizama, C.O., Bos, F.L., Zape, J.P., Ghatpande, P., Peng, Y., Louie, J., Lagna, G., et al. (2017). Let-7 microRNA-dependent control of leukotriene signaling regulates the transition of hematopoietic niche in mice. Nat Commun 8, 128.

Jiang, X., Wooderchak-Donahue, W.L., McDonald, J., Ghatpande, P., Baalbaki, M., Sandoval, M., Hart, D., Clay, H., Coughlin, S., Lagna, G., et al. (2018). Inactivating mutations in Drosha mediate vascular abnormalities similar to hereditary hemorrhagic telangiectasia. Sci Signal 11.

Kanelis, V., Bruce, M.C., Skrynnikov, N.R., Rotin, D., and Forman-Kay, J.D. (2006). Structural determinants for high-affinity binding in a Nedd4 WW3* domain-Comm PY motif complex. Structure 14, 543–553.

Kim, B., Jeong, K., and Kim, V.N. (2017). Genome-wide Mapping of DROSHA Cleavage Sites on Primary MicroRNAs and Noncanonical Substrates. Mol Cell 66, 258–269.e255.

Kim, Y.K., Kim, B., and Kim, V.N. (2016). Re-evaluation of the roles of DROSHA, Export in 5, and DICER in microRNA biogenesis. Proc Natl Acad Sci U S A 113, E1881–1889.

Koulnis, M., Pop, R., Porpiglia, E., Shearstone, J.R., Hidalgo, D., and Socolovsky, M. (2011). Identification and analysis of mouse erythroid progenitors using the CD71/TER119 flow-cytometric assay. J Vis Exp.

Kwon, S.C., Nguyen, T.A., Choi, Y.G., Jo, M.H., Hohng, S., Kim, V.N., and Woo, J.S. (2016). Structure of Human DROSHA. Cell 164, 81–90.

Lahr, R.M., Fonseca, B.D., Ciotti, G.E., Al-Ashtal, H.A., Jia, J.J., Niklaus, M.R., Blagden, S.P., Alain, T., and Berman, A.J. (2017). La-related protein 1 (LARP1) binds the mRNA cap, blocking eIF4F assembly on TOP mRNAs. Elife 6.

Lee, D., and Shin, C. (2018). Emerging roles of DROSHA beyond primary microRNA processing. RNA Biol 15, 186–193.

Leppek, K., Das, R., and Barna, M. (2018). Functional 5’ UTR mRNA structures in eukaryotic translation regulation and how to find them. Nat Rev Mol Cell Biol 19, 158–174.

Leykauf, K., Salek, M., Bomke, J., Frech, M., Lehmann, W.D., Durst, M., and Alonso, A. (2006). Ubiquitin protein ligase Nedd4 binds to connexin43 by a phosphorylation-modulated process. J Cell Sci 119, 3634–3642.

Lieb, J.D., Liu, X., Botstein, D., and Brown, P.O. (2001). Promoter-specific binding of Rap1 revealed by genome-wide maps of protein-DNA association. Nat Genet 28, 327–334.

Link, S., Grund, S.E., and Diederichs, S. (2016). Alternative splicing affects the subcellular localization of Drosha. Nucleic Acids Res 44, 5330–5343.

Lopez-Carballo, G., Moreno, L., Masia, S., Perez, P., and Barettino, D. (2002). Activation of the phosphatidylinositol 3-kinase/Akt signaling pathway by retinoic acid is required for neural differentiation of SH-SY5Y human neuroblastoma cells. J Biol Chem 277, 25297–25304.

Lu, W.T., Hawley, B.R., Skalka, G.L., Baldock, R.A., Smith, E.M., Bader, A.S., Malewicz, M., Watts, F.Z., Wilczynska, A., and Bushell, M. (2018). Drosha drives the formation of DNA:RNA hybrids around DNA break sites to facilitate DNA repair. Nat Commun 9, 532.

Ludwig, L.S., Gazda, H.T., Eng, J.C., Eichhorn, S.W., Thiru, P., Ghazvinian, R., George, T.I., Gotlib, J.R., Beggs, A.H., Sieff, C.A., et al. (2014). Altered translation of GATA1 in Diamond-Blackfan anemia. Nat Med 20, 748–753.

Martin, D.E., Soulard, A., and Hall, M.N. (2004). TOR regulates ribosomal protein gene expression via PKA and the Forkhead transcription factor FHL1. Cell 119, 969–979.

Mazurek, A., Luo, W., Krasnitz, A., Hicks, J., Powers, R.S., and Stillman, B. (2012). DDX5 regulates DNA replication and is required for cell proliferation in a subset of breast cancer cells. Cancer Discov 2, 812–825.

Mersaoui, S.Y., Yu, Z., Coulombe, Y., Karam, M., Busatto, F.F., Masson, J.Y., and Richard, S. (2019). Arginine methylation of the DDX5 helicase RGG/RG motif by PRMT5 regulates resolution of RNA:DNA hybrids. EMBO J 38, e100986.

Meyuhas, O., and Kahan, T. (2015). The race to decipher the top secrets of TOP mRNAs. Biochim Biophys Acta 1849, 801–811.

Miloslavski, R., Cohen, E., Avraham, A., Iluz, Y., Hayouka, Z., Kasir, J., Mudhasani, R., Jones, S.N., Cybulski, N., Rüegg, M.A., et al. (2014). Oxygen sufficiency controls TOP mRNA translation via the TSC-Rheb-mTOR pathway in a 4E-BP-independent manner. Journal of molecular cell biology 6, 255–266.

Morgan, J.T., Fink, G.R., and Bartel, D.P. (2019). Excised linear introns regulate growth in yeast. Nature 565, 606–611.

Mura, M., Hopkins, T.G., Michael, T., Abd-Latip, N., Weir, J., Aboagye, E., Mauri, F., Jameson, C., Sturge, J., Gabra, H., et al. (2015). LARP1 post-transcriptionally regulates mTOR and contributes to cancer progression. Oncogene 34, 5025–5036.

Nomura, M., Gourse, R., and Baughman, G. (1984). Regulation of the synthesis of ribosomes and ribosomal components. Annual review of biochemistry 53, 75–117.

Nudler, E. (2012). RNA polymerase backtracking in gene regulation and genome instability. Cell 149, 1438–1445.

Ørom, U.A., Nielsen, F.C., and Lund, A.H. (2008). MicroRNA-10a binds the 5’UTR of ribosomal protein mRNAs and enhances their translation. Mol Cell 30, 460–471.

Oskowitz, A.Z., Penfornis, P., Tucker, A., Prockop, D.J., and Pochampally, R. (2011). Drosha regulates hMSCs cell cycle progression through a miRNA independent mechanism. Int J Biochem Cell Biol 43, 1563–1572.

Pakos-Zebrucka, K., Koryga, I., Mnich, K., Ljujic, M., Samali, A., and Gorman, A.M. (2016). The integrated stress response. EMBO Rep 17, 1374–1395.

Parenteau, J., Durand, M., Véronneau, S., Lacombe, A.A., Morin, G., Guérin, V., Cecez, B., Gervais-Bird, J., Koh, C.S., Brunelle, D., et al. (2008). Deletion of many yeast introns reveals a minority of genes that require splicing for function. Mol Biol Cell 19, 1932–1941.

Parenteau, J., Maignon, L., Berthoumieux, M., Catala, M., Gagnon, V., and Abou Elela, S. (2019). Introns are mediators of cell response to starvation. Nature 565, 612–617.

Park, Y.S., Liu, Z., Vasamsetti, B.M., and Cho, N.J. (2016). The ERK1/2 and mTORC1 Signaling Pathways Are Involved in the Muscarinic Acetylcholine Receptor-Mediated Proliferation of SNU-407 Colon Cancer Cells. Journal of cellular biochemistry 117, 2854–2863.

Parry, T.J., Theisen, J.W., Hsu, J.Y., Wang, Y.L., Corcoran, D.L., Eustice, M., Ohler, U., and Kadonaga, J.T. (2010). The TCT motif, a key component of an RNA polymerase II transcription system for the translational machinery. Genes Dev 24, 2013–2018.

Patrick, D.M., Zhang, C.C., Tao, Y., Yao, H., Qi, X., Schwartz, R.J., Jun-Shen Huang, L., and Olson, E.N. (2010). Defective erythroid differentiation in miR-451 mutant mice mediated by 14-3-3zeta. Genes Dev 24, 1614–1619.

Patursky-Polischuk, I., Kasir, J., Miloslavski, R., Hayouka, Z., Hausner-Hanochi, M., Stolovich-Rain, M., Tsukerman, P., Biton, M., Mudhasani, R., Jones, S.N., et al. (2014). Reassessment of the role of TSC, mTORC1 and microRNAs in amino acids-meditated translational control of TOP mRNAs. PLoS One 9, e109410.

Pellizzoni, L., Lotti, F., Maras, B., and Pierandrei-Amaldi, P. (1997). Cellular nucleic acid binding protein binds a conserved region of the 5’ UTR of Xenopus laevis ribosomal protein mRNAs. J Mol Biol 267, 264–275.

Perina, D., Korolija, M., Roller, M., Harcet, M., Jeličić, B., Mikoč, A., and Cetković, H. (2011). Over-represented localized sequence motifs in ribosomal protein gene promoters of basal metazoans. Genomics 98, 56–63.

Pevny, L., Lin, C.S., D’Agati, V., Simon, M.C., Orkin, S.H., and Costantini, F. (1995). Development of hematopoietic cells lacking transcription factor GATA-1. Development 121, 163–172.

Pevny, L., Simon, M.C., Robertson, E., Klein, W.H., Tsai, S.F., D’Agati, V., Orkin, S.H., and Costantini, F. (1991). Erythroid differentiation in chimaeric mice blocked by a targeted mutation in the gene for transcription factor GATA-1. Nature 349, 257–260.

Philippe, L., Vasseur, J.J., Debart, F., and Thoreen, C.C. (2018). La-related protein 1 (LARP1) repression of TOP mRNA translation is mediated through its cap-binding domain and controlled by an adjacent regulatory region. Nucleic Acids Res 46, 1457–1469.

Rasmussen, K.D., Simmini, S., Abreu-Goodger, C., Bartonicek, N., Di Giacomo, M., Bilbao-Cortes, D., Horos, R., Von Lindern, M., Enright, A.J., and O’Carroll, D. (2010). The miR-144/451 locus is required for erythroid homeostasis. J Exp Med 207, 1351–1358.

Reichmann, D., Voth, W., and Jakob, U. (2018). Maintaining a Healthy Proteome during Oxidative Stress. Mol Cell 69, 203–213.

Rojas, D.A., Moreira-Ramos, S., Urbina, F., and Maldonado, E. (2018). Transcriptional regulation of ribosomal protein genes in yeast and metazoan cells. J Mol Cell Biochem 2, 1–6.

Rowley, P.T., Ohlsson-Wilhelm, B.M., and Farley, B.A. (1985). K562 human erythroleukemia cells demonstrate commitment. Blood 65, 862–868.

Rudra, D., Zhao, Y., and Warner, J.R. (2005). Central role of Ifh1p-Fhl1p interaction in the synthesis of yeast ribosomal proteins. EMBO J 24, 533–542.

Sanz, L.A., and Chédin, F. (2019). High-resolution, strand-specific R-loop mapping via S9.6-based DNA-RNA immunoprecipitation and high-throughput sequencing. Nat Protoc 14, 1734–1755.

Sanz, L.A., Hartono, S.R., Lim, Y.W., Steyaert, S., Rajpurkar, A., Ginno, P.A., Xu, X., and Chédin, F. (2016). Prevalent, Dynamic, and Conserved R-Loop Structures Associate with Specific Epigenomic Signatures in Mammals. Mol Cell 63, 167–178.

Saporita, A.J., Chang, H.C., Winkeler, C.L., Apicelli, A.J., Kladney, R.D., Wang, J., Townsend, R.R., Michel, L.S., and Weber, J.D. (2011). RNA helicase DDX5 is a p53-independent target of ARF that participates in ribosome biogenesis. Cancer Res 71, 6708–6717.

Schawalder, S.B., Kabani, M., Howald, I., Choudhury, U., Werner, M., and Shore, D. (2004). Growth-regulated recruitment of the essential yeast ribosomal protein gene activator Ifh1. Nature 432, 1058–1061.

Shapiro, J.S., Langlois, R.A., Pham, A.M., and Tenoever, B.R. (2012). Evidence for a cytoplasmic microprocessor of pri-miRNAs. RNA (New York, NY 18, 1338–1346.

Shapiro, J.S., Schmid, S., Aguado, L.C., Sabin, L.R., Yasunaga, A., Shim, J.V., Sachs, D., Cherry, S., and tenOever, B.R. (2014). Drosha as an interferon-independent antiviral factor. Proc Natl Acad Sci U S A 111, 7108–7113.

Shibui-Nihei, A., Ohmori, Y., Yoshida, K., Imai, J., Oosuga, I., Iidaka, M., Suzuki, Y., Mizushima-Sugano, J., Yoshitomo-Nakagawa, K., and Sugano, S. (2003). The 5’ terminal oligopyrimidine tract of human elongation factor 1A-1 gene functions as a transcriptional initiator and produces a variable number of Us at the transcriptional level. Gene 311, 137–145.

Simsek, D., and Barna, M. (2017). An emerging role for the ribosome as a nexus for post-translational modifications. Curr Opin Cell Biol 45, 92–101.

Siomi, H., and Siomi, M.C. (2010). Posttranscriptional regulation of microRNA biogenesis in animals. Mol Cell 38, 323–332.

Spagnol, G., Kieken, F., Kopanic, J.L., Li, H., Zach, S., Stauch, K.L., Grosely, R., and Sorgen, P.L. (2016). Structural Studies of the Nedd4 WW Domains and Their Selectivity for the Connexin43 (Cx43) Carboxyl Terminus. J Biol Chem 291, 7637–7650.

Staub, O., Dho, S., Henry, P., Correa, J., Ishikawa, T., McGlade, J., and Rotin, D. (1996). WW domains of Nedd4 bind to the proline-rich PY motifs in the epithelial Na+ channel deleted in Liddle’s syndrome. EMBO J 15, 2371–2380.

Sulima, S.O., Hofman, I.J.F., De Keersmaecker, K., and Dinman, J.D. (2017). How Ribosomes Translate Cancer. Cancer Discov 7, 1069–1087.

Suzuki, H.I., Young, R.A., and Sharp, P.A. (2017). Super-Enhancer-Mediated RNA Processing Revealed by Integrative MicroRNA Network Analysis. Cell 168, 1000–1014 e1015.

Suzuki, M., Shimizu, R., and Yamamoto, M. (2011). Transcriptional regulation by GATA1 and GATA2 during erythropoiesis. Int J Hematol 93, 150–155.

Tang, X., Li, M., Tucker, L., and Ramratnam, B. (2011). Glycogen synthase kinase 3 beta (GSK3beta) phosphorylates the RNAase III enzyme Drosha at S300 and S302. PLoS One 6, e20391.

Tcherkezian, J., Cargnello, M., Romeo, Y., Huttlin, E.L., Lavoie, G., Gygi, S.P., and Roux, P.P. (2014). Proteomic analysis of cap-dependent translation identifies LARP1 as a key regulator of 5’TOP mRNA translation. Genes Dev 28, 357–371.

To, T.L., Cuadros, A.M., Shah, H., Hung, W.H.W., Li, Y., Kim, S.H., Rubin, D.H.F., Boe, R.H., Rath, S., Eaton, J.K., et al. (2019). A Compendium of Genetic Modifiers of Mitochondrial Dysfunction Reveals Intra-organelle Buffering. Cell 179, 1222–1238 e1217.

Tomoda, T., Kurashige, T., Yamamoto, H., Fujimoto, S., and Taniguchi, T. (1991). Fluctuation of gene expression for poly(ADP-ribose) synthetase during hemin-induced erythroid differentiation of human leukemia K562 cells and its reversion process. Biochim Biophys Acta 1088, 359–364.

Truitt, M.L., and Ruggero, D. (2016). New frontiers in translational control of the cancer genome. Nat Rev Cancer 16, 288–304.

Ulirsch, J.C., Verboon, J.M., Kazerounian, S., Guo, M.H., Yuan, D., Ludwig, L.S., Handsaker, R.E., Abdulhay, N.J., Fiorini, C., Genovese, G., et al. (2018). The Genetic Landscape of Diamond-Blackfan Anemia. Am J Hum Genet 103, 930–947.

Wade, J.T., Hall, D.B., and Struhl, K. (2004). The transcription factor Ifh1 is a key regulator of yeast ribosomal protein genes. Nature 432, 1054–1058.

Warner, J.R. (1999). The economics of ribosome biosynthesis in yeast. Trends Biochem Sci 24, 437–440.

Warner, J.R., and McIntosh, K.B. (2009). How common are extraribosomal functions of ribosomal proteins? Mol Cell 34, 3–11.

Welch, J.J., Watts, J.A., Vakoc, C.R., Yao, Y., Wang, H., Hardison, R.C., Blobel, G.A., Chodosh, L.A., and Weiss, M.J. (2004). Global regulation of erythroid gene expression by transcription factor GATA-1. Blood 104, 3136–3147.

Wu, H., Xu, H., Miraglia, L.J., and Crooke, S.T. (2000). Human RNase III is a 160-kDa protein involved in preribosomal RNA processing. J Biol Chem 275, 36957–36965.

Yang, Q., Li, W., She, H., Dou, J., Duong, D.M., Du, Y., Yang, S.H., Seyfried, N.T., Fu, H., Gao, G., et al. (2015). Stress induces p38 MAPK-mediated phosphorylation and inhibition of Drosha-dependent cell survival. Mol Cell 57, 721–734.

Zengel, J.M., and Lindahl, L. (1994). Diverse mechanisms for regulating ribosomal protein synthesis in Escherichia coli. Prog Nucleic Acid Res Mol Biol 47, 331–370.

Zon, L.I., Youssoufian, H., Mather, C., Lodish, H.F., and Orkin, S.H. (1991). Activation of the erythropoietin receptor promoter by transcription factor GATA-1. Proc Natl Acad Sci U S A 88, 10638–10641.

