## Supplemental Figurse and Tables for "Control of ribosomal protein synthesis by Microprocessor complex"

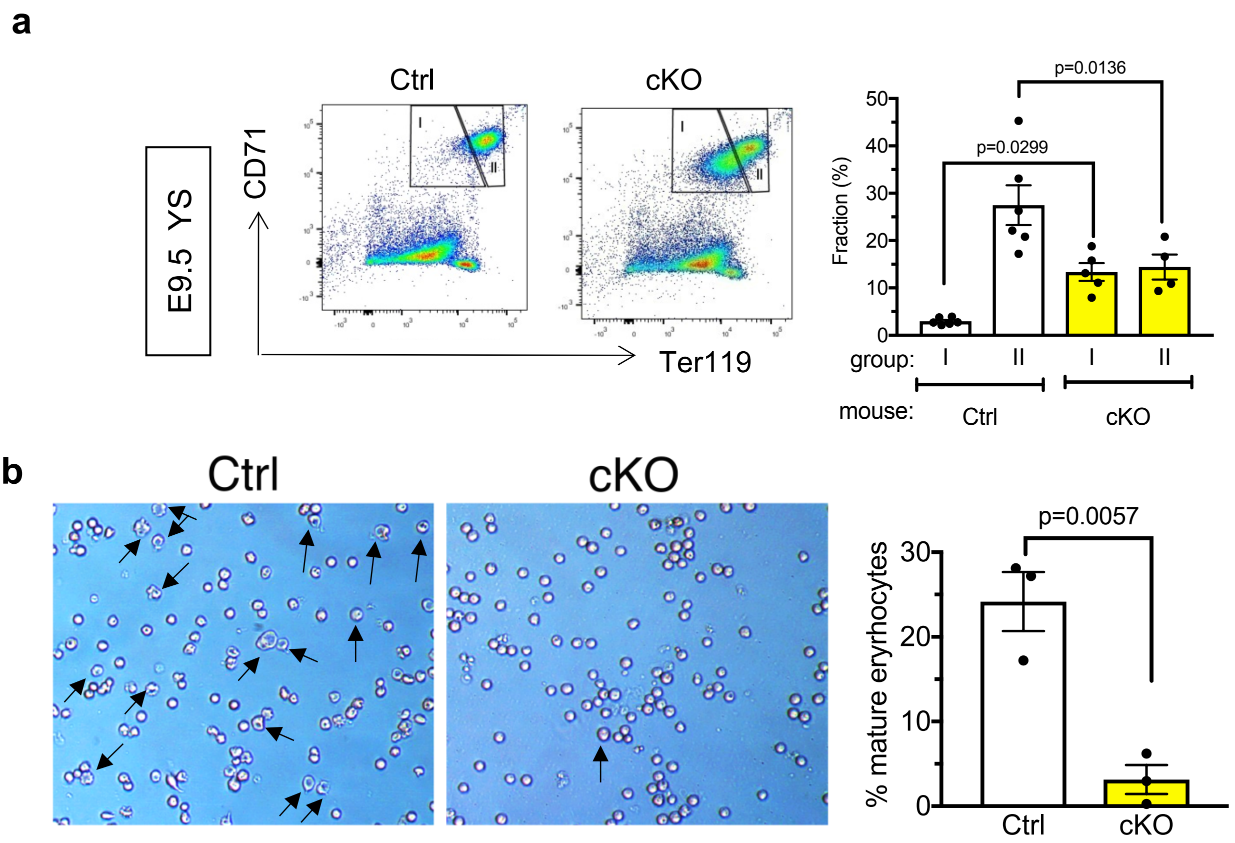


**Supplementary Fig. S1 Endothelial-specific deletion of *Drosha* impairs erythropoiesis. a.** Representative images of flow cytometry analyses on pro-erythroblasts (group I: CD71$^{high}$Ter119$^{low}$) and erythroblasts (group II: CD71$^{high}$Ter119$^{high}$) derived from yolk sac of E9.5 Ctrl or cKO embryos (left panel). Quantitation is shown on the right panel as frequency (%) of total live (DAPI$^{-}$) cells (Mean $\pm$ SEM). n=Ctrl: 6 embryos, cKO: 4 embryos. 2 litters. **b.** Morphology of CD71$^{+}$Ter119$^{+}$Itga4$^{low}$ cells sorted from the peripheral blood of E10.5 Ctrl or CD71$^{+}$Ter119$^{+}$Itga4$^{high}$ cells from Drosha-cKO embryos, followed by being cultured in differentiation media for 3 days (left). Cells with a mature erythrocyte-like morphology are indicated by arrows (left) and the quantitation is shown (right).

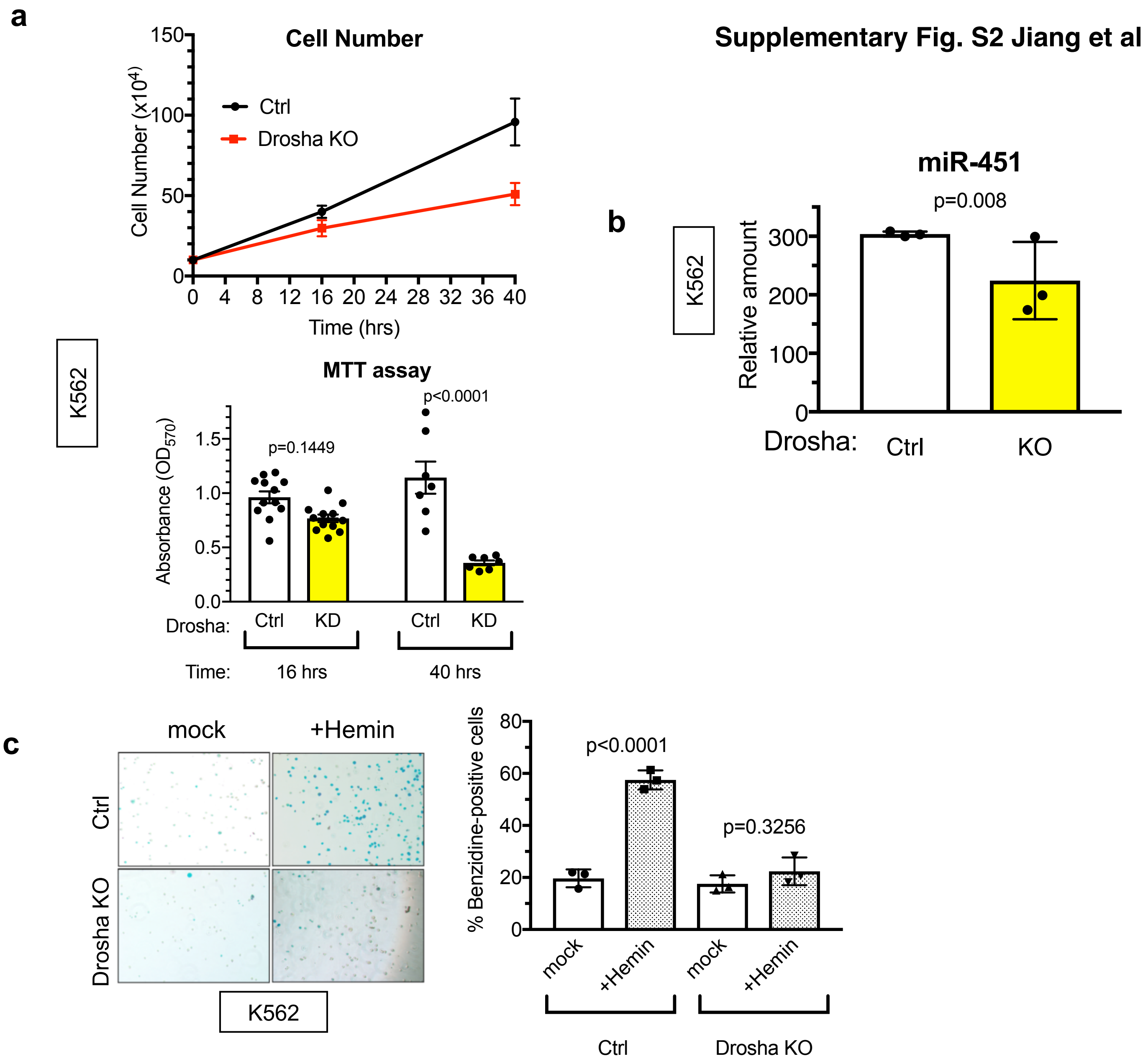


**Supplementary Fig. S2 Knockdown of Drosha in K562 cells affects cell proliferation rate and impairs erythroid maturation. a.** Cell counting (top) and MTT assay (bottom) of K526 cells infected with lentivirus expressing CRISPR-Cas9 with non-specific gRNA (Ctrl) or gRNA against *Drosha* (KO). Data were collected at 16 and 40 hrs after inoculation ($1x10^5$ cells) and plotted as Mean $\pm$ SEM. n=5. **b.** qRT-PCR analysis of *miR-451* (relative to U6 snRNA) in Ctrl or *Drosha* KO K562 cells. n=3. **c.** Erythroid differentiation was induced by hemin treatment in Ctrl or *Drosha* KO K562 cells. Representative images of benzidine staining (left). A fraction (%) of benzidine-positive (blue) cells were plotted as Mean $\pm$ SEM (right). n=3.

Supplementary Fig. S3 Jiang et al.

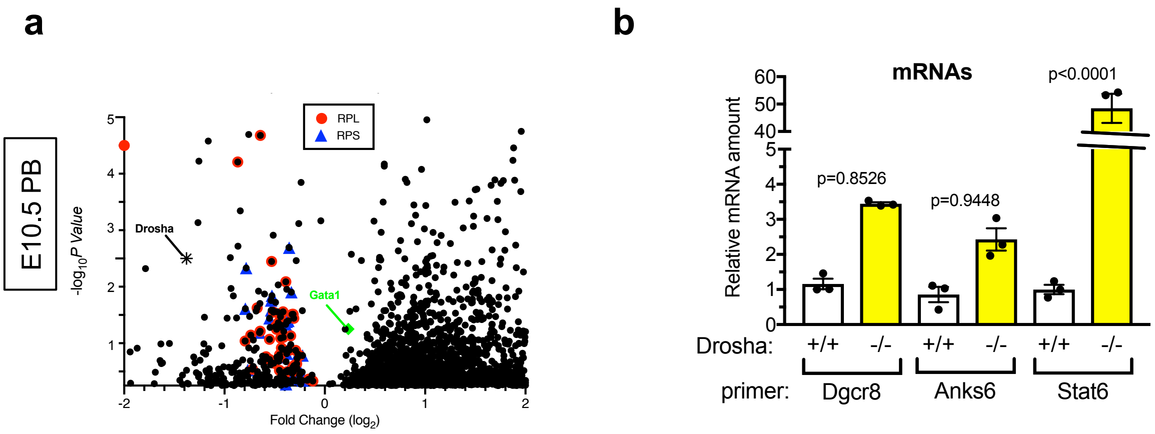


**Supplementary Fig. S3 mRNAs processed by Drosha are increased in the Drosha(-/-) cells. a.** Valcano plot of RNA seq. The log2 fold change of Drosha (-/-) cells normalized with Drosha (+/+) EPCs is plotted. **b.**qRT-PCR analysis of Dgcr8, Anks6 and Stat6 mRNAs relative to GAPDH in Drosha(+/+) erythroid progenitors from Ctrl mice and Drosha (-/-) erythroid progenitors from cKO mice sorted from E10.5 peripheral blood. (Mean ± SEM). n=Ctrl: 5 embryos, cKO: 5 embryos. 3 litters.

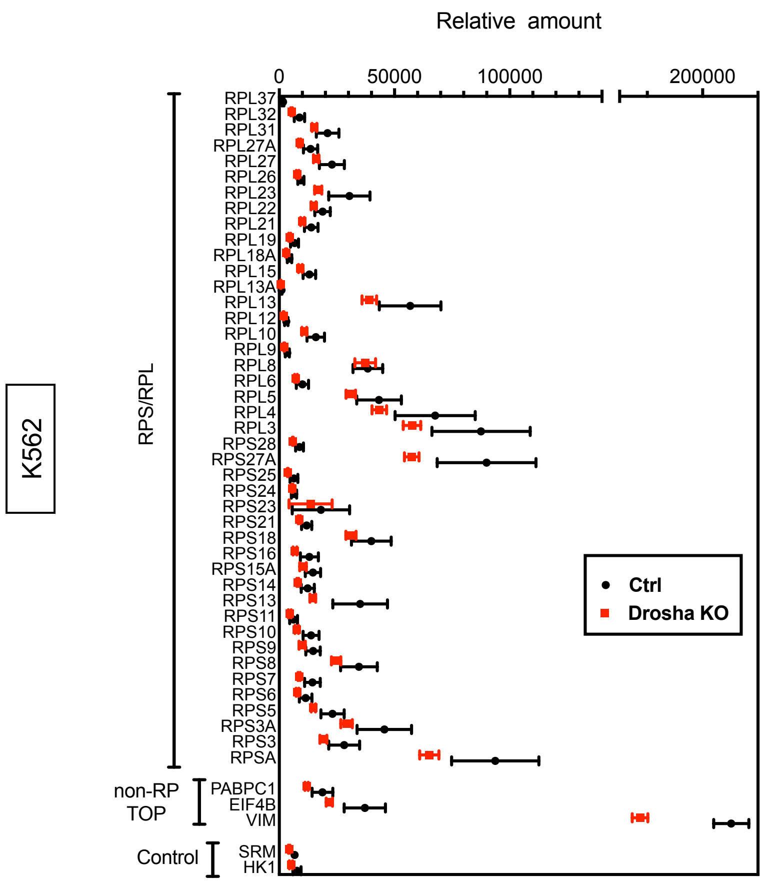


**Supplementary Fig. S4 Quantitative proteomic analysis shows reduced production of RPs and non-RP TOP genes in *Drosha* KO K562 cell compared to Ctrl K562 cells**
Tandem mass-tag (TMT)-based mass-spectrometry was performed and the result was plotted as Mean±SEM; n=5

### Rps genes



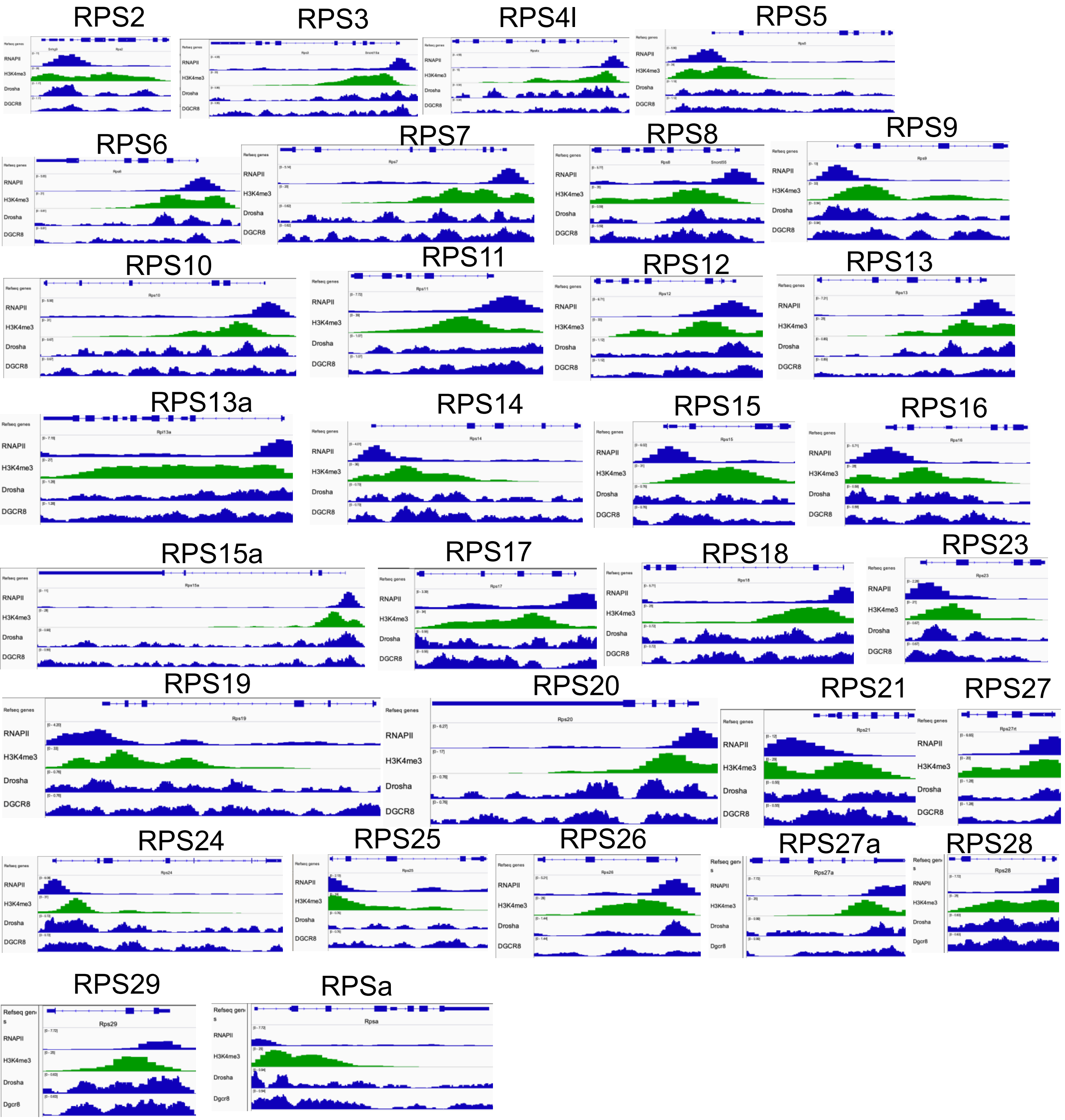
RPS2
RPS3
RPS4l
RPS5
RPS6
RPS7
RPS8
RPS9
RPS10
RPS11
RPS12
RPS13
RPS13a
RPS14
RPS15
RPS16
RPS15a
RPS17
RPS18
RPS23
RPS19
RPS20
RPS21
RPS27
RPS24
RPS25
RPS26
RPS27a
RPS28
RPS29
RPSa
Refseq genes
RNAPII
H3K4me3
Drosha
DGCR8
Dgcr8

Supplementary Fig. S5-2 Jiang et al.

### Rpl genes

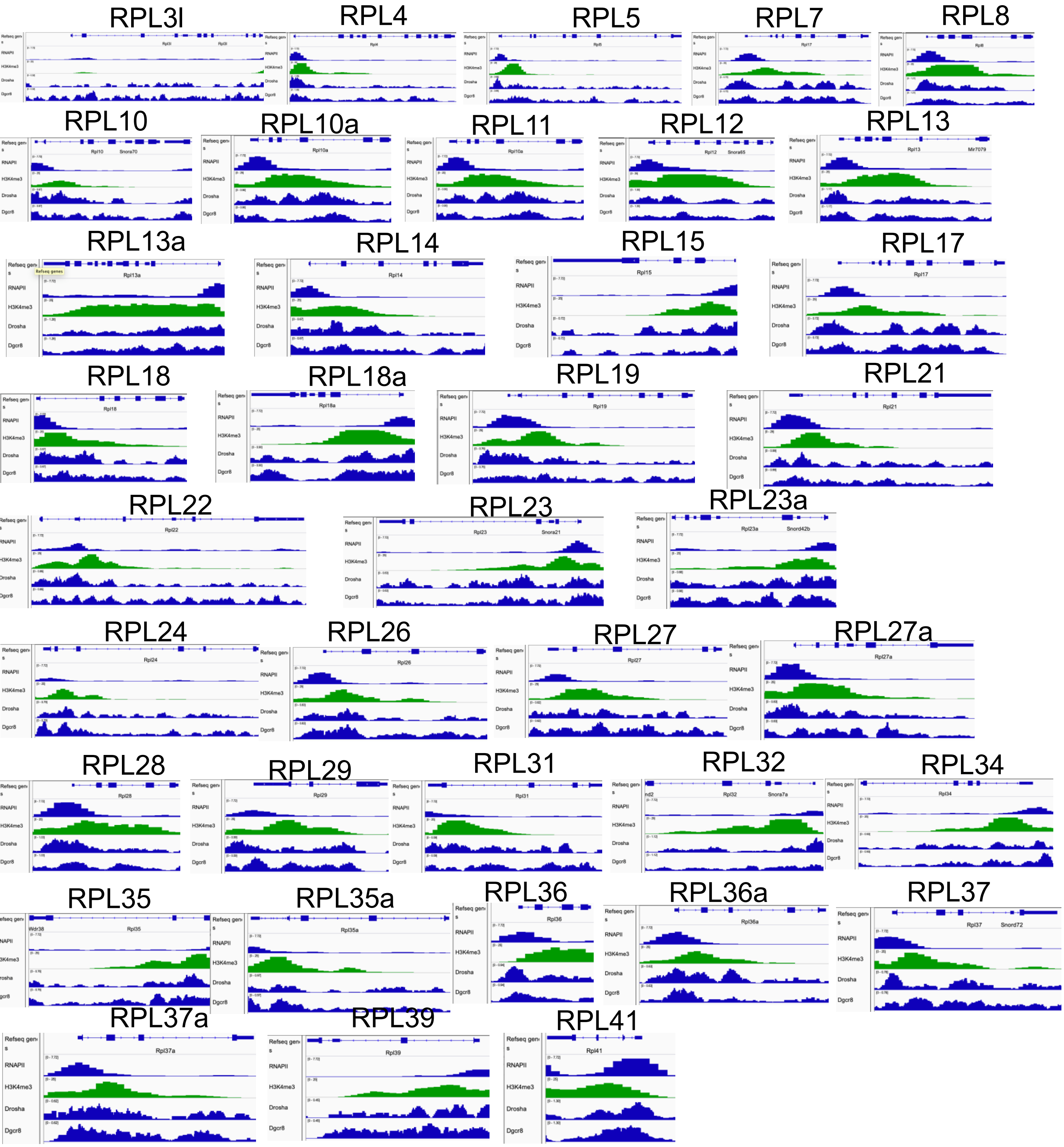


### Non-RP genes

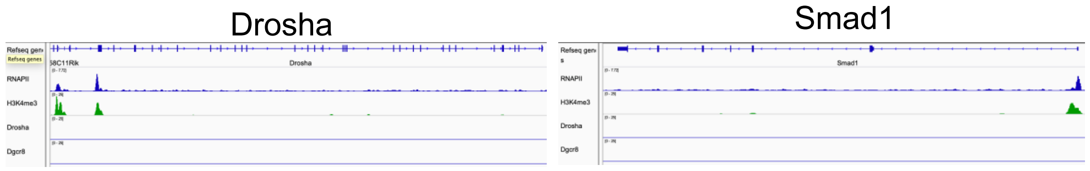

**Supplementary Fig. S5 Association of Drosha/DGCR8 at the RP gene loci.** ChIP-seq analysis of RNA polymerase Ii (RNAPII) and Histone Lysine-4 trimethylation (H3K4me3), Drosha, and Dgcr8, at Rps, Rpl, control (Drosha, Smad1) gene loci in mouse ES cells.

**Supplementary Fig. S6-1 Jiang et al.**

### Rps genes

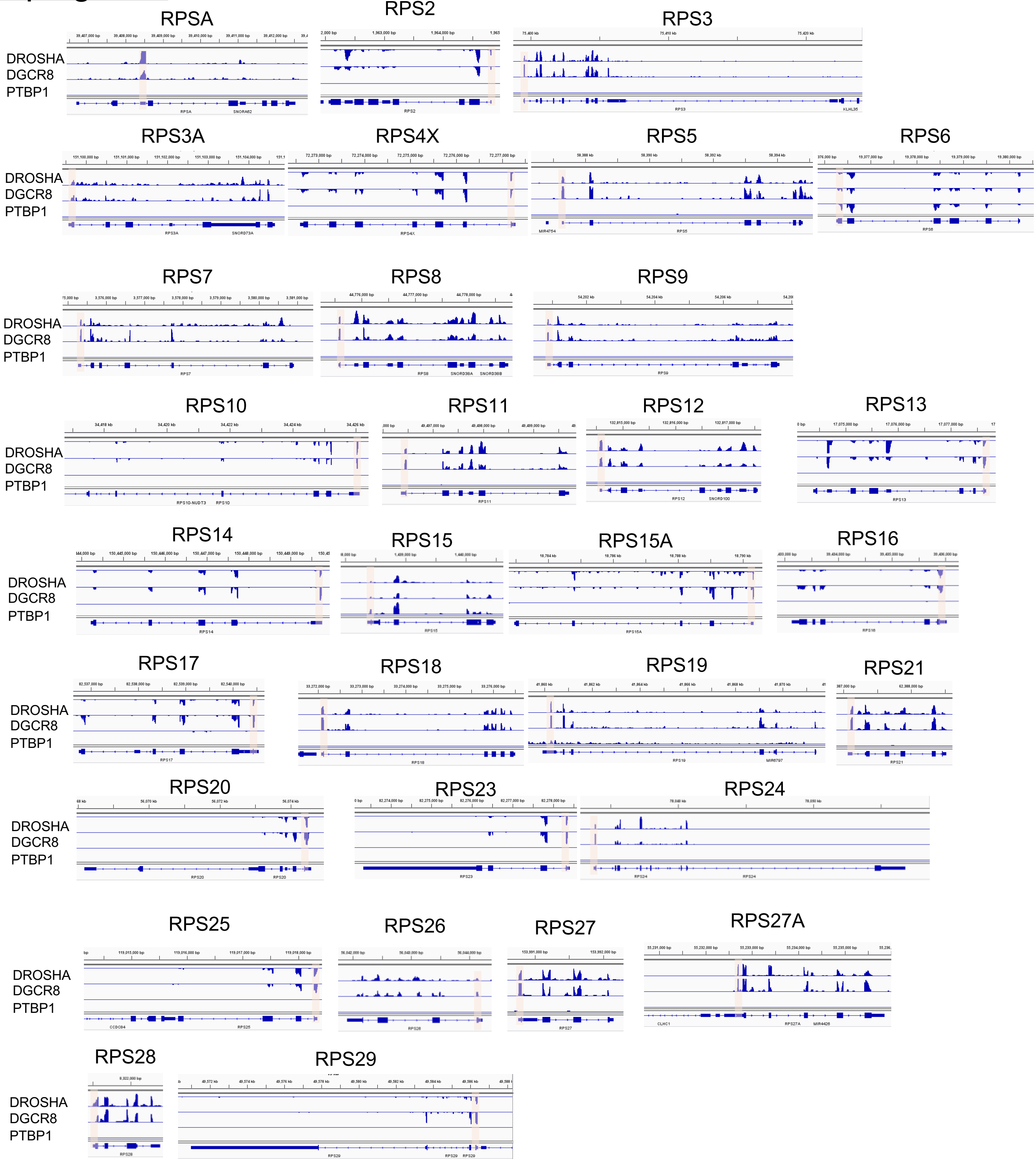

### Rpl genes

Supplementary Fig. S6-2 Jiang et al.

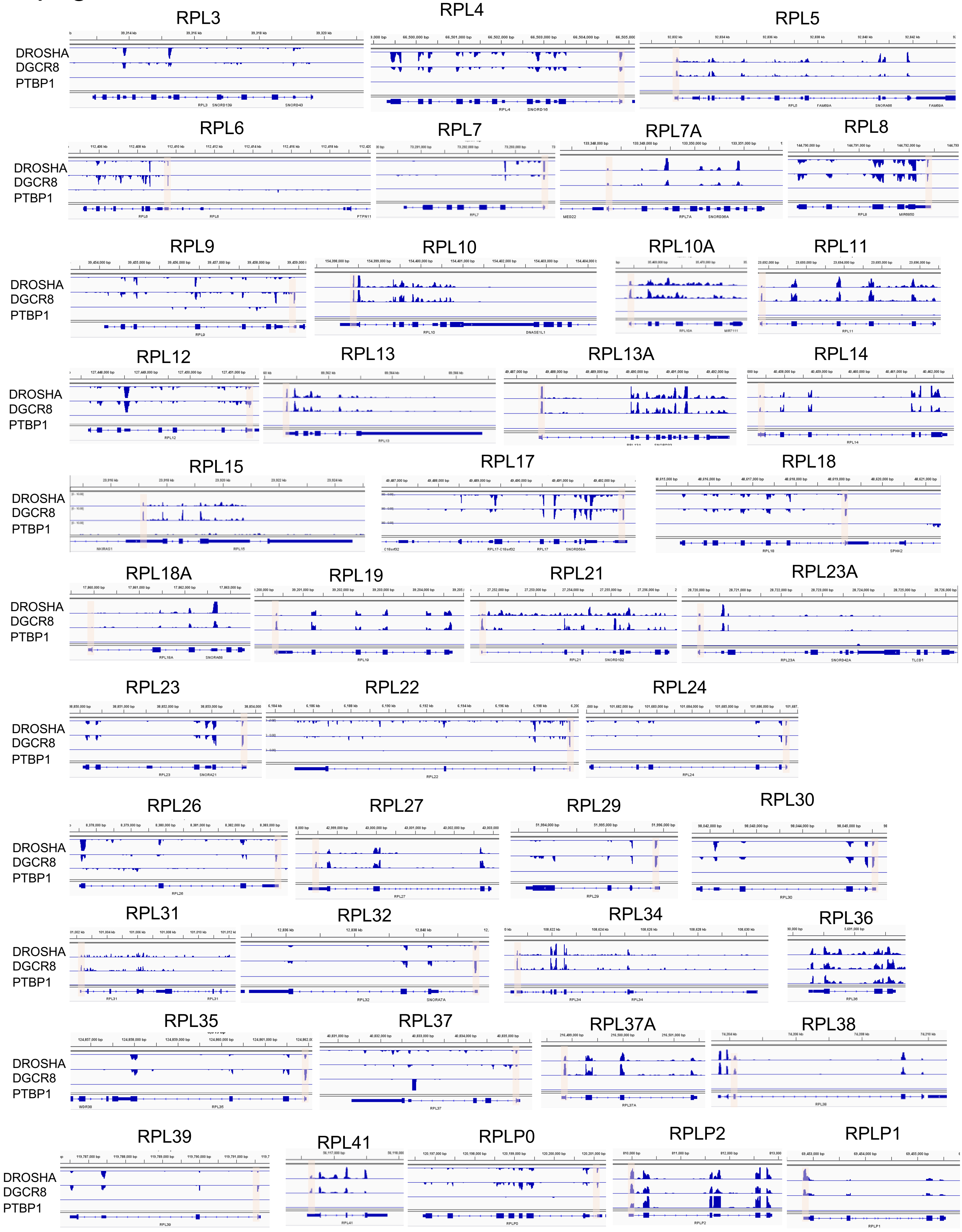

### Control genes

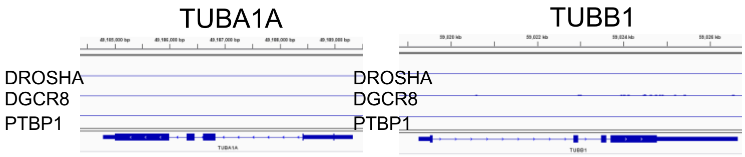


### Non-RP TOP genes

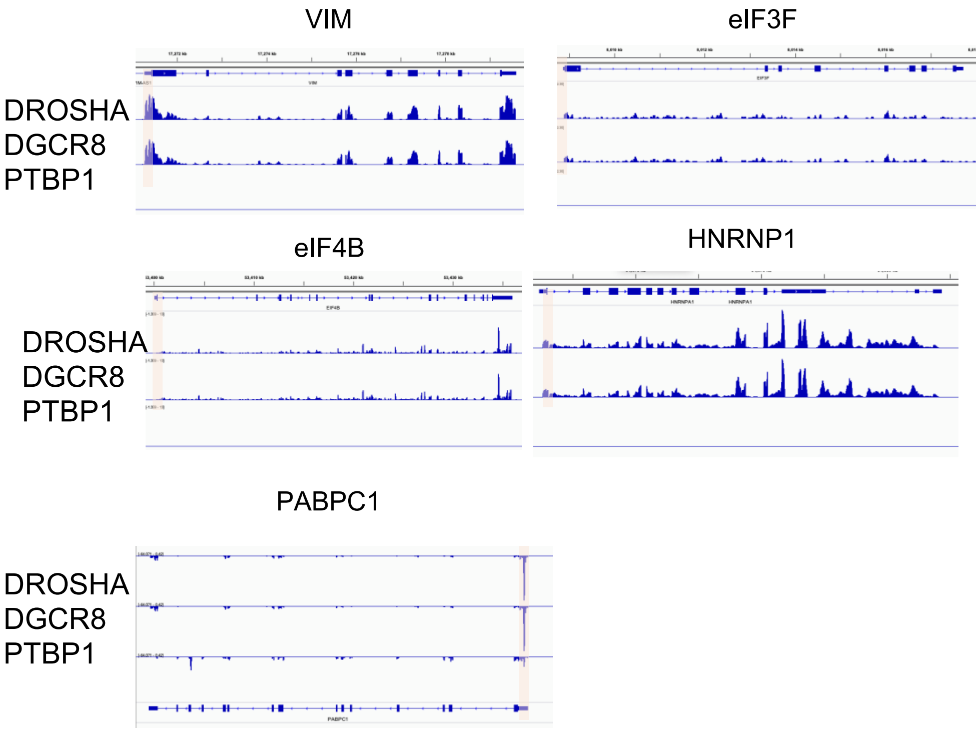


**Supplementary Fig. S6 Drosha/DGCR8 preferentially binds with 5' end of RP mRNAs.** e-CLIP profiles of Drosha, Dgcr8 and PTBP1 at Rps, Rpl, Tub1A1/TubB1 (control), and non-RP TOP genes (VIM, eIF3F, eIF4B, HNRNP1, and PABPC1) loci in K562 cells. Orange shade indicates the location of the 5'-TOP motif where Drosha and Dgcr8 bind peak are detected by eCLIP.

#### Rps genes

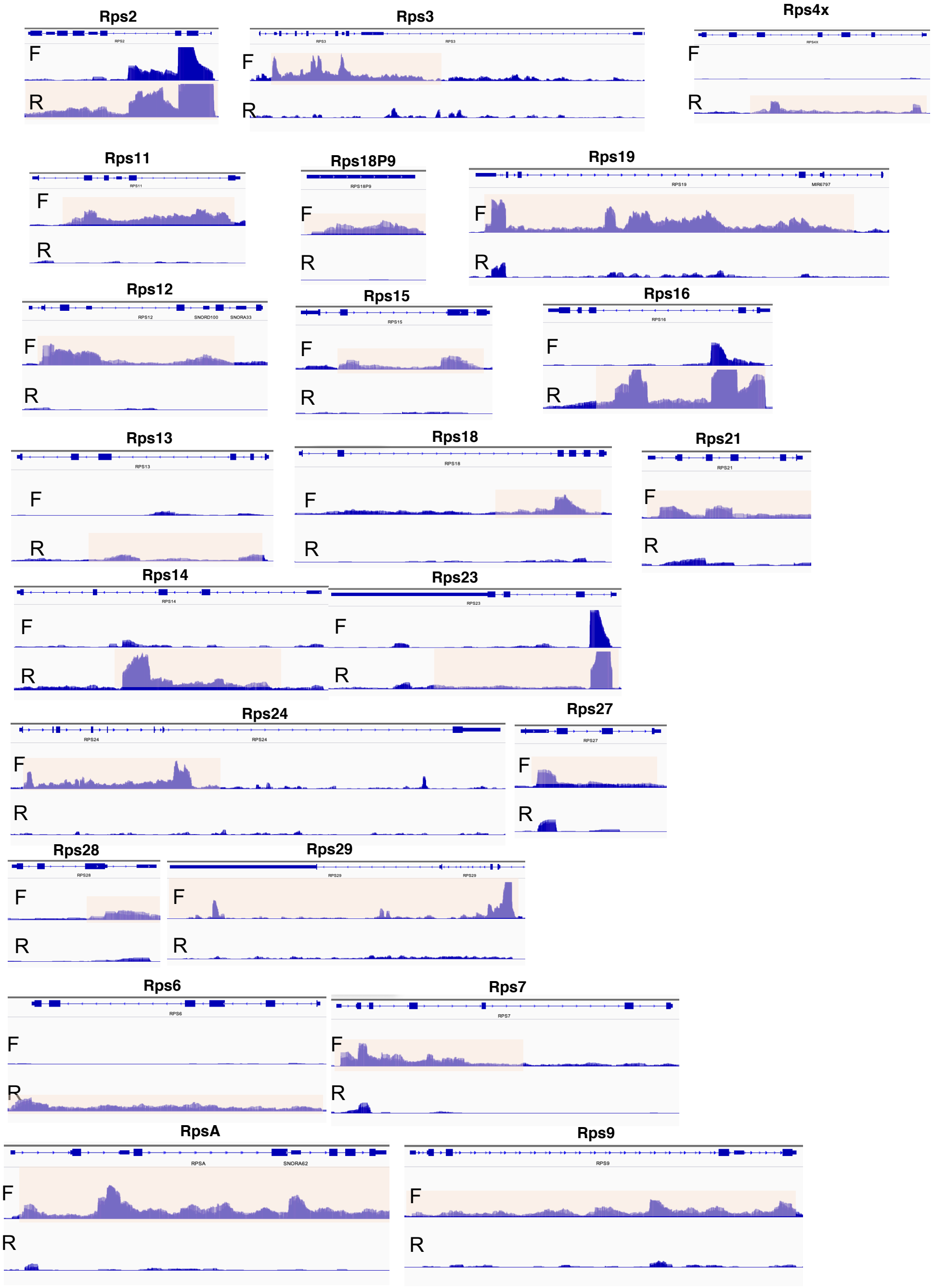
Rps2
F
R
Rps3
F
R
Rps4x
F
R
Rps11
F
R
Rps18P9
F
R
Rps19
F
R
Rps12
F
R
Rps15
F
R
Rps16
F
R
Rps13
F
R
Rps18
F
R
Rps21
F
R
Rps14
F
R
Rps23
F
R
Rps24
F
R
Rps27
F
R
Rps28
F
R
Rps29
F
R
Rps6
F
R
Rps7
F
R
RpsA
F
R
Rps9
F
R

Supplementary Fig. S7-2 Jiang et al.

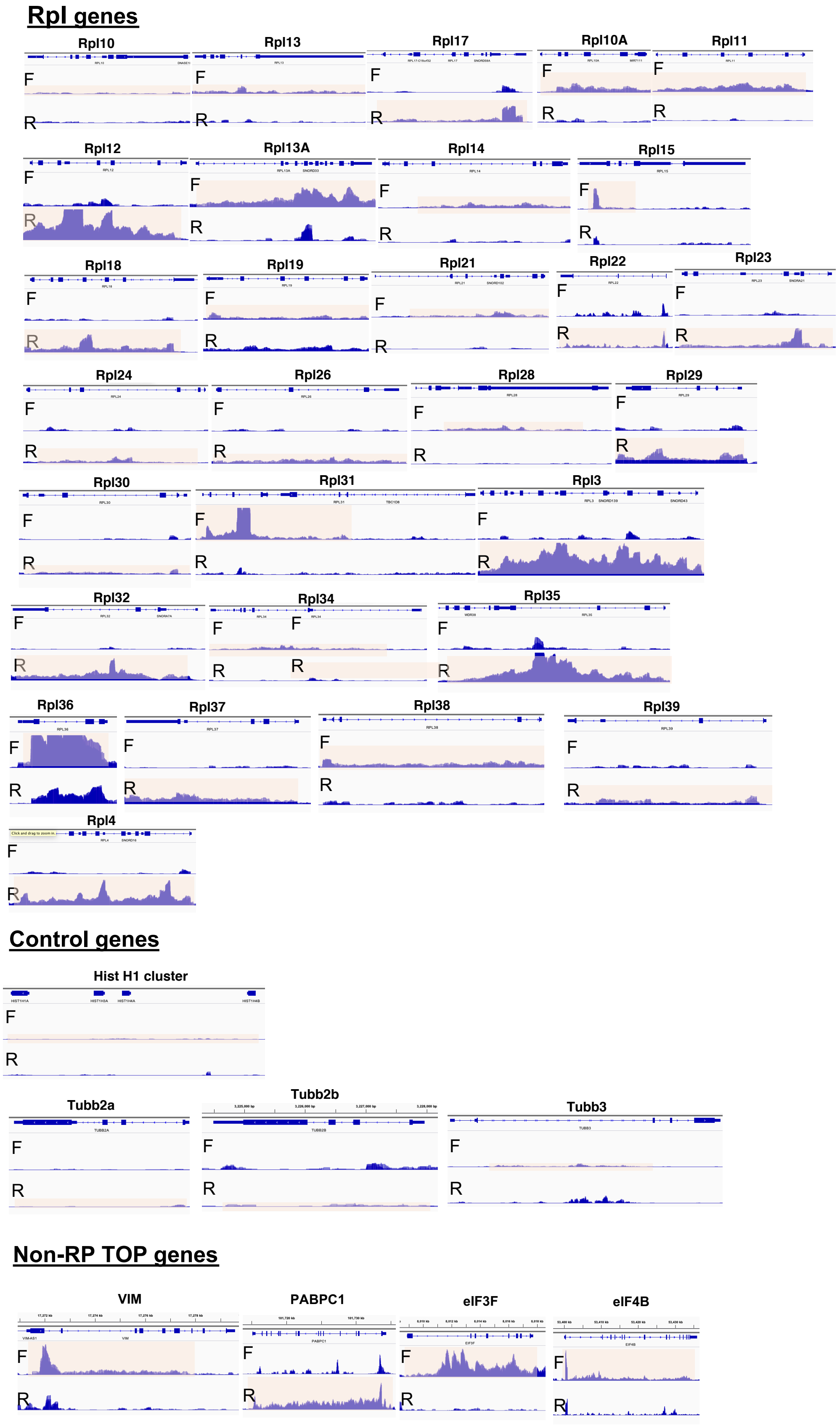


**Supplementary Fig. S7 Formation of R-loops at the RP gene loci.** DRIP seq analysis of the Rps, Rpl, Hist H1 cluster, Tub2a, Tub2b, and Tubb3 (control), and non-RP TOP genes (VIM, eIF3F, eIF4B, HNRNP1, and PABPC1) loci in HeLa cells. Orange shade indicate DRIP peaks at the proper strand.

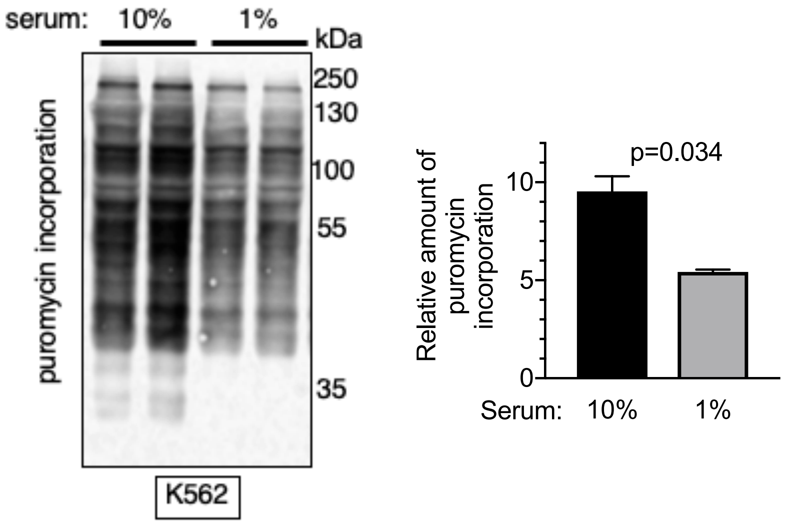


**Supplementary Fig. S8 Reduction of global protein synthesis upon serum starvation.** K562 cells were cultured in normal (10% serum) or serum-starved (1% serum) for 16 hrs, followed by puromycin treatment for 10 min. Total lysate were generated from an equal number of cells ($1x10^6$), followed by immunoblot analysis with anti-puromycin antibody (**left**). The average of the abundance of puromycin-incorporated proteins in duplicate samples is shown (**right**).

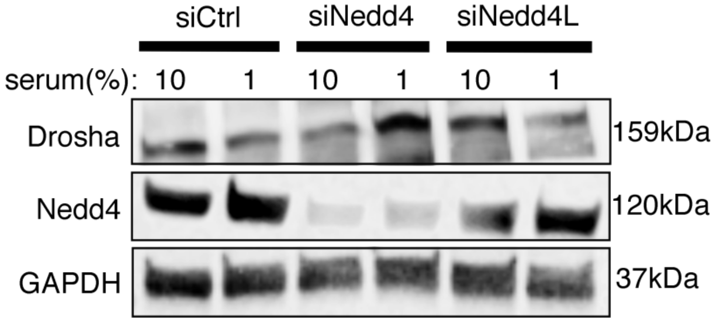


**Supplementary Fig. S9 Nedd4L does not play a role in Drosha degradation** Protein amount of Drosha, Nedd4, and GAPDH (loading control) was examined by western blot in total cell lysates from K562 cells transfected with non-specific control siRNA (siCtrl), siRNA against Nedd4 (siNedd4) or Nedd4L (siNedd4L) cultured in growth media (10% serum) or serum starvation media (1% serum) for 6 hrs.

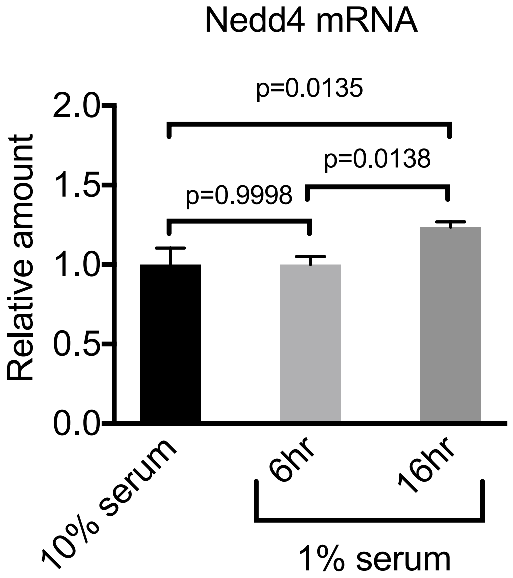


**Supplementary Fig. S10 The amount of Nedd4 mRNA is not elevated upon serum starvation** K562 cells were cultured in normal (10%) or low serum (1%) condition for 6 or 16 hr, followed by RT-PCR analysis of Nedd4 mRNA amount relative to GAPDH mRNA in triplicates. Mean±SEM is shown.

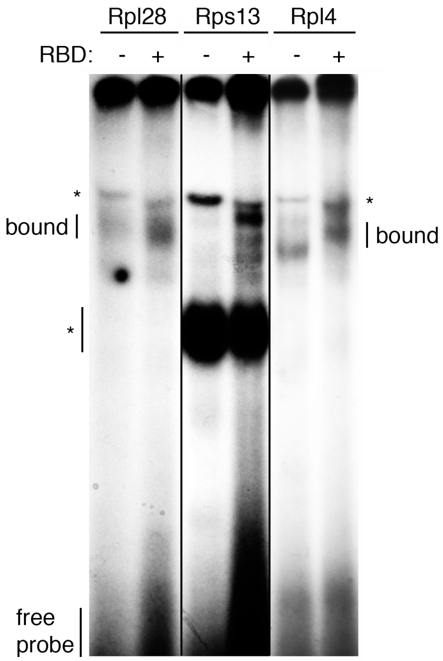


Rpl28
5’- CUCUUUCCGUCUCAGGUCGCCGCUGCGAAGGGAGGUG-3’

Rps13
5’-CCUUUCGUUGCCUGAUCGCCGCCAUCAUGGGUCGCAUGCA-3’

Rpl4
5’-CUUUUCCUGUGGCAGCAGCCGGGCUGAGAGGAGCGUGGCU-3’

**Supplementary Fig. S11 Drosha RNA binding domain (RBD) binds Rps13 and Rpl4**
Representative image of RNA EMSA with Drosha RBD domain, radiolabeled Rpl28, Rps13, and Rpl5 probe (top). Bottom: Sequences of the RNA probes (left) and a predicted secondary structure (right) are shown. TOP motif is highlighted in yellow.

#### Supplementary Information Tables

**Supplementary Table 1:Data Resource of high throughput RNA seq, ChIP seq, eCLIP seq and DRIP seq.** Type of experiment, cell type, accession number and reference are listed.

| Experiment | Cell | Accession Number | Reference |
|---|---|---|---|
| RNA seq | Crispr gRNA against no target in HepG2 | ENCODE: ENCSR481WJH | |
| RNA seq | Crispr gRNA against Drosha in HepG2 | ENCODE: ENCSR991XFO | |
| RNA seq | Control shRNA in K562 cells | ENCODE:ENCSR898NWE | |
| RNA seq | shRNA against AGO2 in K562 cells | ENCODE: ENCSR495YSS | |
| RNA seq | WT and Drosha KO HCT116 cells | GSE80258 | Jeong, et al. BMC Genomics 2016 |
| RNA seq | Control shRNA in HepG2 cells | ENCODE: ENCSR776SXA | |
| RNA seq | shRNA against DDX5 in HepG2 cells | ENCODE: ENCSR808FBR | |
| ChIP seq | DGCR8 and Drosha in mESC | GSE89826 | Suzuki et al. Cell 2017 |
| ChIP seq | RNAPII in ES-Bruce4 | GSM918749 | Pope et al. Nature 2014 |
| ChIP seq | H3K4me3 in ES-Bruce4 | GSM769008 | Pope et al. Nature 2014 |
| Single Cell RNA seq | E9.5 embryonic cardiomyocyte | GSE96648 | Chen et al. Protein& Cell 2018 |
| eCLIP | K562 PTBP1 mock input | ENCODE:ENCSR445FZX | |
| eCLIP | K562 PTBP1 | ENCODE:ENCSR981WKN | |
| eCLIP | K562 DGCR8 mock input | ENCODE:ENCSR616BTD | |
| eCLIP | K562 DGCR8 | ENCODE: ENCSR616BTD | |
| eCLIP | K562 Drosha mock input | ENCODE:ENCSR947JVR | |
| eCLIP | K562 Drosha | ENCODE:ENCSR828NSY | |
| RNA sequencing | Crispr gRNA against no target in HepG2 | ENCODE:ENCSR481WJH | |
| DRIP seq | Hela | GSM1518913 | |

| | | | |
|---|---|---|---|
| DRIP seq | Si-Control/si-Drosha in U2OS | GSE97648 | Lu et al. Nat.Comm. 2018 |

**Supplementary Table 2: Primers for mouse genotyping, qPCR, ChIP and DRIP.** First letter in the name for primers indicate species. M stands for mouse; H stands for human.

| | | Primer Sequence (5' to 3') |
|---|---|---|
| **Primers for Genotyping** | mDrosha-F | GCA GAA AGT CTC CCA CTC CTA ACC TTC |
| | mDrosha-R | CCA GGG GAA ATT AAA CGA GAC TCC |
| | mCre-F | TGC CAC GAC CAA GTG ACA GCA |
| | mCre-R | AGA GAC GGA AAT CCA TCG CTC |
| **Primers for qRT-PCR** | mAnks6-F | GGATCTTGCTGAATACCTGGACC |
| | mAnks6-R | TTCGTCCGCAATCTCCTTCACC |
| | mStat6-F | ACGACAACAGCCTCAGTGTGGA |
| | mStat6-R | CAGGACACCATCAAACCACTGC |
| | mDGCR8-F | CAAGTGAGCCTTTTGGTGCC |
| | mDGCR8-R | GATGTGGTTAAAATACTCCAGTTCT |
| | mDrosha-F | GATAGGAGCTGTTTACTTGGAGG |
| | mDrosha-R | AGTTGCCGATCCGTATTTGG |
| | mGata1-F | ACGACCACTACAACACTCTGGC |
| | mGata1-R | TTGCGGTTCCTCGTCTGGATTC |
| | mFog1-F | TGTGAACGCCATCTCAAGGTGC |
| | mFog1-R | TGTCACCAGGTGACTGTAGAGG |
| | mTal1-F | GCCAGCCGCTCGCCTCACTA |
| | mTal1-R | CCGCACTACTTTGGTGTGAGGA |
| | mKlf1-F | CGGCGAACTTTGGCACCTAAGA |
| | mKlf1-R | AGGAGCAGGCATAAGGCTTCTC |
| | mAlas2-F | GTCTTCAGACACAATGACCCAGG |
| | mAlas2-R | TTCCTCCAGAGGACAGATGGCA |
| | mEbp4.9-F | CATCCTGGACATTGAGCGACCT |

| | | |
|---|---|---|
| | mEbp4.9-R | GGGATGTGGATTTGGGTGACAG |
| | mRps7-F | AGTTCAGTGGCAAGCACGTGGT |
| | mRps7-R | CTAAGTCCTCAAGGATGGCGTC |
| | mRps10-F | GGTCGCCAAAAAGGATGTCCAC |
| | mRps10-R | GCAAACTGTTCCTTCACGTAGCC |
| | mRps17-F | TGAGAGGCATCTCTATCAAGTTGC |
| | mRps17-R | GCTTCAGCATCTCCTTGGTGTC |
| | mRps19-F | GAGAACTGGTTCTACACACGAGC |
| | mRps19-R | CGTTTCTCTGCCGTCCTCCATA |
| | mRps20-F | GCAACGTGAAGTCGCTGGAGAA |
| | mRps20-R | CCTTCACCACAAGGTGTTTTTCTG |
| | mRps24-F | TGGTGGCAAGACCACTGGCTTT |
| | mRps24-R | CTGTTCTTGCGTTCCTTTCGCTG |
| | mRps26-F | CGGAACATTGTAGAAGCCGCTG |
| | mRps26-R | CCTTGCTATGGATGGCACAGCT |
| | mRps27-F | CCTACTTTATGGACGTGAAATGCC |
| | mRps27-R | CTGTCAGCCTTGCTTTTCCACC |
| | mRpl11-F | GAGAGCGGAGACAGACTGACC |
| | mRpl11-R | GGATGCCAAAGGACCTGACAGT |
| | mRpl30-F | TCCTTGCCAACAACTGTCCAGC |
| | mRpl30-R | TTTCCACACGCTGTGCCCAGTT |
| | mRpl34-F | GCACCTAAATCTGCATGTGGCG |
| | mRpl34-R | TGTCACGGACACACTTGGCACA |
| | mRpl35a-F | TTCTACTTAGGCAAGAGATGTGCT |
| | mRpl35a-R | TTGGCACGAACCATACCGCTGT |
| | mRpl36a-F | GGTGACACAGTACAAGAAGGGC |
| | mRpl36a-R | TCAACGCACTCCAGTCTCAGCA |

| | | |
|---|---|---|
| | mRplp1-F | CAATGCCCTCATTAAAGCAGCTG |
| | mRplp1-R | CCCTACATTGCAGATGAGGCTC |
| | hGata1-F | CACGACACTGTGGCGGAGAAAT |
| | hGata1-R | TTCCAGATGCCTTGCGGTTTCG |
| | h18S-F | CTACCACATCCAAGGAAGCA |
| | h18S-R | TTTTTCGTCACTACCTCCCCG |
| | hATF4-F | TTCTCCAGCGACAAGGCTAAGG |
| | hATF4-R | CTCCAACATCCAATCTGTCCCG |
| | hRps19-F | ACTTCAGCCGAGGCTCCAAGAG |
| | hRps19-R | CTCTTTGTCCCTGAGGTGTCAG |
| **Primers for ChIP** | mRpl28-F | CTCAATATTTGCATCCACAGGC |
| | mRpl28-R | TCTGTTCAGTCTGCACTTAGC |
| | mRpsa-F | CTTAGCCTGCTTTGCCATTG |
| | mRpsa-R | GGGTAGCGCGAAAGGAC |
| | mRps2-F | GCGTAGGCAGTCGAGATC |
| | mRps2-R | CTCACGTGTTCTCCCGAAG |
| | mRps10-F | AGTGTACCAGCCATGTATGC |
| | mRps10-R | AGGGCTAAGAGTTCAGGGATAG |
| | mRps12-F | CTTTCAGTGCGTTCAAGATTCG |
| | mRps12-R | GGAGAAAAGCCCAGACCAG |
| | mRps15a-F | TTCCGCTTCCCATAATTCCG |
| | mRps15a-R | GGTAAAGGGATGGCACAGATAC |
| | mRps18-F | GGTCCCTTCCAAAACTGTCTC |
| | mRps18-R | GAGAGCGGAAGTGACGTATAAG |
| | mRps26-F | GATCCTCTTTCGAGTCTTGGC |
| | mRps26-R | ATCTACTTCTACCCCAGTACCC |
| | mRpl8-F | GACCCGTCTATTTCCTCTTTCG |

| | | |
|---|---|---|
| | mRpl8-R | CATGACAGACCGCCAAGTAG |
| | mRpl13-F | GAGATACCCACACTGCGAAG |
| | mRpl13-R | GCCAATCTCAAACGCCAATC |
| | mRpl14-F | CCCTCGGCTGTGTCTTAATT |
| | mRpl14-R | GCCTCAGTCTCCCAAATGTT |
| | mRpl19-F | CCTTTAACTCCTTTCTCCCCAG |
| | mRpl19-R | CTTTGGACACTCACCCCATC |
| | mRpl34-F | CCTAGGATAAAGGCATCGTGG |
| | mRpl34-R | CGTGAGAAGTTAGGTGTAGGG |
| | mRpl36-F | GGAAAGGCCTGCAAGTGG |
| | mRpl36-R | CTTACCTGATGCTCTCCGATG |
| | mRpl41-F | GAGCACGCCATTAAATAGCAG |
| | mRpl41-R | GTTAACGGATTGCACTGGAAAC |
| | hRPS2-ChIP-F | GCCACACTGACTAGTTCCTTC |
| | hRPS2-ChIP-R | CCACTGCCGAAACCTCC |
| | hRPS5-ChIP-F | TTCTAGCTTTTCCCCATAGCG |
| | hRPS5-ChIP-R | GCATCGTGTCCCTAAGAAATGG |
| | hRPS26-ChIP-F | GTCCGTGCCTCCAAGATG |
| | hRPS26-ChIP-R | CTCTCACTTCCCCACAATCC |
| | hRPL4-ChIP-F | CATCTTAGAGGCAACGTGGG |
| | hRPL4-ChIP-R | CTAGGCAGAGACAAGCATCC |
| | hRPL14-ChIP-F | CGTTTGCTGATTTCGTCCATC |
| | hRPL14-ChIP-R | CTCGCTACATATGTCACCTTCTG |
| | hRPL19-ChIP-F | AACGGAGCGAACAAGACC |
| | hRPL19-ChIP-R | ACGTGAAAGCAGAGATCGG |
| | hRPS2-ChIP-F(for RNAPII) | CGCCTTTTCTCCCTTTAGTGG |

| | | |
|---|---|---|
| | hRPS2-ChIP-R(for RNAPII) | CTCCAGGGACTTGATCTTCATG |
| | hRPS26-ChIP-F(for RNAPII) | TTGCGGAGAAACAGGAGATC |
| | hRPS26-ChIP-R(for RNAPII) | TGCGACAGGATGGGAAAAC |
| | hRPL14-ChIP-F(for RNAPII) | GAATTCCCAGAGTTCGAGGTG |
| | hRPL14-ChIP-R(for RNAPII) | TCAATAAACCAGACCCAGGC |
| | hRPL28-ChIP-F(for RNAPII) | TCTCAAAGCAAGTCAGGGTG |
| | hRPL28-ChIP-R(for RNAPII) | GGGAATCAAATGGCTGCAAC |
| | hChIP-NC-F1 | TTTTGTCTCCGCCCTACAC |
| | hChIP-NC-R1 | TTAGATGTCAACAGTGCCCTC |
| | mChIP-NC-F | GGCCTTCAACTCACGACTCT |
| | mChIP-NC-R | CGCCTGCACCTTGTAAACC |
| **Primers for DRIP** | hRPS2-DRIP-F | CGCCTTTTCTCCCTTTAGTGG |
| | hRPS2-DRIP-R | CTCCAGGGACTTGATCTTCATG |
| | hRPS5-DRIP-F | TGGGCCTTTTCTCATTAGGTC |
| | hRPS5-DRIP-R | GTTCAAACCAAGTTCCCCAAG |
| | hRPS10-DRIP-F | TGTGCCCAACCTTCATGTC |
| | hRPS10-DRIP-R | GAAGATCCCTCCATCCCATTC |
| | hRPS26-DRIP-F | TTGCGGAGAAACAGGAGATC |
| | hRPS26-DRIP-R | TGCGACAGGATGGGAAAAC |
| | hRPL4-DRIP-F | GAATTCCCAGAGTTCGAGGTG |

| | | |
|---|---|---|
| | hRPL4-DRIP-R | TCAATAAACCAGACCCAGGC |
| | hRPL14-DRIP-F | GAATTCCCAGAGTTCGAGGTG |
| | hRPL14-DRIP-R | TCAATAAACCAGACCCAGGC |
| | hRPL19-DRIP-F | TCTGATCATTGCAACCCCTG |
| | hRPL19-DRIP-R | ACCTCTAATCCCAGCTACTCAG |
| | hTUBA1A-DRIP-NC-F | TGTTTCTGGCTTCTATGGCG |
| | hTUBA1A-DRIP-NC-R | AGCCTACAGTTCCGCAATG |
| | hRPL28-DRIP-F | CCCCAGCCTCATCTTTTAATG |
| | hRPL28-DRIP-R | ATTAGGGAAAAGGAGCAAGGG |
